# Genetic analysis of triplicated genes affecting sex-specific skeletal deficits in Down syndrome model mice

**DOI:** 10.1101/2025.01.31.635850

**Authors:** Kourtney Sloan, Kristina M. Piner, Pathum Randunu Nawarathna Kandedura Arachchige, Charles R. Goodlett, Yann Herault, Gayla R. Olbricht, Joseph M. Wallace, Randall J. Roper

## Abstract

Down syndrome (DS) is caused by the triplication of human chromosome 21 (Hsa21), resulting in skeletal insufficiency and altered bone development. DS mouse models recapitulate these deficits, including sexual dimorphism in long bone alterations. Historically, Ts65Dn mice provided much of the insight behind DS-related skeletal deficits with ∼100 trisomic orthologous genes, but there are concerns about genetic fidelity in this model due to included triplication of genes not homologous to Hsa21. A new DS mouse model, Ts66Yah, subtracted the non-Hsa21 homologous trisomic genes from Ts65Dn but has not been evaluated for long bone deficits. Comparing skeletal phenotypes between these models can indicate the contributions of non-Hsa21 trisomic genes and whether the Ts66Yah mouse is relevant as a model for DS-associated skeletal deficits. After assessing individual densitometric, morphometric, and mechanical variables in male and female Ts66Yah femurs at similar ages to when skeletal deficits had been observed in Ts65Dn mice, structural phenotypes were directly compared to those of Ts65Dn mice using a novel multivariate principal components analysis (PCA) method to generate composite scores. Overall, structural and mechanical bone phenotypes of the femur appear milder in Ts66Yah compared to Ts65Dn mice. The appearance of developmental trabecular microarchitecture deficits, but not other abnormalities, were evident earlier in Ts65Dn than Ts66Yah mice. Dyrk1a, a gene triplicated in both models, affected skeletal structure differently in each model, likely through differing gene interactions. The novel component score analysis incorporating PCA detected subclinical phenotypes lost in individual analyses, which could be advantageous when determining overall skeletal deficits.

**Article summary:** Mouse models are essential for understanding mechanisms behind human conditions, such as Down syndrome (DS). This study evaluates long bone morphology and strength at key timepoints of development in a new DS mouse model and introduces a novel method derived from principal component analysis (PCA) to compare phenotypes between two different DS models. While both models generally exhibit similar sex-specific deficits, the window of efficacy for genetic and pharmacological intervention on a therapeutic target varies. This illustrates the importance of validating DS phenotypes and mechanisms in multiple mouse models. The method developed could be used in broader scientific applications.

## Introduction

Down syndrome (DS), occurring in about 1 in 800 live births (DE GRAAF *et al*. 2017), caused by Trisomy 21 (Ts21), is the most common aneuploidy, resulting in an extra full or partial human chromosome 21 (Hsa21). Ts21 is known to cause cognitive deficits and affect the musculoskeletal system, resulting in a heightened risk of osteoporosis (ANTONARAKIS *et al*. 2020). The genetic mechanisms, including which and how many genes are involved, are largely unknown and may vary between phenotypes to result in a developmental and/or progeroid trajectory (MOYER *et al*. 2021). Within the Ts21 skeleton, there are developmental differences in addition to advanced aging (THOMAS AND ROPER 2021). These developmental differences include shortened long bones and a compressed period of bone growth, such that people with DS reach peak bone mass 5-10 years earlier than the general population (COSTA *et al*. 2018). This results in low bone mineral density (BMD) at typical skeletal maturity, which is further exacerbated by BMD that decreases more with age in individuals with DS than those without (CARFI *et al*. 2017; GARCIA HOYOS *et al*. 2019; TANG *et al*. 2019). The contribution of developmental differences and premature aging to DS skeletal deficits appears to vary by sex. Women with DS maintain similar BMD at the femoral neck as women without DS until their 40s, whereas men with DS only maintain similar BMD as men without DS until their 30s (CARFI *et al*. 2017). These sex differences are also dependent on bone compartment. Men with DS over the age of 18 have lower integral (cortical + trabecular) BMD, trabecular BMD, cortical BMD, and cortical thickness in their femurs compared to men without DS, whereas women with DS only had lowered cortical variables compared to women without DS (GARCIA HOYOS *et al*. 2019). When directly comparing male and female individuals with DS, males over the age of 15 had lower integral and trabecular BMD than females but no difference in cortical BMD (COSTA *et al*. 2021). These factors of age and sex contributed to increased fractures in individuals with DS (KRIEG *et al*. 2024).

To investigate Ts21 gene-phenotype relationships, multiple DS mouse models with varying content and transmission of triplicated genes are utilized, reflecting differences between mouse and human genetics. The first viable mouse model of DS was the Ts(17^16^)65Dn (Ts65Dn), which has a freely segregating minichromosome that is trisomic for ∼50% of Hsa21 orthologous genes found on mouse chromosome (Mmu)16 from *Mir155* to *Zbtb21* (STURGEON AND GARDINER 2011; DUCHON *et al*. 2022). Male Ts65Dn mice have been studied the most and show many DS-like phenotypic similarities to humans with DS, including in the skeletal system. The cartilage template and primary ossification center of Ts65Dn embryos indicate deficits arise from abnormalities in mineralization rather than the building of the cartilage template (BLAZEK *et al*. 2015b). In postnatal studies, male Ts65Dn mice have cortical deficits as early as postnatal day (P)12 whereas trabecular deficits do not appear until and persist after P30 (LACOMBE *et al*. 2024). Both trabecular and cortical deficits have been reported in male Ts65Dn mice at P36, P42 (6 weeks), 8 weeks, 12 weeks (3 months), 16 weeks, and 24 months (BLAZEK *et al*. 2011; FOWLER *et al*. 2012; WILLIAMS *et al*. 2018; SHERMAN *et al*. 2022; LACOMBE *et al*. 2024). Less is known about female Ts65Dn mice, but current findings suggest trabecular and cortical deficits are evident at P30, subsequently normalize and then are found again at P42; no later ages have been investigated (THOMAS *et al*. 2021; LACOMBE *et al*. 2024).

The Hsa21 gene *dual specificity tyrosine phosphorylation-regulated kinase 1A* (*DYRK1A*) has been linked to altered skeletal development, with a singular effect on trabecular skeletal deficits in male mice, and interactive effects on cortical bone (ARRON *et al*. 2006; LEE *et al*. 2009; EGUSA *et al*. 2011; SLOAN *et al*. 2023). These studies suggested that reducing *Dyrk1a* copy number in male Ts65Dn mice could rescue some skeletal deficits. The therapeutic effects of reduction of *Dyrk1a* copy number are developmentally regulated: normalization of *Dyrk1a* copy number through breeding with a *Dyrk1a* haploinsufficiency model (*Dyrk1a*^tm1Mla^ or *Dyrk1a*^+/-^) does not rescue skeletal deficits in embryonic Ts65Dn mice, but does improve or rescue some skeletal deficits postnatally (P36-P42) (BLAZEK *et al*. 2015a; BLAZEK *et al*. 2015b; LACOMBE *et al*. 2024). The improvement of structural phenotypes in long bones is limited to male Ts65Dn mice only; female mice are not affected by *Dyrk1a* copy number normalization at the same age (LACOMBE *et al*. 2024). Additionally, genetic reduction of *DYRK1A* in euploid individuals leads to skeletal deficits associated with DYRK1A syndrome, and similar effects have been observed in euploid mouse models with reduced *Dyrk1a* levels (OTTE AND ROPER 2024).

Reducing DYRK1A kinase activity is also hypothesized to improve skeletal abnormalities in Ts65Dn mice. Epigallocatechin-3-gallate (EGCG), the main catechin found in green tea extract, and Silmitasertib, also known as CX-4945, both have an affinity for the ATP-binding site of DYRK1A that may reduce DYRK1A activity. Both have been shown to produce minimal or no improvement in, or even worsen, cortical bone of Ts65Dn mice depending on dosage, timing of treatment based on DYRK1A expression, or length of treatment time (GOODLETT *et al*. 2020; JAMAL *et al*. 2022; LACOMBE *et al*. 2024). Leucettinib-21 is a second-generation of Leucettamine B analog that also binds to the DYRK1A ATP-binding pocket (LINDBERG *et al*. 2023) but is more potent for DYRK1A (IC_50_ = 2.4 nM) than EGCG (491 nM) or CX-4945 (6.8 nM) (KIM *et al*. 2016; LINDBERG *et al*. 2023). Leucettinib-21 treatment showed improved memory based on the novel object recognition test in adult male Ts65Dn mice (LINDBERG *et al*. 2023), and partially rescued congenital heart defects in Dp1Tyb embryos (LANA-ELOLA *et al*. 2024). However, effects of this DYRK1A inhibitor on skeletal phenotypes has not been tested.

In addition to Hsa21 homologs, Ts65Dn mice also contain a triplication of ∼60 trisomic genes (∼35 protein-coding) located on the Mmu17 centromeric region from *Scaf8* to *Pde10a* that are orthologous to Hsa6, rather than Hsa21 (DUCHON *et al*. 2011; REINHOLDT *et al*. 2011). The contribution and impact of these non-Hsa21 orthologous trisomic genes in observed Ts65Dn phenotypes are unknown. The Ts(17^16^)66Yah (Ts66Yah) mouse model was developed using CRISPR/Cas9 to remove the trisomic Mmu17 centromeric region from the Ts65Dn minichromosome and has been characterized for craniofacial, behavior, morphological and molecular brain, and growth phenotypes from late embryos to almost middle age (DUCHON *et al*. 2022; GUEDJ *et al*. 2023; ADAMS *et al*. 2024; LANZILLOTTA *et al*. 2024; EMILI *et al*. 2025). To date, Ts66Yah mice mostly exhibit DS-associated phenotypes, with notable exceptions of hippocampal dendrite pathology and neurogenesis alterations. However, Ts66Yah phenotypes are generally less severe than those of Ts65Dn mice, indicating at least a minor role of the triplicated non-Hsa21 homologs. The objectives of this study were first to characterize Ts66Yah femoral morphology and strength during postnatal development at critical periods of time identified in Ts65Dn mice, then to determine whether these phenotypes differ between Ts65Dn and Ts66Yah mice. This comparison allows us to determine whether non-Hsa21 orthologous, trisomic genes contribute to long bone deficits in Ts65Dn mice and to assess the validity of Ts66Yah mice as a model for DS-related skeletal deficits. The assessment of the Ts66Yah mouse model was further extended by characterizing the effects of trisomic *Dyrk1a,* an Hsa21 gene implicated in aspects of skeletal and other DS-related phenotypes, in Ts66Yah mice through genetic and pharmacological means.

## Methods

### Animals

Ts(17^16^)66Yah (Ts66Yah) mice were obtained from the French National Centre for Scientific Research (CNRS) courtesy of Dr. Yann Herault and have been described previously (DUCHON *et al*. 2022). The F1 generation between the male founder mouse and C57BL/6NCrl female mice were backcrossed at least 10 times to F1 hybrids of the C57BL/6NCrl line and C3H/HeN line congenic for the BALB/c allele at the *Pde6b* locus (B6C3BF1) before three female Ts66Yah mice were transferred to the Indiana University (IU) Indianapolis Science Animal Resource Center.

The IU Indianapolis Ts66Yah colony was maintained by breeding female Ts66Yah mice to male B6C3F1 mice obtained through crossing C57BL/6J (Jackson Laboratory strain #000664) and C3H/HeJ mice that carry retinal degeneration allele *Pde6b^rd1^* (Jackson Laboratory strain #000659). For the reduction of *Dyrk1a* copy number experiment, female Ts66Yah mice were bred to male *Dyrk1a* heterozygous mutant (*Dyrk1a*^tm1Mla^ or *Dyrk1a*^+/-^) mice (FOTAKI *et al*. 2002) to yield four potential offspring genotypes: euploid animals with homozygous wildtype *Dyrk1a* (control euploid [euploid]), euploid animals with heterozygous mutant *Dyrk1a* (euploid with one functional copy of *Dyrk1a* [euploid,*Dyrk1a*^+/-^]), trisomic (Ts66Yah) animals with homozygous wildtype *Dyrk1a* (trisomic animals with three functional copies of *Dyrk1a* [Ts66Yah]), and trisomic (Ts66Yah) animals with heterozygous mutant *Dyrk1a* (trisomic animals with two functional copies of *Dyrk1a* [Ts66Yah,*Dyrk1a*^+/+/-^]). The *Dyrk1a*^tm1Mla^ mice were maintained by breeding with a B6C3F1 mouse, like the Ts66Yah mice, in our colony.

Ts66Yah mice were genotyped by PCR to amplify the minichromosome breakpoint between the mouse chromosome (Mmu) 17 centromere and Mmu16 genes (*Mir155* to *Zbtb21*) as previously described (DUCHON *et al*. 2022). *Dyrk1a* heterozygous mutants were genotyped by PCR using two primers to amplify a segment of the neomycin cassette (Dyrk1a Neo P1) and exon 6/intron region 6-7 of *Dyrk1a* (Dyrk1a P3) based on genome assembly GRCm39, gene ENSMUSG00000022897, transcript ENSMUST0000119878.8 (FOTAKI *et al*. 2002).

All animals are weaned around postnatal day (P) 21 and housed in cages of 1-5 mice of mixed genotype with a nestlet for enrichment and access to food and water *ad libitum*. Two male Ts66Yah animals out of 183 total Ts66Yah animals allocated were removed from the study due to the presence of hydrocephalus. All animal use and protocols were approved by the Institutional Animal Care & Use Committee (IACUC) at the IU Indianapolis School of Science (SC338R) and adhered to the requirements in the NIH Guide for the Care and Use of Laboratory Animals. The Animal Research: Reporting of *In Vivo* Experiments (ARRIVE) guidelines were also followed.

### Leucettinib-21 (L21) treatment

Leucettinib*-*21 (L21) solution was made weekly by suspending 2mg of L21 in 1mL of the vehicle, 0.5% weight by volume carboxymethyl cellulose (CMC). The 2mg/mL L21 solution was diluted to 0.05mg/mL L21 working solution, both of which were stored at -20°C. Using the 0.05mg/mL L21 solution, each animal was weighed and treated daily 10mL/kg via oral gavage from P21 through P35 for a final dosage of 0.5mg/kg/day. Animals were euthanized after recording body weight on P36. All animals of the first cohort were given the vehicle treatment, then all animals in the next cohort were given L21 treatment. This alternation occurred throughout the study. A cohort was defined as 3 to 12 mice from 1 to 3 litters born no more than 3 days apart. A total of 17 cohorts consisting of 21 litters were treated: 9 cohorts (10 litters) given vehicle and 8 cohorts (11 litters) given L21.

### Bone storage and micro-computed tomography (µCT) analysis

Right femurs were wrapped in 1x phosphate buffered saline (PBS)-soaked gauze and stored at -20°C until scanning. Six-, nine-, and sixteen-week-old Ts66Yah femurs were wrapped in parafilm to maintain hydration and single-scanned using a SkyScan 1172 µCT system (Bruker, Kontich, Belgium) with the following parameters: 60kV, 167uA, 885ms, 10-micron voxel size, Al 0.5mm filter, 0.7□ rotation step and frame averaging of two. Two hydroxyapatite phantoms (0.25 and 0.75g/cm^3^ CaHA) were used to calibrate bone mineral density for each scanning session.

Due to irreparable damage to the SkyScan 1172 µCT system, femurs from P36 mice were wrapped in parafilm and group-scanned (KOHLER *et al*. 2021) using a SkyScan 1272 µCT system (Bruker, Kontich, Belgium) with the following parameters: 70kV, 142uA, 1265ms, 10-micron voxel size, Al 0.5mm filter, 0.7□ rotation step and frame averaging of two. Two hydroxyapatite phantoms (0.3 and 1.25g/cm^3^ CaHA) were used to calibrate bone mineral density for each scanning session. No unstandardized comparisons were made between scans from different µCT systems.

All scans were reconstructed using NRecon with the following settings: ring artifact reduction 5, beam-hardening correction 20%, dynamic image (attenuation coefficient) boundaries 0.00 and 0.11. Reconstructed scans were rotated so the anterior surface of the distal growth plate faces right in the transaxial view for all bones. Using CTAnalyzer (CTAn), the trabecular region of interest (ROI) was defined as a 1mm section extending proximally from the end of the distal growth plate and isolated from the cortical bone. The cortical ROI was defined as a 1mm (6-, 9-, and 16-week-old mice) or 0.65mm (P36 mice) section extending distally from the beginning of the third trochanter. Trabecular bone variables, including bone mineral density (BMD), bone volume fraction (BV/TV), trabecular thickness (Tb.Th), trabecular separation (Tb.Sp), and trabecular number (Tb.N), were calculated using internal CTAn functions. Cortical bone variables, including total cross-sectional area (Tt.Ar), marrow area (Ma.Ar), cortical bone area (Ct.Ar), cortical bone area fraction (Ct.Ar/Tt.Ar), cortical thickness (Ct.Th), periosteal perimeter (Ps.Pm), endocortical perimeter (Ec.Pm), maximum and minimum moment of inertia (I_max_ and I_min_) for P36 bones, and cortical tissue mineral density (Ct.TMD), were calculated using a custom MATLAB code (BERMAN *et al*. 2015; STRINGER *et al*. 2017a). The American Society for Bone and Mineral Research (ASBMR) µCT guidelines were followed (BOUXSEIN *et al*. 2010).

### Mechanical testing

Mechanical properties were determined as described previously (THOMAS *et al*. 2020). Briefly, 3-point femur bend was performed after µCT analysis of 6-, 9-, and 16-week-old right femurs using a TA ElectroForce 5500 testing machine while bones were hydrated using PBS (Eden Prairie, MN, USA). Right femurs were tested using a 5-, 6-, or 7mm support span (6, 9, and 16 weeks, respectively) in the anterior-posterior direction with the posterior surface in compression. A 10lb load was utilized for 6- and 9-week-old femurs, and a 50lb load was utilized for 16-week-old femurs. All bones were preloaded (0.2-0.4 N) to establish contact with the loading point located at the midshaft, then testing occurred at a rate of 0.025mm/sec to failure. The 0.2% offset method was utilized on the slope of the linear portion of the stress-strain curve to find the yield point. The ultimate point was determined as the maximum force reached, while the failure point was determined as where the bone broke. Two 6-week-old female euploid femurs were excluded from analysis due to not reaching a failure point before maximum displacement at 3.5mm. The following whole bone (extrinsic) variables were reported from the force-displacement curve: yield and ultimate force, displacement and work to yield, postyield displacement and work, total displacement and work, and stiffness. Cortical geometry from µCT analysis was utilized to normalize the stress-strain curve from the force-displacement curve. The following tissue-estimate (intrinsic) properties were reported from the stress-strain curve: yield and ultimate stress, strain to yield, total strain, modulus, resilience, and toughness.

### Comparison between Ts65Dn and Ts66Yah mice

The trabecular and cortical bone variables of each animal were standardized using the mean and standard deviation of all within each mouse model (Equation 1). This was performed separately for each sex and age. Six-week-old euploid and Ts65Dn data came from a subset of data presented in (THOMAS *et al*. 2021), only including euploid littermates and Ts65Dn mice lacking OSX-cre, whereas 16-week-old euploid and Ts65Dn trabecular data came from (BLAZEK *et al*. 2011). The P36 euploid and Ts65Dn data came from (LACOMBE *et al*. 2024).

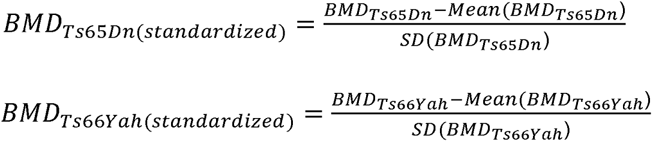

*Equation 1:* Example of how each variable was standardized within mouse model separately for each sex and age. The BMD variable is shown for illustration.

Principal components analysis (PCA) was performed on standardized µCT data of Ts65Dn mice, Ts66Yah mice, and each of their respective euploid littermate controls, separately for each sex, age, and bone compartment. Other PCAs were performed on standardized µCT data of trisomic Ts65Dn mice, Ts66Yah mice, and *Dyrk1a* copy number reduction mice of both models, separately for each sex and bone compartment. The principal component 1 (PC1) loading of each variable was multiplied by the animal’s standardized value. Then, the sum of these transformations was calculated to produce a single value to represent the animal’s trabecular or cortical bone, which will be referred to as the trabecular or cortical composite score (Equation 2, 3).

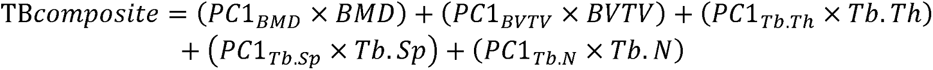

*Equation 2:* Calculation performed for each animal to find the trabecular composite score using standardized trabecular variables. For illustration, *PC*1*_BMD_* represents the loading on PC1 for the BMD variable.

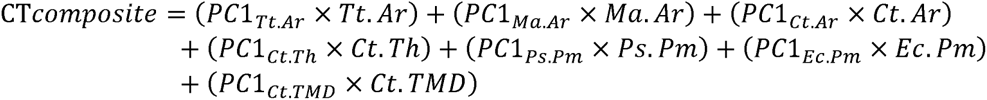

*Equation 3:* Calculation performed for each animal to find the cortical composite score using standardized cortical variables. For illustration, *PC*1*_Tt.Ar_* represents the loading on PC1 for the Tt.Ar variable.

### Statistical analysis

Chi-squared goodness of fit tests were performed using Microsoft Excel to determine differences between expected Mendelian and observed genotype ratios for total counts of pre-and post-weaned offspring of Ts66Yah × B6C3F1 and Ts66Yah × *Dyrk1a*^+/-^ breeding schemes. Correlation and regression analyses of body weight and femur length at P36 in euploid and Ts66Yah mice treated with vehicle and L21 were performed in Microsoft Excel separately for each sex using group averages and sample size. Significance of these correlations were determined by two-tailed Student’s t-distribution (□ = 0.05). Remaining statistical analyses were performed using IBM SPSS Statistics (v29.0.1.0). The Shapiro-Wilk test (□ = 0.05) was used to assess normality. In cases where the data were non-normal, data were log-transformed and assessed again. The assumption of equal variances was assessed using Levene’s test (□ = 0.05).

For most 6-, 9-, and 16-week-old Ts66Yah data, three-way ANOVAs were performed with genotype, sex, and age as between-subject factors along with their interactions. Due to differences in 3-point bend set-up between each age, mechanical data were assessed separately for each age by two-way ANOVA with genotype and sex as between-subject factors along with their interaction. For P36 L21 treatment change in weight, femur length, and µCT data, three-way ANOVAs with genotype, sex, and treatment as between-subject factors and their interactions were performed. For the P36 germline reduction of *Dyrk1a* experiments, three-way ANOVAs were performed using genotype (euploid vs. trisomic), sex, and *Dyrk1a* reduction (wildtype vs. mutant) as between-subject factors. A significant finding was defined as p < 0.05. In cases where both Levene’s test and ANOVA p values were significant, the data were re-assessed by Welch’s *F* statistic. In rare instances when data were not significant during re-assessment, the variable was not reported as significant. Pairwise comparisons with Sidak correction were performed in cases of significant interactions.

Additionally, for the P36 L21 treatment experiment, a repeated measures ANOVA was performed for body weight using genotype, sex, and treatment as between-subject factors and postnatal day as the within-subject factor. Sphericity was assessed using Mauchly’s test, and the Greenhouse-Geisser test was utilized to detect significant effects and interactions due to a significant Mauchly’s test. Pairwise comparisons with Sidak correction were performed for significant interactions.

Using R (version 4.3.0), trabecular and cortical composite scores generated from the PCAs were utilized as responses in two-way ANOVAs with either Ts66Yah genotype (euploid vs. trisomic) and sex (male vs. female), mouse model (Ts65Dn vs. Ts66Yah) and genotype (euploid vs. trisomic), or mouse model (Ts65Dn vs. Ts66Yah) and *Dyrk1a* copy number (2 vs. 3) as between subject main effect factors, along with their interaction. When the interaction was significant, contrast analyses were performed with Bonferroni correction to test for differences between genotypes or *Dyrk1a* copy number within sex or mouse model as applicable, enabling an understanding of effect differences between models. Type III Sum of Squares were utilized for testing each factor due to differing numbers of mice per group.

## Results

### Higher rate of trisomy in Ts66Yah animals

The Ts66Yah x B6C3F1 breeding scheme yielded 114 litters with an average litter size of 4-5 mice in our laboratory (Supplemental Table 1). The transmission of trisomy was 44% between P6 and P10 and at weaning, similar to that reported in another lab for the first year of breeding on a similar background (44%) but higher than their 6-year total (33%) (DUCHON *et al*. 2022). Compared to Ts65Dn mice, this rate is also higher than previously reported transmission at weaning (∼34%) (MOORE 2006; ROPER *et al*. 2006). The transmission rate at weaning was similar in both sexes when averaged across the years, but there were varying rates throughout the colony’s time. When Ts66Yah mice were bred with *Dyrk1a*^+/-^ mice, litter sizes were slightly higher (average 5-6 mice) across 26 litters (Supplemental Table 2). When partitioned into the four possible genotypes, fewer euploid,*Dyrk1a*^+/-^ and Ts66Yah mice were found between P6 and P10 whereas more Ts66Yah,*Dyrk1a*^+/+/-^ mice were present than expected. This was the case in both male and female mice at weaning, indicating abnormal dosage of *Dyrk1a* may result in more prenatal and perinatal death.

### Structural phenotypes of the femur vary by age and sex in Ts66Yah mice

Appendicular structural phenotypes of Ts66Yah mice were evaluated after sexual maturity during skeletal accrual (6 and 9 weeks) and around skeletal maturity (16 weeks) as done previously for Ts65Dn and other DS mouse models. There were no three-way interactions between genotype, sex, and age, but both body weight and femur length had significant genotype and sex interactions (body weight: *p* = 0.019; femur length: *p =* 0.002). Pairwise comparisons between genotypes within each sex and age showed no significant differences in body weight (Supplemental Figure 1A); however, male Ts66Yah mice had significantly shorter femurs at 9 and 16 weeks compared to male euploid mice (Supplemental Figure 1B). Femur length also displayed a significant sex and age interaction (*p* = 0.016), with pairwise comparisons between each age within each genotype and sex showing significant increases between ages in femur length for all four groups. The similar body weight in Ts66Yah mice compared to euploid mice is unlike the reduced weight of Ts65Dn mice (ROPER *et al*. 2006; BLAZEK *et al*. 2015a; WILLIAMS *et al*. 2018), but another DS mouse model also lacked body weight differences past 4 weeks of age (LANA-ELOLA *et al*. 2021).

For trabecular bone structure of 6, 9, and 16-week-old euploid and Ts66Yah mice (see representative images in Figure 1A), no significant three-way interactions between genotype, sex, and age were detected, but bone mineral density (BMD), bone volume fraction (BV/TV) and trabecular thickness (Tb.Th) had significant genotype and sex interactions (BMD: *p* = 0.020; BV/TV: *p* = 0.022; Tb.Th: *p* = 0.012). Pairwise comparisons between genotypes within each sex and age showed significantly lower BMD and BV/TV in male Ts66Yah mice compared to male euploid mice at 6 and 16 weeks and significantly lower Tb.Th in male Ts66Yah mice at 6 and 9 weeks (Figure 1B-D). No significant differences were found between genotypes of female mice in any trabecular variable. A significant age and sex interaction (*p* = 0.011) was found in Tb.Th with pairwise comparisons between each age within each genotype and sex showing no significant differences with age in either genotype of male mice, increased Tb.Th at 9 and 16 weeks compared to 6 weeks in female euploid mice and increased Tb.Th at 16 weeks compared to 6 weeks in Ts66Yah mice. Trabecular separation (Tb.Sp) and number (Tb.N) had no significant interactions in the 3-way ANOVA with genotype, sex, and age as main effects (Figure 1E-F).

**Figure 1:**
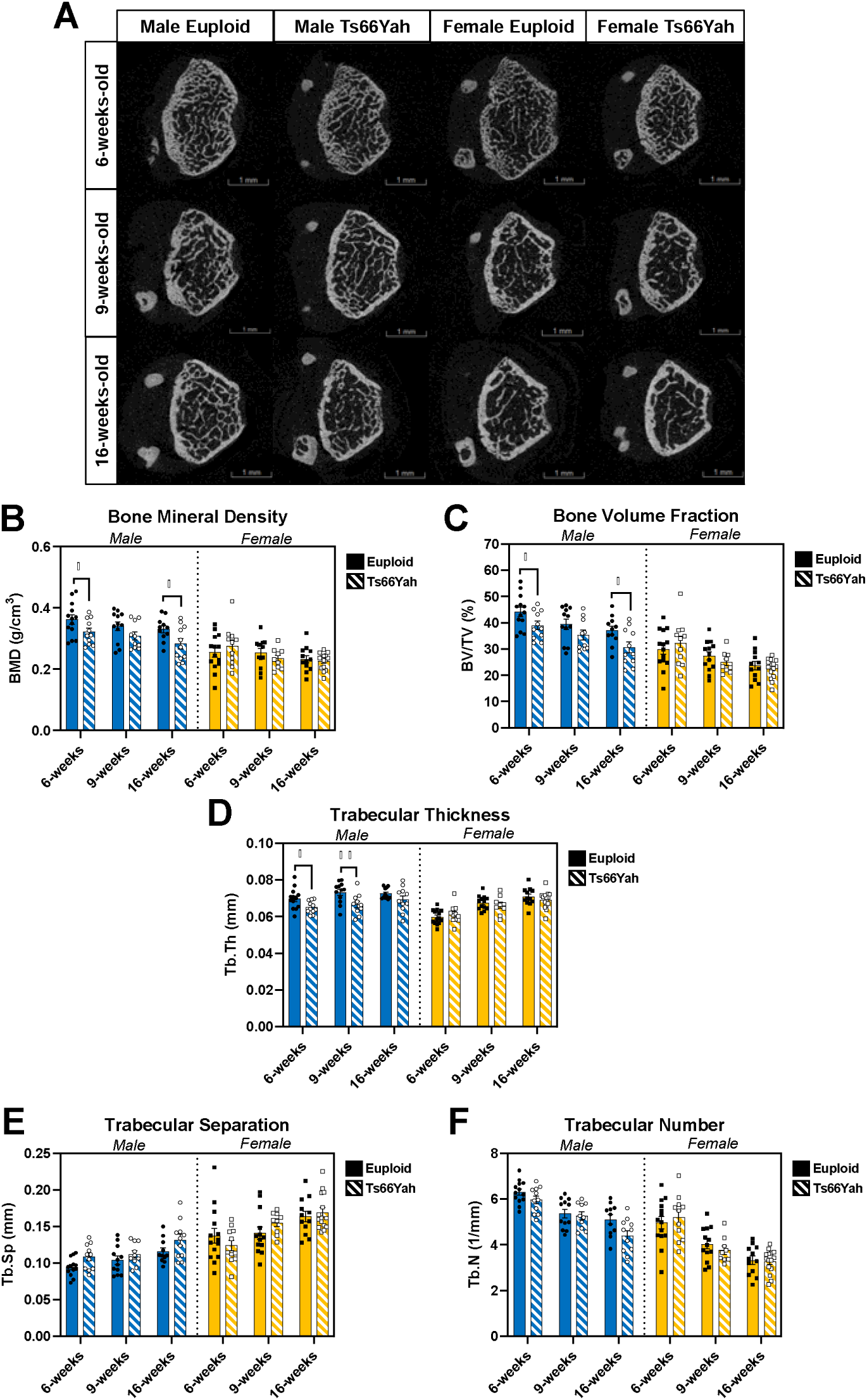
Trabecular bone variables for 6-, 9-, and 16-week-old Ts66Yah mice. **A)** Representative images of trabecular bone taken halfway through the 1mm trabecular region as determined by finding the animal with the closest average distance away from the mean of each trabecular variable. **B-F)** Data are mean ± SEM. Asterisks indicate a significant difference between groups in pairwise comparisons with Sidak correction. * *p* < 0.05, ** *p* < 0.01. **D)** Pairwise comparisons between ages within genotype and sex for Tb.Th: no significant differences in male euploid or Ts66Yah mice, significantly increased at 9 and 16 weeks compared to 6 weeks in female euploid mice, and significantly increased at 16 weeks compared to 6 weeks in female Ts66Yah mice. 6 weeks: male euploid (n = 13), male Ts66Yah (n = 11), female euploid (n = 14), female Ts66Yah (n = 11); 9 weeks: male euploid (n = 12), male Ts66Yah (n = 10), female euploid (n = 13), female Ts66Yah (n = 9); 16 weeks: male euploid (n = 11), male Ts66Yah (n = 11), female euploid (n = 12), female Ts66Yah (n = 15).

In the cortical bone compartment of Ts66Yah x B6C3F1 offspring (see average cross-section representations in Figure 2A), three-way interactions between genotype, sex, and age were detected for marrow area (Ma.Ar; *p* = 0.006), cortical bone area fraction (Ct.Ar/Tt.Ar; *p* = 0.014), and endocortical perimeter (Ec.Pm; *p* = 0.004). Pairwise comparisons between genotypes within each sex and age showed significantly decreased Ma.Ar and Ec.Pm in male Ts66Yah mice compared to male euploid mice at 9 and 16 weeks and in female Ts66Yah mice at 6 weeks (Figure 2E,H). However, Ct.Ar/Tt.Ar was significantly increased in male Ts66Yah mice at 9 and 16 weeks compared to male euploid mice and in female Ts66Yah mice at 6 and 9 weeks (Figure 2D).

**Figure 2:**
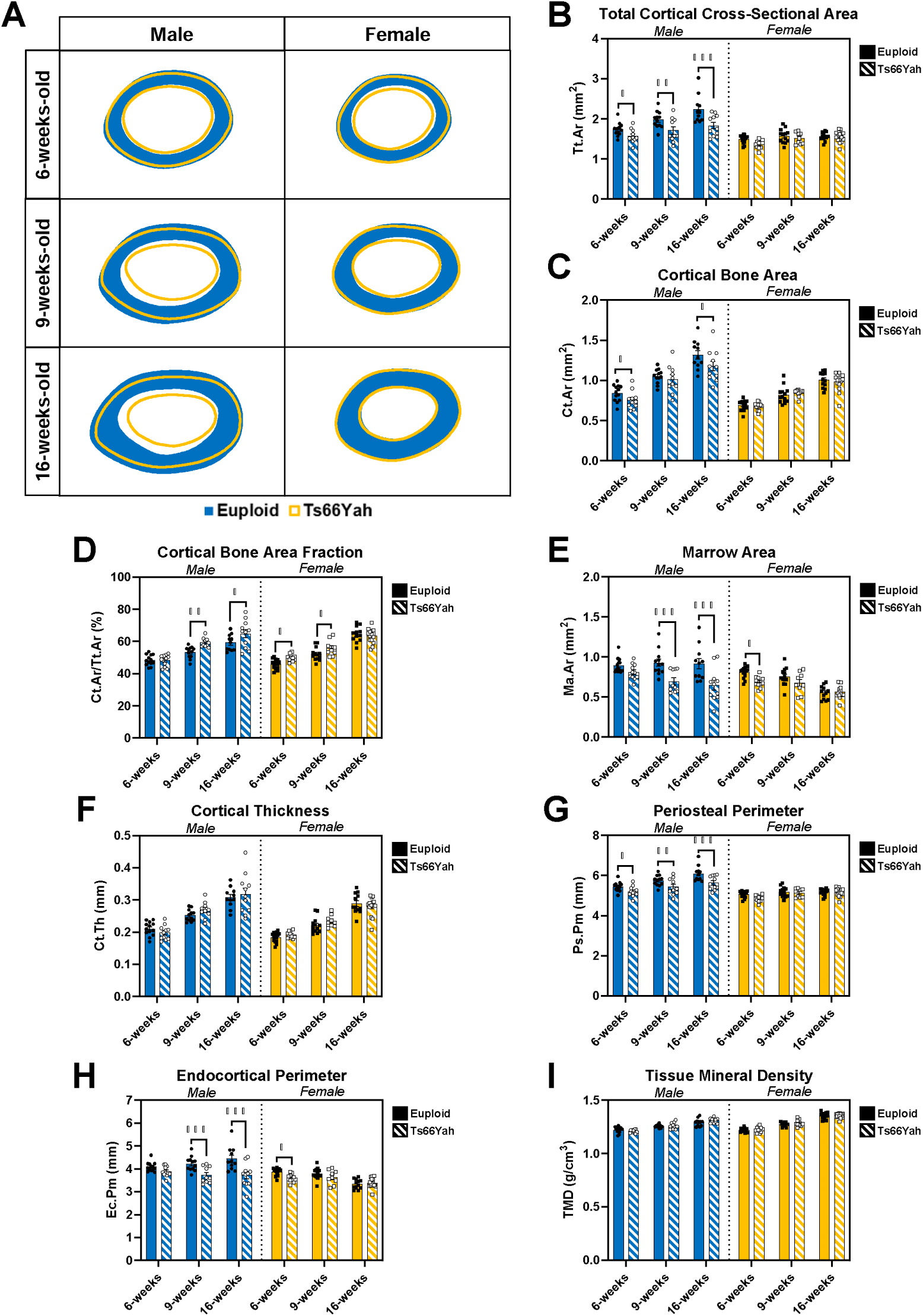
Cortical bone analysis of 6-, 9-, and 16-week-old Ts66Yah mice. **A)** Representations of the average cortical cross-section for each group made by calculating the centroid of each cortical section and finding the distance from the centroid to the endocortical and periosteal surfaces every 0.5 degrees. The average is found for each animal, then the average of the group is found to make the radar graph. **B-I)** Data are mean ± SEM. Asterisks indicate a significant difference between groups in pairwise comparisons with Sidak correction. **B)** Pairwise comparisons between ages within genotype and sex for Tt.Ar: significantly increased between each age in male euploid mice and no significant differences in male Ts66Yah mice or either genotype of female mice. **D)** Pairwise comparisons between ages within genotype and sex for Ct.Ar/Tt.Ar: significantly increased between each age in male euploid mice and both female groups and significantly increased at 9 and 16 compared to 6 weeks in male Ts66Yah mice. **E)** Pairwise comparisons between ages within genotype and sex for Ma.Ar: no significant differences in male euploid mice, significantly decreased at 16 compared to 6 weeks in both sexes of Ts66Yah mice, and significantly decreased at 16 compared to 6 and 9 weeks in female euploid mice. **G)** Pairwise comparisons between ages within genotype and sex for Ps.Pm: significantly increased between each age in male euploid mice, significantly increased at 16 compared to 6 weeks in both sexes of Ts66Yah mice, and no significant differences in female euploid. **H)** Pairwise comparisons between ages within genotype and sex for Ec.Pm: significantly increased at 16 compared to 6 weeks in male euploid mice, no significant differences in either sex of Ts66Yah mice, and significantly decreased at 16 compared to 6 and 9 weeks in female euploid mice. **I)** Pairwise comparisons between ages within genotype and sex for TMD: significantly increased between each age in all four groups. * *p* < 0.05, ** *p* < 0.01, *** *p* < 0.001. 6 weeks: male euploid (n = 13), male Ts66Yah (n = 11), female euploid (n = 14), female Ts66Yah (n = 11); 9 weeks: male euploid (n = 12), male Ts66Yah (n = 10), female euploid (n = 13), female Ts66Yah (n = 9); 16 weeks: male euploid (n = 11), male Ts66Yah (n = 11), female euploid (n = 12), female Ts66Yah (n = 15).

Pairwise comparisons between ages within each genotype and sex showed no significant differences in Ma.Ar in male euploid mice, while Ma.Ar significantly decreased from 6 to 16 weeks in male and female Ts66Yah mice, and female euploid mice saw significantly smaller Ma.Ar at 16 weeks than 6 and 9 weeks (Figure 2E). Ct.Ar/Tt.Ar significantly increased between each age in male euploid and both female groups and only significantly increased between 6 and 9 weeks in male Ts66Yah mice (Figure 2D). Ec.Pm was significantly greater at 16 weeks than 6 weeks in male euploid mice, but no differences were detected in either sex of Ts66Yah mice, and female euploid showed lower Ec.Pm at 16 weeks than 6 or 9 weeks (Figure 2H).

Two-way interactions between genotype and sex were detected for total cortical cross-sectional area (Tt.Ar; Figure 2B, *p* = 0.001), cortical bone area (Ct.Ar; Figure 2C, *p* = 0.036), and periosteal perimeter (Ps.Pm; Figure 2G, *p* = 0.006). Pairwise comparisons between genotypes within each sex and age showed male Ts66Yah mice had significantly decreased Tt.Ar and Ps.Pm at all three ages and significantly decreased Ct.Ar at 6 and 16 weeks compared to male euploid mice, whereas female mice had no significant differences.

Additional two-way interactions between sex and age were detected for Tt.Ar (Figure 2B, *p* = 0.005), Ps.Pm (Figure 2G, *p* = 0.004), and tissue mineral density (Ct.TMD; Figure 2I, *p* < 0.001). Pairwise comparisons between ages within each genotype and sex showed only male euploid mice significantly increased Tt.Ar at each age while male Ts66Yah mice and both female groups did not show any significant differences. Male euploid mice also significantly increased Ps.Pm at each age, while both male and female Ts66Yah mice had significantly greater Ps.Pm at 16 weeks than at 6 weeks, and female euploid mice did not significantly differ between ages. Ct.TMD significantly increased between each age in all four groups. Only cortical thickness (Ct.Th) did not display any significant interactions in a three-way ANOVA with genotype, sex, and age as main effects (Figure 2F).

To understand if Ts66Yah femurs above were more susceptible to fracture, their structural and tissue mechanics were evaluated separately at each age by the three-point bend test. For 6-week-old structural properties, only yield force (*p* = 0.047) and ultimate force (*p* = 0.023) had a significant interaction between genotype and sex. Pairwise comparisons between genotypes within sex showed male, but not female, Ts66Yah femurs had a significantly lower yield force and ultimate force compared to sex-matched euploid femurs (Figure 3A; Table 1). For estimated tissue mechanics at 6-weeks, a significant interaction between genotype and sex was only found in yield stress (*p* = 0.050). Pairwise comparisons between genotypes within sex showed female, but not male, Ts66Yah femurs had a significantly higher yield stress than sex-matched euploid femurs (Figure 3B; Table 2).

**Figure 3:**
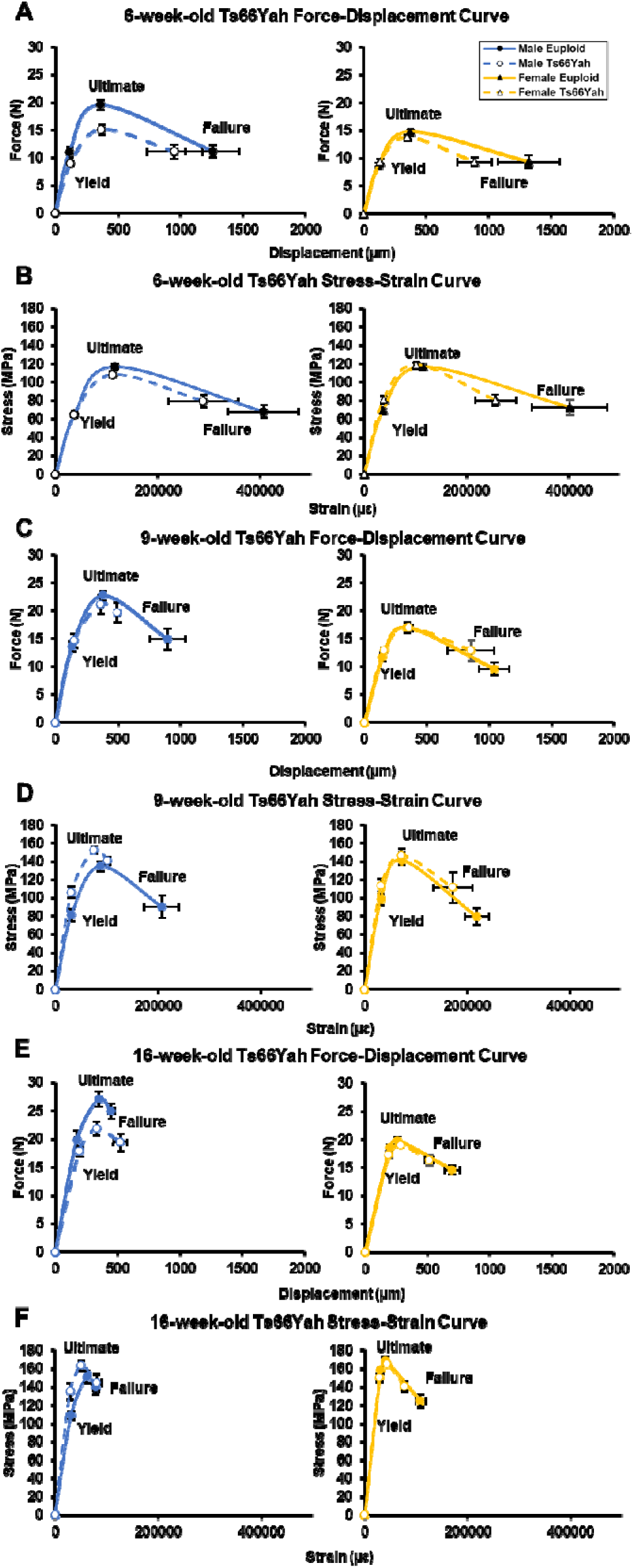
Mechanical curves generated from 3-point femur bend of 6-, 9- and 16-week-old Ts66Yah mice. Data are mean ± SEM. 6 weeks: male euploid (n = 12), male Ts66Yah (n = 11), female euploid (n = 11), female Ts66Yah (n = 11); 9 weeks: male euploid (n = 12), male Ts66Yah (n = 10), female euploid (n = 13), female Ts66Yah (n = 9); 16 weeks: male euploid (n = 11), male Ts66Yah (n = 11), female euploid (n = 12), female Ts66Yah (n = 15).

**Table 1:** Structural (extrinsic) mechanical properties of 6-, 9-, and 16-week-old Ts66Yah mice. Data are mean ± SEM. Asterisks indicate a significant difference compared to sex-matched euploid in pairwise comparisons with Sidak correction. **\* *p* < 0.05**. See Figure 3 for group numbers.

|  | Yield Force | Ultimate Force | Displacement to Yield | Postyield Displacement | Total Displacement | Stiffness | Work to Yield | Postyield Work | Total Work |
| --- | --- | --- | --- | --- | --- | --- | --- | --- | --- |
| **6-week-old** |  |  |  |  |  |  |  |  |  |
| Male Euploid | 11.03 ± 0.904 | 19.58 ± 0.882 | 114.02 ± 5.42 | 1146 ± 213 | 1260 ± 214 | 101.73 ± 5.52 | 0.687 ± 0.084 | 15.37 ± 2.27 | 16.06 ± 2.32 |
| Male Ts66Yah | **8.93 ± 0.414\*** | **15.12 ± 1.000\*** | 119.74 ± 5.10 | 833.18 ± 226.9 | 952.92 ± 225.4 | 80.59 ± 4.96 | 0.564 ± 0.037 | 9.85 ± 2.22 | 10.41 ± 2.22 |
| Female Euploid | 8.85 ± 0.385 | 14.79 ± 0.364 | 120.51 ± 6.37 | 1200 ± 246 | 1321 ± 249 | 79.87 ± 4.95 | 0.566 ± 0.039 | 10.80 ± 1.48 | 11.37 ± 1.49 |
| Female Ts66Yah | 9.40 ± 0.434 | 13.78 ± 0.404 | 132.20 ± 3.52 | 753.64 ± 140.6 | 885.83 ± 140.9 | 75.60 ± 3.43 | 0.656 ± 0.038 | 8.22 ± 1.03 | 8.87 ± 1.03 |
| **9-week-old** |  |  |  |  |  |  |  |  |  |
| Male Euploid | 13.63 ± 0.908 | 22.77 ± 0.885 | 133.95 ± 8.49 | 763.50 ± 147.0 | 897.45 ± 143.3 | 110.0 ± 4.50 | 1.01 ± 0.115 | 12.34 ± 1.88 | 13.34 ± 1.84 |
| Male Ts66Yah | 14.65 ± 1.267 | 21.20 ± 1.724 | 148.91 ± 4.95 | 346.50 ± 14.35 | 495.41 ± 14.87 | 104.72 ± 7.00 | 1.18 ± 0.129 | 6.58 ± 0.39 | 7.76 ± 0.50 |
| Female Euploid | 11.74 ± 0.829 | 17.03 ± 0.992 | 149.65 ± 7.53 | 892.23 ± 122.5 | 1042 ± 122 | 84.57 ± 4.14 | 0.959 ± 0.098 | 10.72 ± 0.88 | 11.68 ± 0.88 |
| Female Ts66Yah | 13.02 ± 0.424 | 16.97 ± 0.642 | 159.96 ± 6.42 | 695.89 ± 186.0 | 855.85 ± 187.1 | 87.94 ± 3.27 | 1.11 ± 0.069 | 8.22 ± 1.42 | 9.33 ± 1.42 |
| **16-week-old** |  |  |  |  |  |  |  |  |  |
| Male Euploid | 19.93 ± 1.53 | 27.11 ± 1.34 | 173.11 ± 14.9 | 275.00 ± 34.63 | 448.11 ± 27.42 | 126.46 ± 6.49 | 1.92 ± 0.311 | 6.81 ± 0.913 | 8.73 ± 0.848 |
| Male Ts66Yah | 17.84 ± 0.903 | 21.88 ± 1.20 | **192.69 ± 7.76\*** | 328.27 ± 53.35 | 520.96 ± 54.20 | **101.23 ± 6.55\*** | 1.82 ± 0.121 | 6.49 ± 0.974 | 8.31 ± 0.967 |
| Female Euploid | 18.57 ± 0.556 | 19.84 ± 0.652 | 199.35 ± 3.40 | 498.17 ± 60.28 | 697.52 ± 60.86 | 100.38 ± 3.87 | 2.00 ± 0.070 | 8.52 ± 0.882 | 10.51 ± 0.916 |
| Female Ts66Yah | 17.30 ± 0.630 | 19.06 ± 0.651 | 191.64 ± 5.99 | **325.73 ± 38.75\*** | **517.37 ± 37.96\*** | 98.08 ± 3.59 | 1.80 ± 0.106 | 5.63 ± 0.603 | 7.43 ± 0.600 |

**Table 2:** Material (intrinsic) mechanical properties of 6-, 9-, and 16-week-old Ts66Yah mice. Data are mean ± SEM. Asterisks indicate a significant difference compared to sex-matched euploid in pairwise comparisons with Sidak correction. ***\* p* < 0.05**. See Figure 3 for group numbers.

|  | Yield Stress | Ultimate Stress | Strain to Yield | Total Strain | Modulus | Resilience | Toughness |
| --- | --- | --- | --- | --- | --- | --- | --- |
| **6-week-old** |  |  |  |  |  |  |  |
| *Male Euploid* | 65.25 ± 3.60 | 117.28 ± 2.46 | 36860 ± 1749 | 406939 ± 69443 | 1.89 ± 0.083 | 1.30 ± 0.122 | 30.37 ± 3.94 |
| *Male Ts66Yah* | 64.84 ± 2.29 | 108.51 ± 3.59 | 36074 ± 1361 | 288994 ± 68900 | 1.92 ± 0.067 | 1.24 ± 0.077 | 22.73 ± 4.89 |
| *Female Euploid* | 69.62 ± 1.69 | 116.98 ± 2.22 | 37141 ± 11965 | 402180 ± 73850 | 2.04 ± 0.089 | 1.37 ± 0.088 | 28.28 ± 4.11 |
| *Female Ts66Yah* | **81.19 ± 3.61\*** | 118.77 ± 2.55 | 38400 ± 953 | 256826 ± 39572 | 2.24 ± 0.083 | 1.65 ± 0.100 | 22.30 ± 2.64 |
| **9-week-old** |  |  |  |  |  |  |  |
| *Male Euploid* | 81.55 ± 6.62 | 134.79 ± 5.38 | 30698 ± 1915 | 207424 ± 33870 | 2.86 ± 0.153 | 1.39 ± 0.175 | 17.75 ± 2.23 |
| *Male Ts66Yah* | 106.4 ± 5.96 | 152.7 ± 3.95 | 30634 ± 1362 | 101401 ± 2369 | 3.77 ± 0.206 | 1.74 ± 0.139 | 11.65 ± 0.500 |
| *Female Euploid* | 98.44 ± 6.41 | 141.1 ± 4.60 | 31593 ± 1549 | 218855 ± 23915 | 3.36 ± 0.176 | 1.69 ± 0.153 | 20.52 ± 1.35 |
| *Female Ts66Yah* | 113.9 ± 7.02 | 147.0 ± 6.67 | 31930 ± 1184 | 172033 ± 38007 | 3.86 ± 0.253 | 1.93 ± 0.132 | 16.38 ± 2.69 |
| **16-week-old** |  |  |  |  |  |  |  |
| *Male Euploid* | 109.0 ± 3.81 | 150.9 ± 6.48 | 317071 ± 3533 | 79006 ± 5153 | 4.09 ± 0.285 | 1.83 ± 0.234 | 8.49 ± 0.745 |
| *Male Ts66Yah* | **135.8 ± 8.97\*** | 163.4 ± 6.01 | 29306 ± 888.16 | 80175 ± 9012 | 4.99 ± 0.320 | 2.12 ± 0.172 | 9.54 ± 1.13 |
| *Female Euploid* | 158.2 ± 3.29 | 168.8 ± 3.60 | 30840 ± 630.10 | 108200 ± 9548 | 5.53 ± 0.168 | 2.63 ± 0.077 | 13.70 ± 0.994 |
| *Female Ts66Yah* | 150.2 ± 3.96 | 165.0 ± 2.12 | 28678 ± 717.26 | 77739 ± 5657 | 5.67 ± 0.120 | 2.34 ± 0.109 | **9.67 ± 0.734\*** |

At 9 weeks, no significant interactions between genotype and sex were detected for structural or material properties. At 16 weeks, significant interactions between genotype and sex were detected for displacement to yield (*p* = 0.050), postyield displacement (*p* = 0.030), total displacement (*p* = 0.009) and stiffness (*p* = 0.029). Pairwise comparisons between genotypes within sex showed female, but not male, Ts66Yah femurs had significantly lower postyield displacement and total displacement while male, but not female, Ts66Yah femurs had significantly increased displacement to yield and lower stiffness compared to sex-matched euploid femurs (Figure 3E; Table 1). For material properties at 16 weeks, significant interactions between genotype and sex were detected for yield stress (*p* = 0.002) and toughness (*p* = 0.013). Pairwise comparisons between genotypes within sex showed male Ts66Yah femurs had significantly higher yield stress while female Ts66Yah mice had significantly lower toughness (Figure 3F; Table 2).

Together, Ts66Yah mice appear to have skeletal deficits depending on age, bone compartment, and sex, as similarly illustrated in previously characterized DS mouse models. At 6-weeks-old, only male Ts66Yah mice had trabecular deficits while both sexes had cortical deficits related to overall size, although male Ts66Yah mice had more deficits in cortical geometry than female Ts66Yah mice. This was supported by trabecular (Figure 4A-B) and cortical (Figure 4C-D) composite scores that utilized each trabecular or cortical variable in sex-stratified principal components analyses (PCA) to determine the principal component 1 loading value of each variable (Supplemental Table 3, Supplemental Figure 2), then calculating a single value for each animal to represent each bone compartment using these loadings and the raw value of each variable (see Equations 1-3 in Methods). These morphological changes were sufficient to result in reduced overall strength in femurs of male Ts66Yah mice, but when corrected using cortical geometry, only female Ts66Yah mice showed material alterations in elastic deformation, indicating greater stiffness.

**Figure 4:**
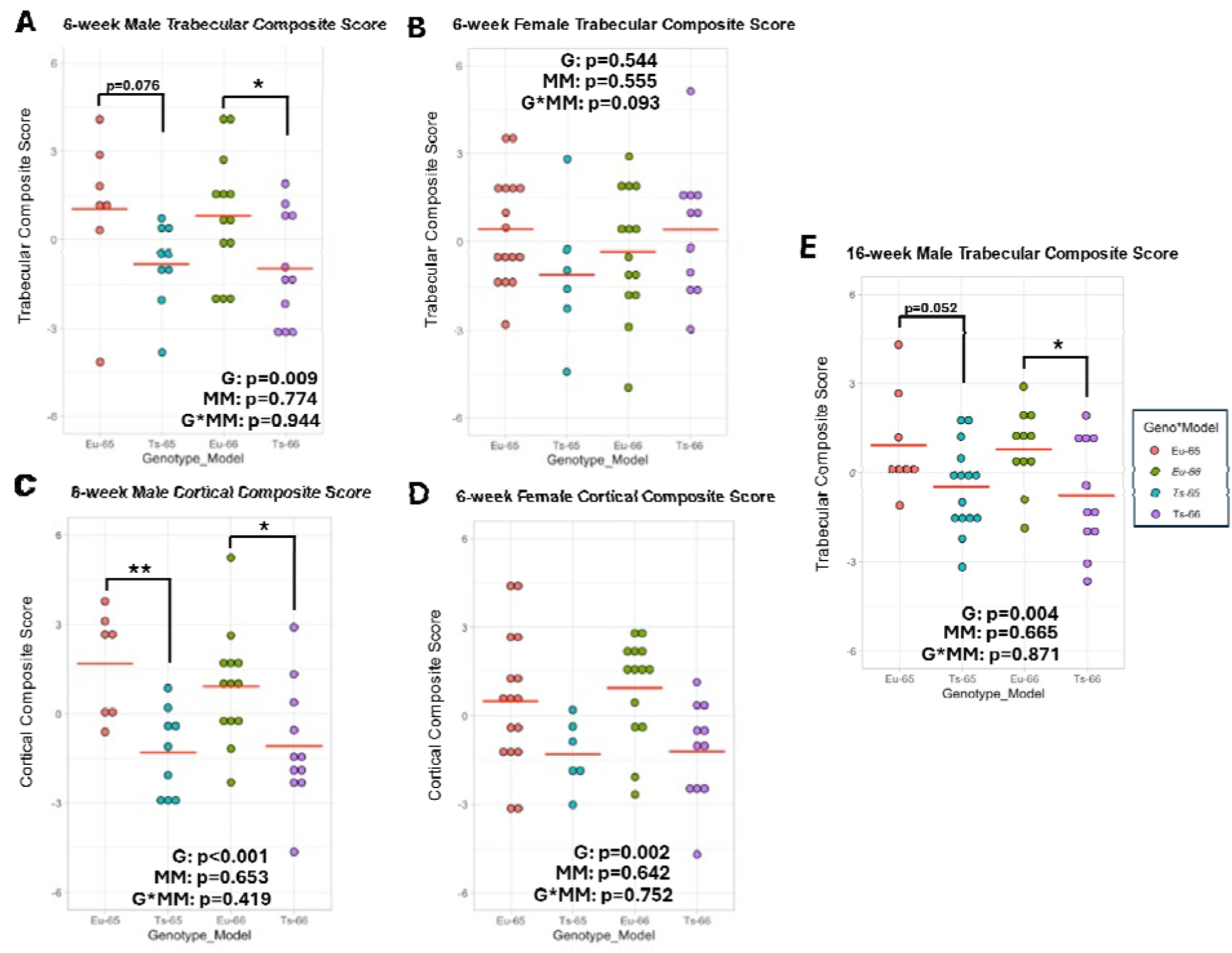
Trabecular and cortical composite scores of 6- and 16-week-old Ts65Dn and Ts66Yah mice. A-B) 6-week-old trabecular composite scores. **C**-**D)** 6-week-old cortical composite scores. **E)** 16-week-old male trabecular composite scores. Red horizontal line indicates group mean. Asterisks indicate a significant difference between groups in contrast analysis with Bonferroni correction after performing two-way ANOVAs with genotype (G) and mouse model (MM) as between subject factors. * *p* < 0.05, ** *p* < 0.01.

By 9 weeks of age, there were fewer differences between Ts66Yah and euploid mice. In male mice, largely no differences were seen in trabecular bone microarchitecture, other than decreased thickness, suggesting a normalization from the 6-week structure. This was supported by the trabecular composite score, which was not significantly different between male euploid and Ts66Yah mice (Supplemental Figure 3A; *p* = 0.107). Female Ts66Yah mice at 9 weeks had no significant differences in individual trabecular variables or trabecular composite score (Supplemental Figure 3A; *p =* 0.272) compared to female euploid mice, similar to findings at 6 weeks. However, female Ts66Yah mice appeared to catch up to female euploid mice and even surpassed them in terms of overall proportion of cortical bone, although this did not translate to any large differences in cortical composite score (Supplemental Figure 3B; *p* = 0.544) or mechanics. Cortical bone geometry alterations persisted in male Ts66Yah mice, which were supported by a significantly decreased cortical composite score (Supplemental Figure 3B; *p* = 0.010); however, changes on the endocortical surface appeared to mostly alleviate mechanical differences. The only deficit to newly appear in 9-week-old Ts66Yah mice was a significantly shorter femur.

Finally, deficits in femoral length, trabecular microarchitecture, and cortical geometry in male Ts66Yah mice persisted or reappeared at 16 weeks of age whereas female Ts66Yah mice did not significantly differ from female euploid mice. These trends in trabecular and cortical structure were supported by significantly decreased trabecular and cortical composite scores (*p =* 0.015 and < 0.001, respectively) in male Ts66Yah mice compared to male euploid mice and no significant differences in trabecular and cortical composite scores (*p =* 0.559 and 0.944, respectively) between female euploid and Ts66Yah mice (Supplemental Figure 3C-D). Mechanically, 16-week-old male Ts66Yah mice presented with more deficits in the elastic region than at 6 weeks of age, showing both structural and estimated tissue level brittleness. Despite a lack of significant differences in cortical geometry, female Ts66Yah mice showed subtle changes in postyield deformation based on structural mechanics and overall inability to absorb energy on an estimated tissue level.

### Ts65Dn and Ts66Yah mice are similar in their bone deficits at 6 and 16 weeks

#### 6 weeks of age

To find potential effects of the triplicated, non-Hsa21 homologous genes in the Ts65Dn as compared to the Ts66Yah mouse bone structural phenotypes, trabecular bone variables were standardized for euploid Ts65Dn littermates, Ts65Dn mice, euploid Ts66Yah littermates, and Ts66Yah mice and then analyzed by principal component analysis (PCA) separately for male and female mice (Supplemental Figure 2A-B). Principal component 1 (PC1) was found to account for 92.68% of the variance for 6-week male trabecular bone variables and 89.09% of the variance for female trabecular bone variables (Supplemental Table 3). The coefficients (loadings) for each of the five variables (BMD, BV/TV, Tb.Th, Tb.Sp, and Tb.N) for PC1 were nearly equal in magnitude for both sexes. However, Tb.Sp had a reverse sign compared to the other variables. This is expected as Tb.Sp has been shown to be inversely correlated with the other trabecular variables, and a higher trabecular separation is considered a skeletal deficit (SLOAN *et al*. 2023).

Because the first principal component (PC1) explained most of the variance in the trabecular bone variables, it was utilized to develop the trabecular composite score. These composite scores were assessed by 2-way ANOVA with mouse model (Ts65Dn, Ts66Yah) and genotype (euploid, trisomic), along with their interaction, as between-subject factors for each sex. The interaction term enabled testing whether the difference in average trabecular composite score between euploid and trisomic mice differed between the two mouse models. The interaction was not significant for male (*p* = 0.944) or female (*p* = 0.093) mice. The mouse model main effect was also not significant for either sex (male *p* = 0.774, female *p* = 0.555), which was due to the standardization procedure. For male mice, there was a significant effect (*p* = 0.009) of genotype on trabecular composite score; notably, for female mice, the genotype effect was not significant (*p* = 0.544). This indicated that Ts65Dn and Ts66Yah mice at 6 weeks have similar genotypic effects on trabecular bone structure, but only male mice exhibit significant deficits relative to respective euploid controls (Figure 4A-B).

PCA was performed separately for standardized cortical bone variables (Tt.Ar, Ma.Ar, Ct.Ar, Ct.Th, Ps.Pm, Ec.Pm, and Ct.TMD) of 6-week-old male and female Ts65Dn and Ts66Yah mice (Supplemental Figure 2C-D). PC1 accounted for a lower percentage of the variance than for trabecular variables at 67.3% and 61.94% for male and female mice, respectively (Supplemental Table 3). Two of the seven variables, Ct.Th and Ct.TMD, contribute less to PC1 than the other five, which have roughly equal contributions. This was not surprising given that Ct.TMD and Ct.Th are not always affected by trisomy depending on the mouse model and age (FOWLER *et al*. 2012; THOMAS *et al*. 2020; THOMAS *et al*. 2021; SHERMAN *et al*. 2022; LACOMBE *et al*. 2024; LAMANTIA *et al*. 2024).

The cortical composite score was developed using the PC1 loadings for each variable and then assessed by 2-way ANOVA, which showed a significant effect of genotype in male (*p <* 0.001) and female (*p* = 0.002) mice. Neither sex exhibited a mouse model effect (male *p =* 0.653, female *p* = 0.642) nor interaction between genotype and mouse model (male *p =* 0.419, female *p =* 0.752). These outcomes indicate Ts65Dn and Ts66Yah mice have similar genotypic effects on cortical bone geometry at 6 weeks, with deficits evident in both sexes, differing from trabecular deficits presenting only in males of both models (Figure 4C-D).

### 16 weeks of age

The same method was performed using trabecular variables from 16-week-old male Ts65Dn and Ts66Yah mice (Supplemental Figure 2E), although BMD was excluded due to differing methodologies resulting in an areal BMD for Ts65Dn data and a volumetric BMD for Ts66Yah data. PC1 accounted for 74.37% of the variance at this age (Supplemental Table 3). Tb.Th contributed less to the proportion of variance at 16 weeks than at 6 weeks, and Tb.Sp was still reversed in its sign. Two-way ANOVA of the male trabecular composite scores showed a significant genotype effect (*p =* 0.004) but no mouse model effect (*p =* 0.665) or interaction (*p*= 0.871). This indicates male Ts65Dn and Ts66Yah mice do not differ in the genotypic effects on trabecular microarchitecture at 16 weeks and both DS models are at a deficit compared to euploid mice (Figure 4E). Female trabecular structure could not be compared due to a lack of data in female Ts65Dn mice at this age. Additionally, cortical measures could not be compared in either sex using this method due to a lack of comparable measures between Ts65Dn and Ts66Yah mice.

### Treatment with Leucettinib-21, a DYRK1A inhibitor, did not improve skeletal deficits in Ts66Yah mice at postnatal day 36

Because of the similarities between Ts66Yah and Ts65Dn bone structure at 6 weeks and our previous work describing the window of trisomic *Dyrk1a* modulation (LACOMBE *et al*. 2024), male and female mice were treated with 0.5mg/kg of Leucettinib-21 (L21), a DYRK1A inhibitor, daily from postnatal day (P) 21 until P36 by oral gavage. In a repeated measures ANOVA based on daily body weight, significant interactions were observed for postnatal day, sex, and treatment (*p* = 0.006), postnatal day and genotype (*p* = 0.042), postnatal day and sex (*p* < 0.001), and postnatal day and treatment (*p* = 0.012). In pairwise comparisons of each postnatal day within genotype, sex, and treatment, all groups increased body weight over the course of treatment (Figure 5A-B). Comparisons between treatments within each sex and genotype found male Ts66Yah mice treated with L21 had significantly lower body weight compared to male Ts66Yah mice treated with the vehicle starting at P21 and continuing through P35; no other groups had significant differences between treatments (Figure 5A). In comparisons between genotypes within each sex and treatment, there were no genotype differences in body weight in either sex of vehicle-treated mice. This indicates there may have been differences between the male Ts66Yah treatment groups present prior to L21 administration despite randomization of litters to each treatment. To understand if L21 treatment contributed to differences in overall weight gain, change in body weight over L21 treatment was calculated by subtracting weight at P21 from weight at P36. A three-way ANOVA with genotype, sex, and treatment as main effects found only a significant sex and treatment interaction (*p* = 0.013). Pairwise comparisons between treatments within genotype and sex found only male mice of both genotypes had significantly lower weight gains when treated with L21, and comparisons between genotypes within vehicle-treated mice found no significant differences in either sex (Figure 5C).

**Figure 5:**
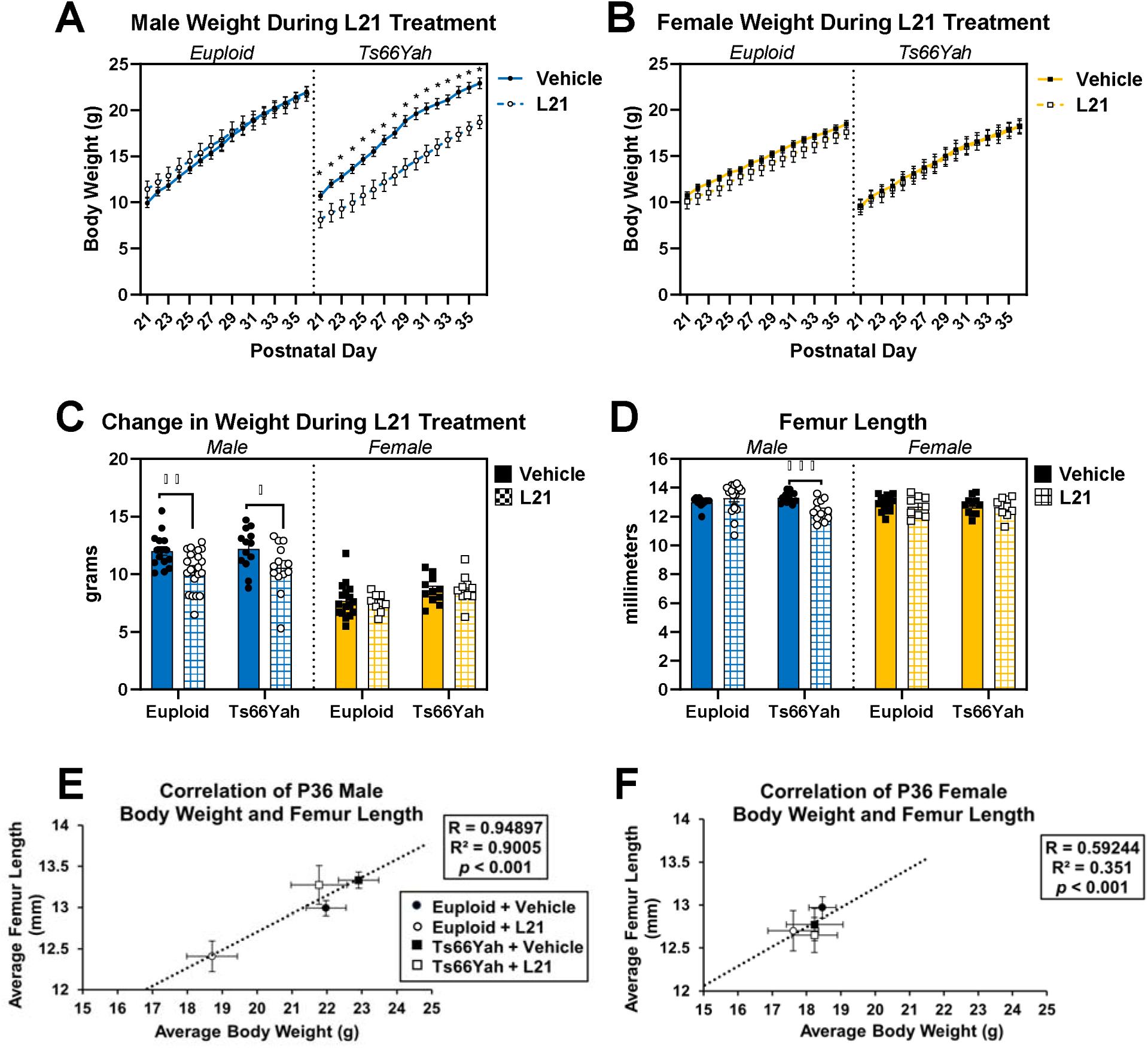
Body weight and femur lengths of Leucettinib-21-treated Ts66Yah mice. **A-B)** Body weight during treatment, beginning with postnatal day (P) 21 and ending with P35. Comparisons between each postnatal day within genotype, sex, and treatment showed significant increases with each day in all 8 groups. **C)** Change in body weight calculated as body weight at P21 subtracted from body weight at P36 for each animal. **D)** Femur length at P36. **E-F)** Correlation analysis between body weight and femur length at P36. Data are mean ± SEM. Asterisks indicate a significant difference between groups in pairwise comparisons with Sidak correction. * *p* < 0.05, ** *p* < 0.01,*** *p* < 0.001. Male mice: vehicle-treated euploid (n = 15 [weights] or 13 [femur length]), L21-treated euploid (n = 20 [weights] or 18 [femur length]), vehicle-treated Ts66Yah (n = 13), L21-treated Ts66Yah (n = 13). Female mice: vehicle-treated euploid (n = 17), L21-treated euploid (n = 9), vehicle-treated Ts66Yah (n = 11), L21-treated Ts66Yah (n = 10).

For femur length, there was a three-way interaction between genotype, sex, and treatment (*p* = 0.012). Similar to body weight, male L21-treated Ts66Yah mice had significantly shorter femurs than male vehicle-treated Ts66Yah mice in comparisons between treatments within each sex and genotype, and there were no significant genotype differences within either sex of vehicle-treated mice in comparisons between genotypes within sex and treatment (Figure 5D). However, based on the significant positive correlation between body weight and femur length at P36 (Figure 5E-F), the low body weight of male L21-treated Ts66Yah mice at the beginning of treatment may have influenced femoral length and structure independently of L21 treatment.

For trabecular structure, no significant three-way interactions between genotype, sex, and treatment were detected, but significant interactions between genotype and sex were found for BMD (Supplemental Figure 4B; *p* = 0.016), BV/TV (Supplemental Figure 4C; *p* = 0.017), and Tb.Th (Supplemental Figure 4D; *p* = 0.017). In all three variables, comparisons between treatment within genotype and sex indicated that male L21-treated Ts66Yah mice had significantly lower values than male vehicle-treated Ts66Yah mice, contrary to our hypothesis. There were no significant genotype differences in vehicle-treated mice for either sex. This was confirmed by the trabecular composite score, which was not significantly different between vehicle-treated euploid and Ts66Yah mice of either sex (Supplemental Figure 3E; Male: *p* = 0.294; Female: *p* = 0.705). The lack of genotype differences in trabecular structure could explain the adverse effect of DYRK1A inhibition; previous studies have shown haploinsufficiency of *Dyrk1a* can result in skeletal structure deficits (OTTE AND ROPER 2024).

For cortical geometry, three-way interactions between genotype, sex, and treatment were detected for Ct.Ar (Figure 6D; *p* = 0.008), Ct.Th (Figure 6F; *p* = 0.006), maximum moment of inertia (I_max_; Figure 6I; *p* = 0.017), and minimum moment of inertia (I_min_; Figure 6J; *p* = 0.043), and an additional genotype and treatment interaction was detected for Ct.TMD (Figure 6K; *p* = 0.008). In pairwise comparisons between treatments within genotype and sex, male L21-treated Ts66Yah mice again had significantly lower Ct.Ar, Ct.Th, I_max_, and I_min_ compared to male vehicle-treated Ts66Yah mice. Both sexes of L21-treated Ts66Yah mice had significantly greater Ct.TMD compared to sex-matched vehicle-treated Ts66Yah mice. In comparisons between genotypes within sex and treatment, the only significant genotype difference in vehicle-treated mice was for I_max_, where female vehicle-treated Ts66Yah mice were significantly lower than female vehicle-treated euploid mice. The lack of genotype differences in male vehicle-treated mice was confirmed by the cortical composite score, which was not significantly different between male euploid and Ts66Yah mice (Supplemental Figure 3F; *p* = 0.129). While there was no significant difference in any of the geometric variables, the cortical composite score was significantly decreased in female Ts66Yah mice compared to female euploid mice (Supplemental Figure 3F; *p* = 0.006), and in conjunction with the significantly lower I_max_, an indicator of strength based on cortical geometry, this suggests there may be a subclinical cortical structure phenotype at P36.

**Figure 6:**
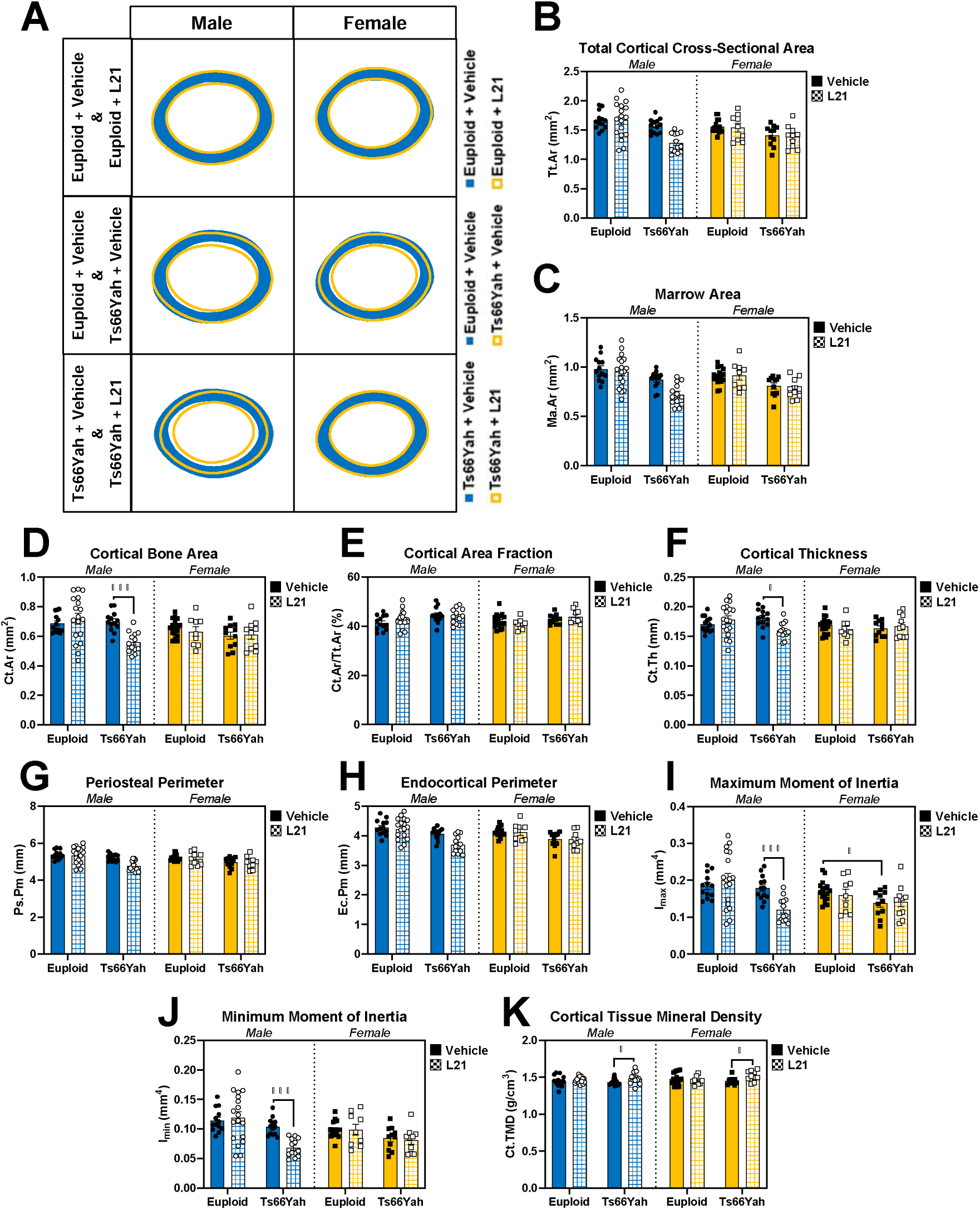
Cortical bone variables for Leucettinib-21-treated Ts66Yah mice. **A)** Representations of the average cortical cross-section for each group made by calculating the centroid of each cortical section and finding the distance from the centroid to the endocortical and periosteal surfaces every 0.5 degrees. The average is found for each animal, then the average of the group is found to make the radar graph. **B-K)** Data are mean ± SEM. Asterisks indicate a significant difference between groups in pairwise comparisons with Sidak correction. * *p* < 0.05, * *p* < 0.001. Male mice: vehicle-treated euploid (n = 13), L21-treated euploid (n = 18), vehicle-treated Ts66Yah (n = 13), L21-treated Ts66Yah (n = 13). Female mice: vehicle-treated euploid (n = 17), L21-treated euploid (n = 9), vehicle-treated Ts66Yah (n = 11), L21-treated Ts66Yah (n = 10).

Treatment with Leucettinib-21 may have stunted growth as shown by gain in weight, femoral length, trabecular microarchitecture, and most cortical geometry, contrary to the initial hypothesis. This may be due to inappropriate treatment timing, where DYRK1A inhibition when DYRK1A was not overexpressed resulted in haploinsufficiency, or the low body weight of the male Ts66Yah mice treated with L21 at P21 impacted bone structure. Interestingly, these effects were specific to male mice; female Ts66Yah mice were minimally affected by treatment. Cortical density, indicated by Ct.TMD, did improve with treatment in Ts66Yah mice, indicating alterations in treatment timing or dosage may provide better results.

### Germline reduction of Dyrk1a copy number improves cortical deficits in male Ts66Yah mice at postnatal day 36

The unexpected results of Leucettinib-21 treatment raised the question of whether *Dyrk1a* was contributing to structural phenotypes in Ts66Yah femurs in a manner similar to Ts65Dn mice. To evaluate the role of trisomic *Dyrk1a* in the appendicular skeleton of Ts66Yah mice and see if this role was similar to trisomic *Dyrk1a* in Ts65Dn mice, Ts66Yah femurs with a germline reduction of *Dyrk1a* copy number were analyzed at P36. Three-way interactions between genotype, sex, and *Dyrk1a* reduction were detected for body weight (*p* = 0.002) and femur length (*p* < 0.001). Pairwise comparisons between genotypes within sex and *Dyrk1a* reduction showed male Ts66Yah mice with three copies of *Dyrk1a* (Ts66Yah) had significantly lower body weights and shorter femurs than male euploid mice with two copies of *Dyrk1a* (euploid) (Supplemental Figure 5). In comparisons between *Dyrk1a* reduction within sex and genotype, male Ts66Yah mice with two copies of *Dyrk1a* (Ts66Yah,*Dyrk1a*^+/+/-^) had significantly longer femurs than male Ts66Yah mice, but no difference in body weight. These comparisons also found both sexes of euploid mice with one copy of *Dyrk1a* (euploid,*Dyrk1a*^+/-^) were significantly smaller in weight and had shorter femurs than sex-matched euploid mice. These data support the hypothesis that normalization of *Dyrk1a* copy number can improve femur length but not body weight in male Ts66Yah at P36 and that haploinsufficiency of *Dyrk1a* negatively affects overall weight and femur length in both sexes.

All trabecular bone variables had significant three-way interactions between genotype, sex, and *Dyrk1a* reduction (BMD: *p* = 0.014; BV/TV: *p* = 0.008; Tb.Th: *p* = 0.012; Tb.Sp: *p* = 0.007; Tb.N: *p* = 0.037; Supplemental Figure 6B-F). Pairwise comparisons between genotypes within sex and *Dyrk1a* reduction showed no significant differences in either sex between euploid and Ts66Yah mice, but comparisons between *Dyrk1a* reduction within genotype and sex found male euploid,*Dyrk1a*^+/-^ mice to be deficient in all variables compared to male euploid mice. Most cortical variables had significant three-way interactions between genotype, sex, and *Dyrk1a* reduction (Tt.Ar, Ma.Ar, Ct.Ar, Ps.Pm, Ec.Pm, I_max_, I_min_: *p* < 0.001; Ct.Th: *p* = 0.002) or a significant interaction between genotype and *Dyrk1a* reduction (Ct.Ar/Tt.Ar: *p* = 0.036) except for Ct.TMD. In pairwise comparisons between genotypes within sex and *Dyrk1a* reduction of variables with three-way interactions, male Ts66Yah mice were significantly lower than male euploid mice in most (Figure 7B-D,F-J), while Ct.Ar/Tt.Ar was significantly increased in male Ts66Yah mice compared to male euploid mice (Figure 7E). In comparisons between *Dyrk1a* reduction within genotype and sex, Ts66Yah,*Dyrk1a*^+/+/-^ mice were significantly improved in all variables with a three-way interaction compared to Ts66Yah mice except for Ct.Th, which was not significantly different. These comparisons also found that haploinsufficiency of *Dyrk1a* in euploid mice resulted in significantly lower Tt.Ar, Ma.Ar, Ps.Pm, Ec.Pm, I_max_, and I_min_ in male mice and significantly lower Ct.Ar, Ct.Ar/Tt.Ar, and Ct.Th in both sexes compared to sex-matched euploid mice.

**Figure 7:**
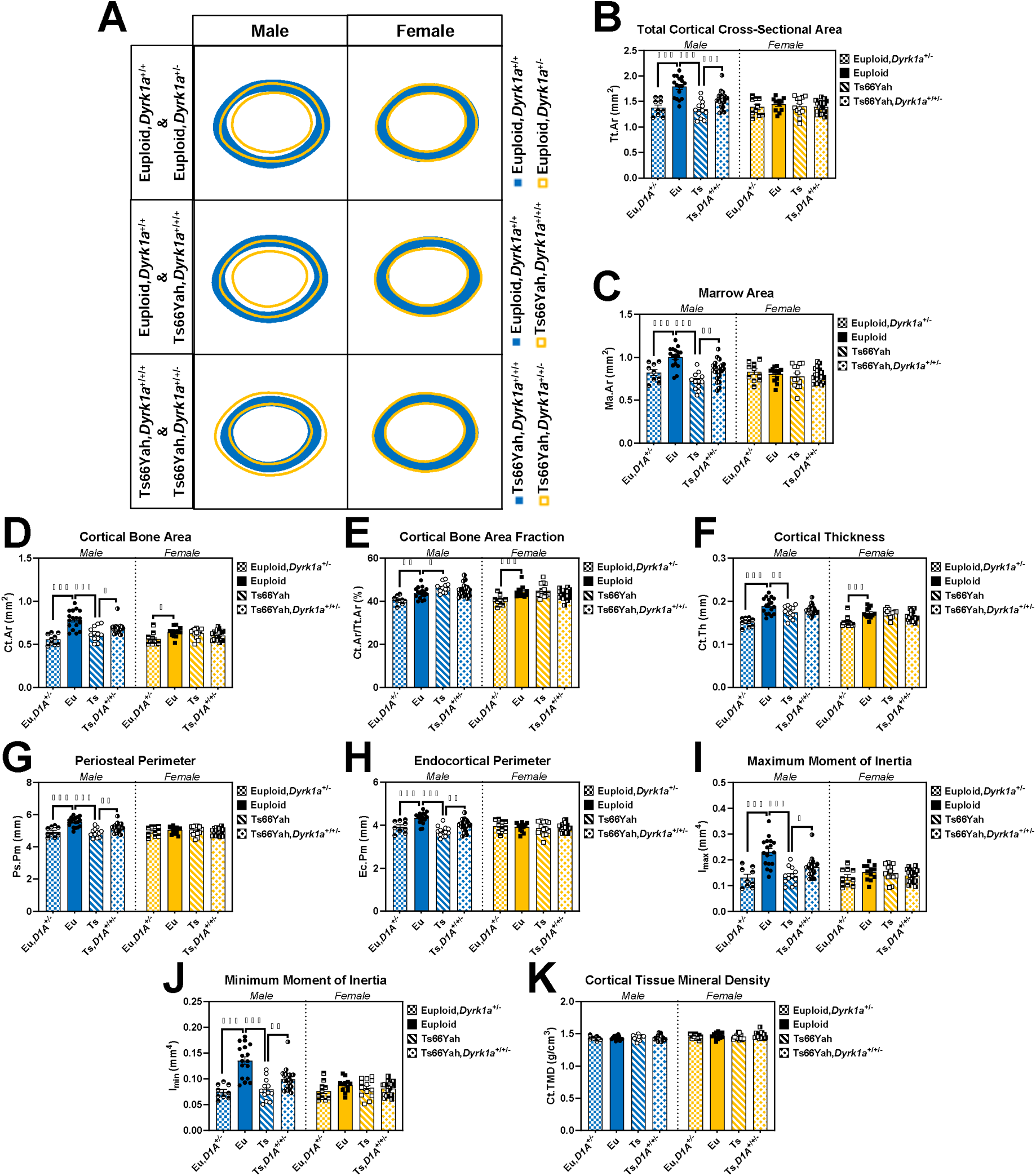
Cortical bone variables for P36 Ts66Yah,*Dyrk1a*^+/+/-^ mice. **A)** Representations of the average cortical cross-section for each group made by calculating the centroid of each cortical section and finding the distance from the centroid to the endocortical and periosteal surfaces every 0.5 degrees. The average is found for each animal, then the average of the group is found to make the radar graph. **B-K)** Data are mean ± SEM. Asterisks indicate a significant difference between groups in pairwise comparisons with Sidak correction. * *p* < 0.05, ** *p* < 0.01, *** *p* < 0.001. Male mice: euploid (n = 17); euploid,*Dyrk1a*^+/-^ (n = 9); Ts66Yah (n = 12); Ts66Yah,*Dyrk1a*^+/+/-^ (n = 21). Female mice: euploid (n = 12); euploid,*Dyrk1a*^+/-^ (n = 10); Ts66Yah (n = 13); Ts66ah,*Dyrk1a*^+/+/-^ (n = 19).

Overall, no trabecular deficits were seen at P36 in either sex of Ts66Yah mice and only male Ts66Yah mice had cortical deficits at P36, unlike vehicle-treated Ts66Yah mice. This was supported by the trabecular and cortical composite scores, where neither sex had significant differences in trabecular composite score (Figure 8A-B; Male: *p =* 0.207; Female: *p* = 0.400) and only male Ts66Yah mice had a significant decrease in cortical composite score (Figure 8C-D; Male: *p* < 0.001; Female: *p* = 0.614). Copy number reduction of *Dyrk1a* had differing effects at P36 depending on ploidy, bone compartment, and sex. In the trabecular region, female mice were not affected by *Dyrk1a* reduction; in the cortical region, female euploid mice, but not female Ts66Yah mice, were negatively affected by *Dyrk1a* reduction, indicating trisomic *Dyrk1a* does not play a significant role in the protective effect of female sex in Ts66Yah mice at this timepoint. Only euploid male mice were negatively affected by *Dyrk1a* reduction in the trabecular region. In the cortical region, while male euploid mice were negatively affected by *Dyrk1a* reduction (more so than females), male Ts66Yah mice were positively affected by *Dyrk1a* reduction in most variables, although this did not reach the level of euploid mice with two copies of *Dyrk1a*. Together, this indicates that *Dyrk1a* and other trisomic genes are contributing to cortical deficits in male Ts66Yah mice at P36, which differs from cortical results in male Ts65Dn and Ts65Dn,*Dyrk1a*^+/+/-^ mice at P36, where reduction in *Dyrk1a* in an otherwise trisomic mouse did not improve cortical deficits (LACOMBE *et al*. 2024).

**Figure 8:**
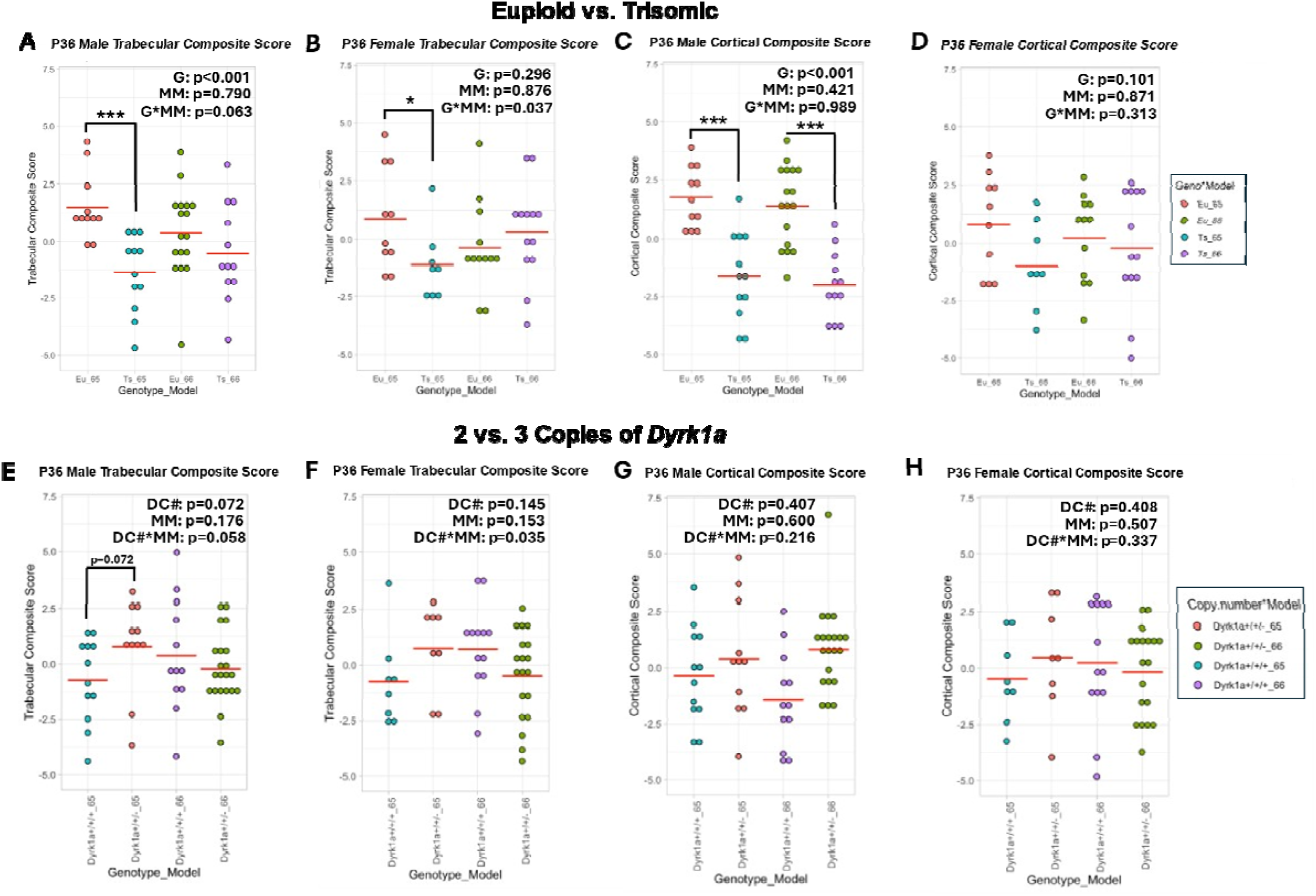
Trabecular and cortical composite scores for P36 Ts65Dn, Ts66Yah, and germline reduction of *Dyrk1a* copy number mice. **A-D** Comparisons between euploid and trisomic mice. Two-way ANOVA with genotype (G) and mouse model (MM) as between subject factors. **E-H)** Comparisons between trisomic mice with 2 (*Dyrk1a*^+/+/-^) or 3 (*Dyrk1a*^+/+/+^) copies of *Dyrk1a*. Two-way ANOVA with *Dyrk1a* copy number (DC#) and mouse model (MM) as between subject factors. Red horizontal line indicates group mean. Asterisks indicate significant difference between genotypes or *Dyrk1a* copy number in mouse model-stratified contrast analysis. * *p* < 0.05, *** *p* < 0.001.

### Differences between Ts65Dn and Ts66Yah structural phenotypes and effects of Dyrk1a normalization are present at P36

For P36 data, PCA was performed first on Ts65Dn and Ts66Yah mice with three copies of *Dyrk1a* and their respective euploid controls with two copies of *Dyrk1a*, as described for 6-week data (Supplemental Figure 7C-F). PC1 accounted for 85.6% and 85.5% of variance for male and female trabecular variables, respectively (Supplemental Table 4). In both sexes, the loading of each trabecular variable was similar in magnitude with Tb.Th contributing slightly less and Tb.Sp still reversed in sign. Two-way ANOVA of male trabecular composite scores showed a significant genotype effect (*p* = 0.001) and no mouse model effect (*p* = 0.790), confirming that male euploid mice had higher trabecular composite scores than male trisomic mice (Figure 8A). The interaction between genotype and mouse model approached significance (*p* = 0.063), and contrast analysis within each mouse model indicated trisomic Ts65Dn mice had a significantly lower trabecular composite score than their euploid littermates (*p* < 0.001) while trisomic Ts66Yah mice were not significantly different from their euploid littermates (*p* = 0.207). Female trabecular composite scores showed a significant interaction between genotype and mouse model (*p* = 0.037), due to Ts65Dn mice showing a significantly lower score compared to euploid mice (*p =* 0.042) and Ts66Yah showing a non-significant, but higher score (*p* = 0.400) than their respective euploid control mice (Figure 8B). For trabecular structure, male Ts65Dn and Ts66Yah models do not differ in their genotypic effects at P36, with both having deficits in trabecular microarchitecture compared to euploid mice, but female Ts65Dn and Ts66Yah differ in their trabecular microarchitecture compared to euploid mice. This suggests the trisomic, non-Hsa21 orthologous genes in female Ts65Dn mice may contribute to the trabecular deficits at P36.

For cortical variables, PC1 accounted for 77.7% and 68.5% of variance for male and female mice, respectively (Supplemental Table 4). In male mice, the loadings of most cortical variables were similar in magnitude with Ct.Th contributing slightly less and Ct.TMD contributing very little and reversed in sign. In female mice, the loadings were similar for all but Ct.Th and Ct.TMD; these were similar to each other except that Ct.TMD was reversed in sign. Two-way ANOVA of cortical composite scores found a significant genotype effect in male mice (*p* < 0.001) but not female mice (*p* = 0.101) and no significant effect of mouse model (male *p* = 0.421, female *p* = 0.871) or interaction (male *p* = 0.989, female *p* = 0.313) in either sex. This indicates the Ts65Dn and Ts66Yah models do not differ in their genotypic effects or lack thereof on cortical geometry at P36 and only male trisomic mice show deficits compared to euploid mice while female euploid and trisomic mice are not different (Figure 8C-D).

Next, PCA was performed on P36 Ts65Dn, Ts66Yah, and germline reduction of *Dyrk1a* copy number mice of each mouse model to understand if the effects of *Dyrk1a* copy number reduction from three copies to two varied between the two mouse models (Supplemental Figure 7G-J). For trabecular variables, PC1 accounted for 80.6% and 85.1% of variance for male and female mice, respectively (Supplemental Table 4). In both sexes, the loadings of all trabecular variables except Tb.Th were similar in magnitude with Tb.Th contributing slightly less and Tb.Sp being reversed in sign. There was no significant main effect of mouse model or *Dyrk1a* copy number on trabecular composite score in either sex (male *p* = 0.176 and 0.072, female *p* = 0.153 and 0.145) by two-way ANOVA. Notably, however, there was a significant interaction between mouse model and *Dyrk1a* copy number (*p* = 0.035) in female mice, where Ts65Dn mice with 2 copies of *Dyrk1a* had a higher trabecular composite score whereas Ts66Yah mice with 2 copies of *Dyrk1a* had a lower trabecular composite score compared to their respective trisomic mice with 3 copies of *Dyrk1a* (Figure 8F). The interaction approached significance (*p* = 0.058) in male mice, showing a similar trend as female mice (Figure 8E). Together, this indicated that the effects of germline normalization of *Dyrk1a* copy number on trabecular bone at P36 was dependent on mouse model and similar between sexes, where normalization was beneficial for Ts65Dn trabecular microarchitecture and negative for Ts66Yah trabecular microarchitecture.

For cortical variables, PC1 accounted for 73.7% and 69.3% of the variance for male and female mice, respectively (Supplemental Table 4). In both sexes, the loadings of all cortical variables except Ct.Th and Ct.TMD were similar in magnitude with Ct.Th contributing less and Ct.TMD contributing the least. However, Ct.Th seems to contribute more to male variance than female variance whereas the opposite is true for Ct.TMD. The sign of Ct.TMD was reversed in both sexes. There were no significant main effects of mouse model (male *p =* 0.600, female *p* = 0.507) or *Dyrk1a* copy number (male *p* = 0.407, female *p* = 0.408), nor a significant interaction between them (male *p* = 0.216, female *p =* 0.337) on cortical composite score in either sex by two-way ANOVA, indicating *Dyrk1a* copy number does not significantly rescue cortical deficits in either mouse model at P36 (Figure 8G-H).

## Discussion

### Ts66Yah mice as a model for DS skeletal deficits

Previous research demonstrated that male Ts65Dn and Ts66Yah had similar cranial and mandibular deficits compared to euploid mice, but these deficits were less severe in male Ts66Yah mice compared to Ts65Dn mice (DUCHON *et al*. 2022). In the current study of femurs, only male Ts66Yah mice displayed deficits in trabecular bone structure at 6 and 16 weeks (Table 3). Previous studies in Ts65Dn mice reported deficits in trabecular microarchitecture at 6 weeks in both sexes (THOMAS *et al*. 2021) and at 16 weeks in male mice (BLAZEK *et al*. 2011). The mature skeletal phenotype in female Ts65Dn mice is unknown, but female Dp1Tyb mice do not show trabecular deficits at 16 weeks (THOMAS *et al*. 2020), consistent with current findings in female Ts66Yah mice. Male Ts65Dn mice also show significant trabecular microarchitecture deficits at P36 (LACOMBE *et al*. 2024), which contrasts with current findings in male Ts66Yah mice (Table 3). While Ts65Dn mice have not been investigated previously at exactly 9 weeks of age, areal BMD was lower at the proximal tibia (including trabecular and cortical bone) in 8-week-old male Ts65Dn mice (WILLIAMS *et al*. 2018). This differs, perhaps due to differences in methodology and/or age, from what was found in Ts66Yah mice at 9 weeks of age, where skeletal phenotypes of the femur seem to mostly normalize (Table 3). Bone growth is still occurring between 8 and 9 weeks in mice (JILKA 2013), and areal measurements of BMD can overestimate the severity of the deficit in DS due to a smaller bone size (CARFI *et al*. 2017; GARCIA-HOYOS *et al*. 2017). Additionally, the trabecular BMD reported in this paper and others from our lab does not include cortical bone whereas the BMD measurement of Williams *et al*. (2018) does. Since the tibial midshaft (cortical bone only) also has low BMD, it is possible the cortical bone included at the proximal tibia is driving the low BMD. Together, these data suggest the additional genes present in Ts65Dn mice result in earlier trabecular deficits in both sexes of mice. One gene located within the trisomic Mmu17 centromeric region of Ts65Dn mice, *Fndc1,* has been implicated in contributing to trabecular bone loss in ovariectomized rats when highly expressed (XIAO *et al*. 2019) and may play a similar role when present in three copies in Ts65Dn mice.

**Table 3:**
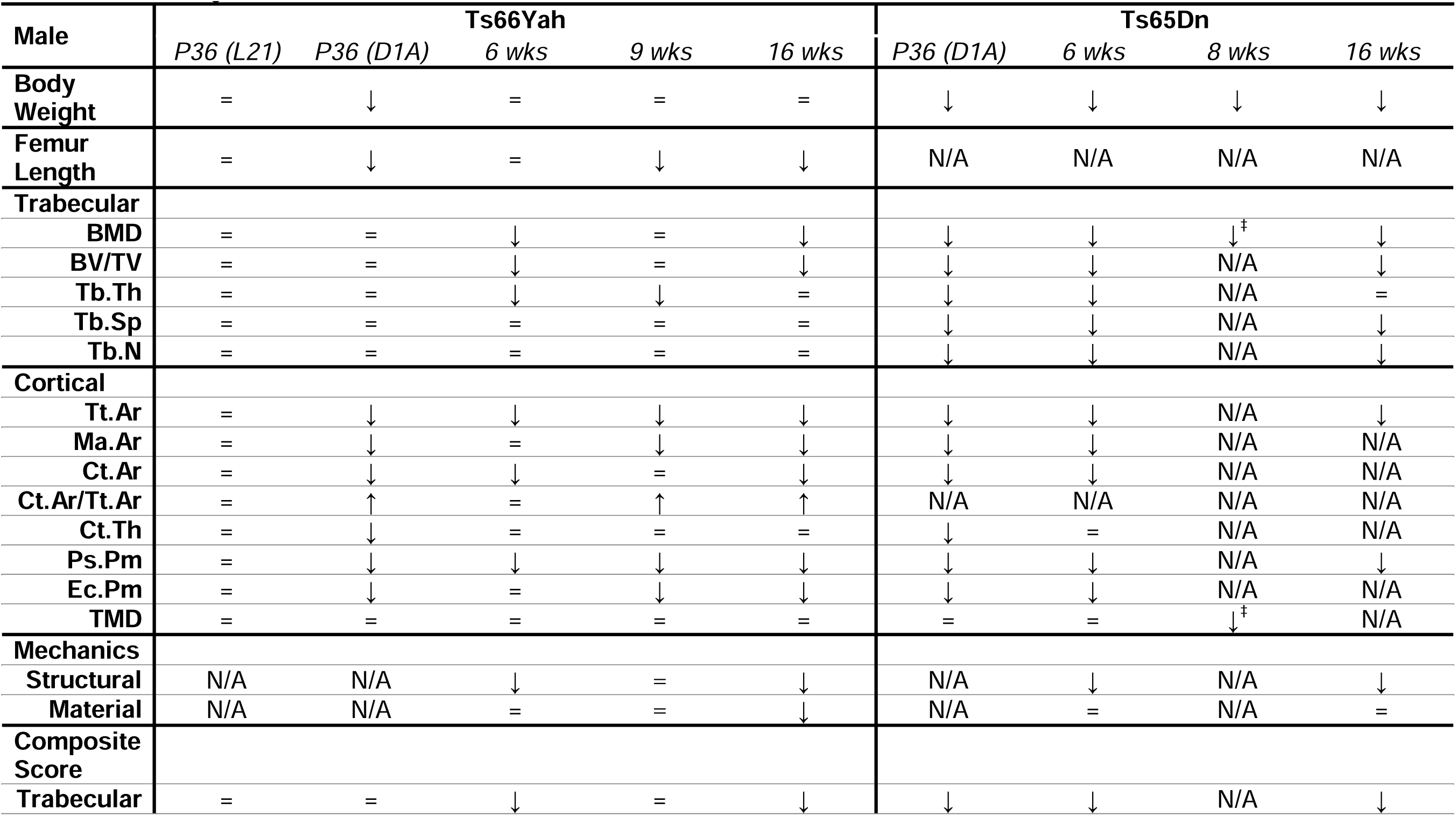
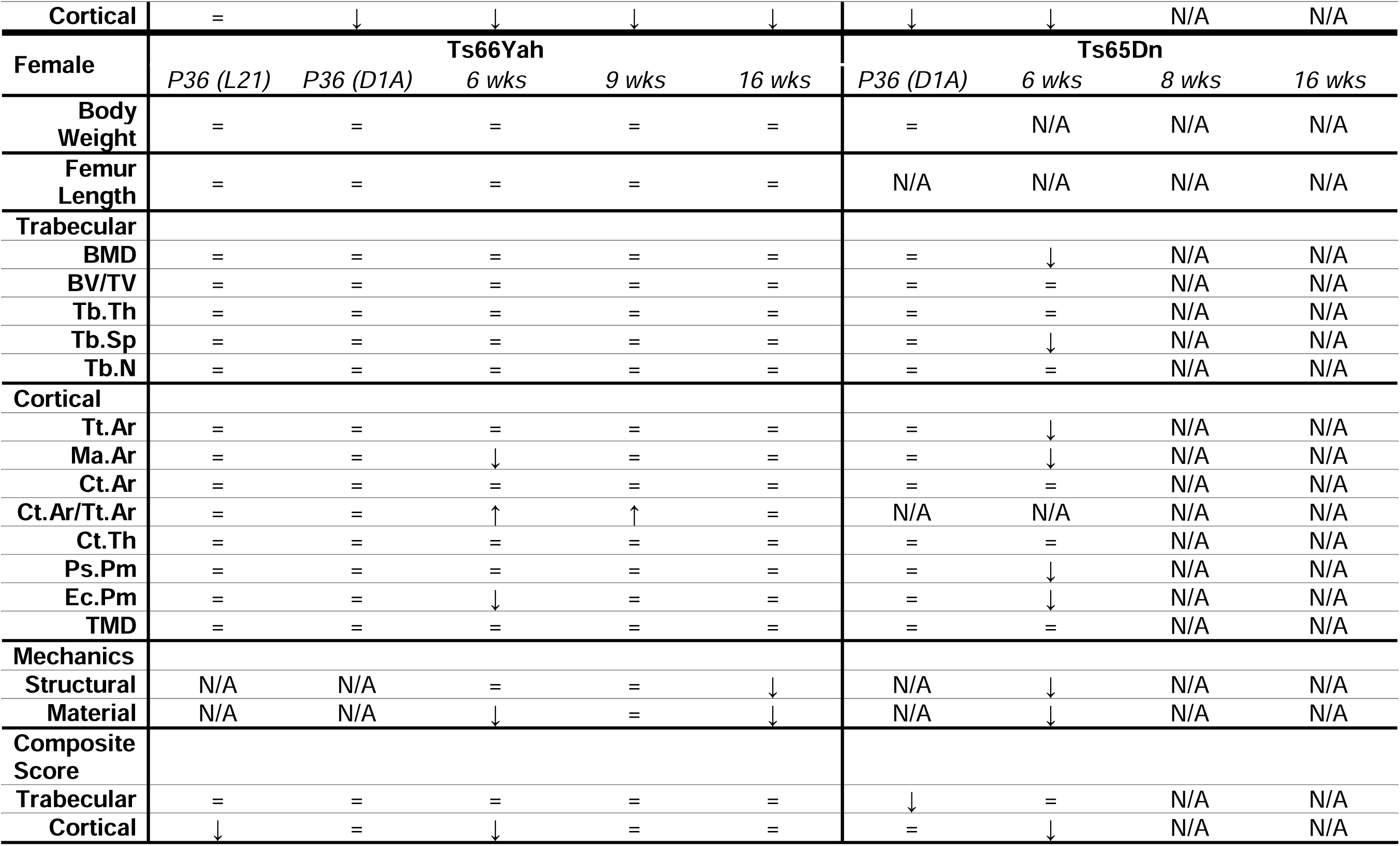
Summary of weight and femoral structural and mechanical phenotypes in Ts66Yah and Ts65Dn mice. ↓ indicates a significant deficit compared to euploid littermate mice, ↑ indicates a significant improvement compared to euploid littermate mice, *=* indicates no significant difference compared to euploid littermate mice, N/A indicates data is not available, ^‡^ indicates tibial data by dual x-ray absorptiometry & includes cortical bone. P36 (L21) interpretations utilize vehicle-treated euploid and Ts66Yah data while P36 (D1A) interpretation utilizes euploid and trisomic data from mice with two and three copies of *Dyrk1a*, respectively. All Ts66Yah and composite score interpretations made with data from this study. Ts65Dn interpretations came from LaCombe *et al*. 2024 for all P36 data, Thomas *et al*. 2021 for all 6-week bone data, Blazek *et al*. 2015a for 6-week weight data, Williams *et al*. 2018 for all 8-week data, Blazek *et al*. 2011 for all 16-week bone data, and Roper *et al*. 2006 for 16-week (4-month) weight data.

| Male | Ts66Yah |  |  |  |  | Ts65Dn |  |  |  |
| --- | --- | --- | --- | --- | --- | --- | --- | --- | --- |
|  | P36 (L21) | P36 (D1A) | 6 wks | 9 wks | 16 wks | P36 (D1A) | 6 wks | 8 wks | 16 wks |
| **Body Weight** | = | ↓ | = | = | = | ↓ | ↓ | ↓ | ↓ |
| **Femur Length** | = | ↓ | = | ↓ | ↓ | N/A | N/A | N/A | N/A |
| **Trabecular** |  |  |  |  |  |  |  |  |  |
| **BMD** | = | = | ↓ | = | ↓ | ↓ | ↓ | ↓‡ | ↓ |
| **BV/TV** | = | = | ↓ | = | ↓ | ↓ | ↓ | N/A | ↓ |
| **Tb.Th** | = | = | ↓ | ↓ | = | ↓ | ↓ | N/A | = |
| **Tb.Sp** | = | = | = | = | = | ↓ | ↓ | N/A | ↓ |
| **Tb.N** | = | = | = | = | = | ↓ | ↓ | N/A | ↓ |
| **Cortical** |  |  |  |  |  |  |  |  |  |
| **Tt.Ar** | = | ↓ | ↓ | ↓ | ↓ | ↓ | ↓ | N/A | ↓ |
| **Ma.Ar** | = | ↓ | = | ↓ | ↓ | ↓ | ↓ | N/A | N/A |
| **Ct.Ar** | = | ↓ | ↓ | = | ↓ | ↓ | ↓ | N/A | N/A |
| **Ct.Ar/Tt.Ar** | = | ↑ | = | ↑ | ↑ | N/A | N/A | N/A | N/A |
| **Ct.Th** | = | ↓ | = | = | = | ↓ | = | N/A | N/A |
| **Ps.Pm** | = | ↓ | ↓ | ↓ | ↓ | ↓ | ↓ | N/A | ↓ |
| **Ec.Pm** | = | ↓ | = | ↓ | ↓ | ↓ | ↓ | N/A | N/A |
| **TMD** | = | = | = | = | = | = | = | ↓‡ | N/A |
| **Mechanics** |  |  |  |  |  |  |  |  |  |
| **Structural** | N/A | N/A | ↓ | = | ↓ | N/A | ↓ | N/A | ↓ |
| **Material** | N/A | N/A | = | = | ↓ | N/A | = | N/A | = |
| **Composite Score** |  |  |  |  |  |  |  |  |  |
| **Trabecular** | = | = | ↓ | = | ↓ | ↓ | ↓ | N/A | ↓ |
| **Cortical** | = | ↓ | ↓ | ↓ | ↓ | ↓ | ↓ | N/A | N/A |
| **Female** | **Ts66Yah** |  |  |  |  | **Ts65Dn** |  |  |  |
|  | *P36 (L21)* | *P36 (D1A)* | *6 wks* | *9 wks* | *16 wks* | *P36 (D1A)* | *6 wks* | *8 wks* | *16 wks* |
| **Body Weight** | = | = | = | = | = | = | N/A | N/A | N/A |
| **Femur Length** | = | = | = | = | = | N/A | N/A | N/A | N/A |
| **Trabecular** |  |  |  |  |  |  |  |  |  |
| **BMD** | = | = | = | = | = | = | ↓ | N/A | N/A |
| **BV/TV** | = | = | = | = | = | = | = | N/A | N/A |
| **Tb.Th** | = | = | = | = | = | = | = | N/A | N/A |
| **Tb.Sp** | = | = | = | = | = | = | ↓ | N/A | N/A |
| **Tb.N** | = | = | = | = | = | = | = | N/A | N/A |
| **Cortical** |  |  |  |  |  |  |  |  |  |
| **Tt.Ar** | = | = | = | = | = | = | ↓ | N/A | N/A |
| **Ma.Ar** | = | = | ↓ | = | = | = | ↓ | N/A | N/A |
| **Ct.Ar** | = | = | = | = | = | = | = | N/A | N/A |
| **Ct.Ar/Tt.Ar** | = | = | ↑ | ↑ | = | N/A | N/A | N/A | N/A |
| **Ct.Th** | = | = | = | = | = | = | = | N/A | N/A |
| **Ps.Pm** | = | = | = | = | = | = | ↓ | N/A | N/A |
| **Ec.Pm** | = | = | ↓ | = | = | = | ↓ | N/A | N/A |
| **TMD** | = | = | = | = | = | = | = | N/A | N/A |
| **Mechanics** |  |  |  |  |  |  |  |  |  |
| **Structural** | N/A | N/A | = | = | ↓ | N/A | ↓ | N/A | N/A |
| **Material** | N/A | N/A | ↓ | = | ↓ | N/A | ↓ | N/A | N/A |
| **Composite Score** |  |  |  |  |  |  |  |  |  |
| **Trabecular** | = | = | = | = | = | ↓ | = | N/A | N/A |
| **Cortical** | ↓ | = | ↓ | = | = | = | ↓ | N/A | N/A |

Overall cortical bone structure of male and female Ts66Yah mice appear to be largely consistent with Ts65Dn mice at the examined ages, although not as many variables were affected in Ts66Yah femurs compared to Ts65Dn femurs (Table 3). Like Ts66Yah mice in the current study, male Ts65Dn mice exhibited cortical geometry deficits at P36, 6 weeks, and 16 weeks and female mice exhibited deficits only at 6 weeks (BLAZEK *et al*. 2011; THOMAS *et al*. 2021; LACOMBE *et al*. 2024). Again, it is unknown if female Ts65Dn mice have cortical geometry deficits at 16 weeks, but female Dp1Tyb mice display cortical geometry deficits at 16 weeks (THOMAS *et al*. 2020). While 16-week female Ts66Yah femurs lack cortical structure deficits present in 16-week female Dp1Tyb femurs, this may be the result of differing number of triplicated genes ∼100 vs. ∼145). The lack of cortical deficits at 16 weeks in female Ts66Yah mice may be transient and later ages could show the expected deficits.

Mechanics are often performed as a functional study of bone strength, as not all microarchitecture and geometry deficits or enhancements are sufficient to affect a bone’s susceptibility to fracture. Alterations of structural bone mechanics, which can be affected by bone size and cortical geometry, are found in male Ts65Dn (BLAZEK *et al*. 2011; THOMAS *et al*. 2021) and male Ts66Yah mice (this study) at 6 and 16 weeks (Table 3). However, like in cortical geometry, there were fewer variables altered in male Ts66Yah mice than male Ts65Dn mice. Female Ts66Yah mice displayed structural mechanical deficits at 16-weeks despite a lack of cortical geometry deficits, which together with the material property deficits, indicates the bone quality may be lower in female Ts66Yah mice around skeletal maturity. Future studies utilizing Raman spectroscopy could evaluate bone composition to confirm this. Material properties estimate the tissue level mechanics by normalizing structural bone mechanics using bone size, so alterations in material properties may not be present even if structural bone mechanics are altered. Alterations in material properties were found in 6-week-old female Ts65Dn and Ts66Yah mice. Although material properties have not been evaluated in 16-week-old Ts65Dn mice, both male and female Ts66Yah and Dp1Tyb mice show alterations in material properties. Taken together, these data illustrate that alterations in mechanics are generally conserved between male Ts66Yah and Ts65Dn mouse models of DS, although the severity is lessened in Ts66Yah mice.

### Comparisons of skeletal deficits in Ts66Yah and Ts65Dn DS mouse models

Ts66Yah mice had no significant differences in body weight after P36, however low body weight in Ts65Dn mice compared to euploid littermate mice has been observed across development into male adulthood (Table 3) (ROPER *et al*. 2006). Mutations in a trisomic Mmu17 gene in Ts65Dn mice, *ARID1B,* generally result in low body weight with normal body mass index distribution in humans and may contribute to low body weight in Ts65Dn mice (LIU *et al*. 2020). Some reports on the loss of this phenotype in Ts65Dn mice during postnatal development has been attributed to strain differences (Jackson laboratory #001924 vs. #005252) (SHAW *et al*. 2020) but could also be affected by genetic drift, differences in genetic background, or allelic heterogeneity (DEITZ AND ROPER 2011). Despite the lack of body weight differences in Ts66Yah mice, femur length was still decreased in male Ts66Yah mice at 9 and 16 weeks, which has also been reported in Ts65Dn mice postnatally but not prenatally (OLSON *et al*. 2004; BLAZEK *et al*. 2015b). This suggests that while body weight and femoral length may be correlated in DS mice, shortened long bones can be present without a difference in body weight.

Comparisons of structural bone variable values between different mouse models of DS can be difficult due to conflated background strain effects (DEITZ AND ROPER 2011; PAPAGEORGIOU *et al*. 2020; FÖGER-SAMWALD *et al*. 2021) or methodological differences in measuring bone microstructure (e.g. using different µCT machines or different analyzers) (BONNET *et al*. 2009; VERDELIS *et al*. 2011). By standardizing each animal’s value using the mean and standard deviation of all mice within each mouse model, these effects are minimized so direct comparisons can be made. Additionally, multiple variables with varying degrees of correlation are obtained from each bone compartment of every animal and effects of trisomy may vary (SLOAN *et al*. 2023). This multivariate problem was addressed by using linear combination of the standardized variables and principal component 1 loadings of each variable derived from PCA to yield a new (uncorrelated) score. After comparing these scores between mouse models, we have determined that structural phenotypes are not significantly different in male and female Ts65Dn and Ts66Yah femurs at 6 weeks, male femurs at 16 weeks, or male and female cortical bone at P36 (Table 3). However, both sexes of Ts66Yah mice lacked deficits in the trabecular bone composite score, unlike Ts65Dn mice, suggesting trisomic, non-Hsa21 orthologous genes in Ts65Dn may result in earlier trabecular microarchitecture deficits during bone accrual either directly or through interactions with trisomic, Hsa21 orthologous genes or disomic genes.

This multivariate approach has also resulted in differing conclusions than interpreting variables individually. Female Ts65Dn mice were suggested to have trabecular deficits at 6 weeks due to low BMD and high Tb.Sp (THOMAS *et al*. 2021), but these deficits were not sufficient to produce a significant deficit in trabecular composite score (Table 3). The trabecular composite score findings suggest female Ts65Dn mice do not have a gross trabecular deficit at 6 weeks, which is consistent with trabecular phenotypes of other female DS mouse models (THOMAS *et al*. 2020; LAMANTIA *et al*. 2024). Additionally, female Ts65Dn mice (LACOMBE *et al*. 2024) had no deficits in any individual trabecular variable at P36, but there is a decrease in trabecular composite score (Table 3). This was also found in female Ts66Yah mice given 0.5% carboxymethylcellulose by oral gavage, where these mice had no alterations in any individual cortical variables at P36 yet had a significantly decreased cortical composite score (Table 3). Together, this suggests the composite score method has the additional use of detecting low severity or subclinical phenotypes, which can aid in determining treatment timepoints.

Statistically comparing DS mouse models across studies can be done in the future by utilizing the new described method with composite scores obtained from PCA. Although it is currently used in the context of DS and single gene effects on structural phenotypes in the skeleton, the method may be applied to other phenotypic tests that pose a multivariate problem, having multiple dependent variables with varying degrees of correlation. Outside of the DS research community, this method can also be used for other µCT studies as an additional measure to objectively confirm overall deficits in trabecular and cortical regions, since it is uncommon for every variable to be affected. Utilizing PCA to create a single composite variable will be most useful when the phenotypic variables are highly correlated and the first component is sufficient in explaining a large portion of the variation (SONG *et al*. 2013). Otherwise, combinations of more than one component may be needed to fully capture the information embedded in the phenotypic data.

### Effects of normalization of Dyrk1a copy number differs between male Ts65Dn and Ts66Yah mice

Reducing *Dyrk1a* copy number in Ts66Yah animals (Ts66Yah,*Dyrk1a*^+/+/-^) did not improve trabecular bone microarchitecture at P36. There were also no significant trabecular deficits in male or female Ts66Yah mice with three copies of *Dyrk1a* compared to euploid animals with two copies of *Dyrk1a* at this age, which suggests trisomic *Dyrk1a* is not a major contributor to Ts66Yah trabecular structure at P36. Cortical bone size was significantly decreased in male Ts66Yah mice compared to euploid mice and significantly improved by *Dyrk1a* normalization at P36 but was not completely rescued to euploid levels. Previous findings in male Ts65Dn animals found that normalization of *Dyrk1a* copy number by the same means as the current study resulted in improvement in all trabecular variables and Ct.Th at P36 (LACOMBE *et al*. 2024). Although one aspect of cortical structure was improved in Ts65Dn mice, Ts66Yah mice benefitted more from *Dyrk1a* copy normalization in the cortical region at this age. Together, this indicates genetic interactions vary between mouse models. It may be that non-Hsa21 orthologous, trisomic genes in male Ts65Dn mice work synergistically with *Dyrk1a* to result in earlier trabecular microarchitecture deficits, which are amenable to *Dyrk1a* normalization, but contribute more to cortical growth delay. For example, *Arid1b*, a gene located in the trisomic Mmu17 centromeric region of Ts65Dn mice, was associated with growth delay by repressing canonical Wnt signaling (like DYRK1A) (VASILEIOU *et al*. 2015; LIU *et al*. 2020), and conditions related to *ARID1B* have noted delayed bone growth and bone age (VERGANO *et al*. 1993; TAN *et al*. 2022; TAO *et al*. 2022). *Dyrk1a* copy number normalization may improve trabecular deficits at a later age in Ts66Yah mice, as our previous study showed persistent trabecular deficits preceded trisomic *Dyrk1a*’s involvement in Ts65Dn mice (LACOMBE *et al*. 2024).

### Pharmacological DYRK1A inhibition and confounding factors

Leucettinib-21 treatment may negatively affect male mouse growth as evidenced by significantly less weight gain between the beginning and end of treatment in euploid and Ts66Yah mice. Despite these effects on growth that affected both genotypes, only male L21-treated Ts66Yah mice appeared to have trabecular and cortical structural deficits and femoral length deficits. While this could be due to the dosage-sensitive nature of *Dyrk1a*, these results may be confounded by an unexpected initial weight difference between male Ts66Yah vehicle-and L21-treated mice at P21. Body weight differences often suggest lower bone mass, and body weight was found to correlate with femur length, so it is possible that the L21-treated male Ts66Yah group had a more severe bone phenotype than the vehicle-treated Ts66Yah group before treatment was administered. This makes it difficult to assess if the difference found in the femurs at P36 was due to Leucettinib-21 treatment or the innate differences between the groups. This may be due to uncontrolled differences present during litter selection and treatment assignment or to potential variability in the phenotype penetrance in Ts66Yah mice, which has been observed in individuals with DS.

There was also seemingly no difference in cortical bone between vehicle-treated male euploid and Ts66Yah mice, which contrasts with the results of the *Dyrk1a* reduction experiment at the same age. This could potentially be due to epigenetic factors or a result of the vehicle intubation treatment itself. The vehicle used in this study was 0.5% carboxymethylcellulose (CMC), which has been found to decrease osteoclastogenesis and increase osteoblastogenesis *in vitro* (AGIS *et al*. 2010; QI *et al*. 2018). It is possible treatment with CMC, either through known osteogenic properties or other epigenetic avenues, alleviates the cortical deficits in Ts66Yah mice. Nonetheless, the unexpected results of the vehicle-treated animals make it difficult to assess L21’s potential as a treatment in this study. Leucettinib cognitive treatment studies in DS mouse models are reviewed elsewhere (MEIJER *et al*. 2024). Leucettinib-21 has been used in mouse models of DS with no noted adverse effects despite the difference in age (embryos vs. adults) and tissue investigated (heart vs. brain) (LINDBERG *et al*. 2023; LANA-ELOLA *et al*. 2024). However, this study was the first to our knowledge to directly treat with Leucettinib-21 during a period of rapid postnatal development. The previously treated embryos may have been protected from potentially adverse effects on growth due to necessity of crossing the placental barrier, and adult mice are not growing nearly as fast. These data illustrate the importance of timing the use of DYRK1A inhibitors for when *Dyrk1a* is overexpressed as implicated in previous studies (STRINGER *et al*. 2017b; LACOMBE *et al*. 2024). There also needs to be consideration the impact of DYRK1A inhibition on other non-target tissues. DYRK1A overexpression varies based on tissue and age (HAWLEY *et al*. 2023; LACOMBE *et al*. 2024), which means treating cognitive deficits could worsen bone, as evidenced by previous EGCG treatments (JAMAL *et al*. 2022). This is not to say that studies of DYRK1A inhibition treatments for tissue-specific deficits should not go forward; rather multiple tissues should be examined when treatments are tested to ensure deterioration of one tissue is not at the expense of a modest improvement of another.

### Periodic normalization of skeletal deficits in DS

At 9 weeks of age, male Ts66Yah mice had almost no trabecular deficits, in contrast to those seen at 6 and 16 weeks, illustrating a periodic developmental normalization of trabecular deficits. This has also been observed in male (P24 and P27) and female (P36) Ts65Dn mice in trabecular and cortical bone compartments (LACOMBE *et al*. 2024). A similar phenomenon has also been noted in boys (11 to nearly 18 years old) and girls (10-16.5 years old) with DS in the cortical bone compartment, which was then termed a “bone recovery phase” that covered 2.5 years (GARN *et al*. 1972). Anthropometric and radiologic studies have also noted atypical non-linear growth and skeletal maturation rates in children and adolescents with DS, suggesting there is a period of time during childhood and adolescence (around 8-15 years of age) when children/adolescents with DS “catch-up” to and eventually surpass their chronological age (POZSONYI *et al*. 1964; RARICK *et al*. 1964; ROCHE 1967; RUNDLE *et al*. 1972).

Taken together, these data suggest that periodic normalization of skeletal deficits in multiple mouse models of DS is representative of individuals with DS. However, there is not yet a consensus on the timing of the periodic normalizations in each mouse model, which may be due to differences in the number of triplicated genes, which genes are triplicated, the transmission type of triplicated genes, or lack of multiple timepoints with available data. More studies need to be conducted in both DS mouse models and individuals with DS to understand when and why these periodic normalizations occur. New therapies may be developed from understanding the mechanism behind these normalizations, and critical times for treatments can be better understood.

### Limitations

Although the goal of this study was to compare structural phenotypes of the femur between Ts66Yah and Ts65Dn mice, further characterization of the skeleton should be conducted to determine the phenotypic similarity between Ts66Yah mice and individuals with DS. Such studies should include additional sites, such as the lumbar vertebrae, and evaluation of the cellular mechanisms behind the structural phenotypes using serum biomarkers and histomorphometry.

### Conclusions

The trisomic, non-Hsa21 orthologous *Scaf8*-*Pde10a* region in Ts65Dn mice appears to modulate an aspect of appendicular structural phenotypes by producing earlier trabecular deficits and more severe trabecular and cortical deficits in both sexes of mice. Additionally, the effect of *Dyrk1a* normalization, a highly investigated target for treatment in humans with DS, varies between Ts66Yah and Ts65Dn mice. These differences between mouse models illustrate the importance of confirming genetic and cellular mechanisms in multiple mouse models of DS, then in individuals with DS. Structural and mechanical evaluation of Ts66Yah mice indicate this is a potential model for DS-related appendicular skeletal phenotypes, but further evaluation of bone cellular dynamics and other skeletal sites should also be considered. The composite score method derived from PCA allows for direct comparisons between differing datasets, a simplified yet comprehensive overview of phenotypes involving multiple variables, and the possible detection of subclinical phenotypes.

## Data Availability

Data has been uploaded to Dryad (DOI: 10.5061/dryad.kkwh70sf1). Reviewer URL: http://datadryad.org/stash/share/Wc0UyGfRfLvEzlF_FsYoCz7v4KjG9Gn2-y4F3hgutD8.

## Acknowledgements

We would like to thank Faith Prochaska, Isabella Crawford, Rebekah Hurd, and Joshua Lamantia for acquiring µCT scans, Dr. Laurent Meijer for providing the Leucettinib-21 and treatment protocol, and Claire Chevalier of Dr. Yann Herault’s group for preparing Ts66Yah breeders for the IU Indianapolis colony. We would also like to acknowledge the Small Animal Skeletal Phenotyping Core at the IU School of Medicine for the use of the SkyScan 1172.

## CRediT Authorship Contribution Statement

**Kourtney Sloan:** Conceptualization, Methodology, Formal analysis, Investigation, Visualization, Supervision, Project administration, Writing – Original Draft

**Kristina M. Piner:** Formal analysis, Investigation, Writing – Review & Editing

**Pathum Randunu Nawarathna Kandedura Arachchige:** Conceptualization, Methodology, Formal analysis, Visualization, Writing – Review & Editing

**Charles R. Goodlett**: Conceptualization, Methodology, Supervision, Writing – Review & Editing

**Yann Herault:** Resources, Writing – Review & Editing

**Gayla R. Olbricht:** Conceptualization, Methodology, Supervision, Writing – Review & Editing

**Joseph M. Wallace:** Methodology, Software, Resources, Writing – Review & Editing

**Randall J. Roper:** Conceptualization, Resources, Funding acquisition, Supervision, Project administration, Writing – Original Draft

## Supplemental Tables and Figures

**Supplemental Figure 1:**
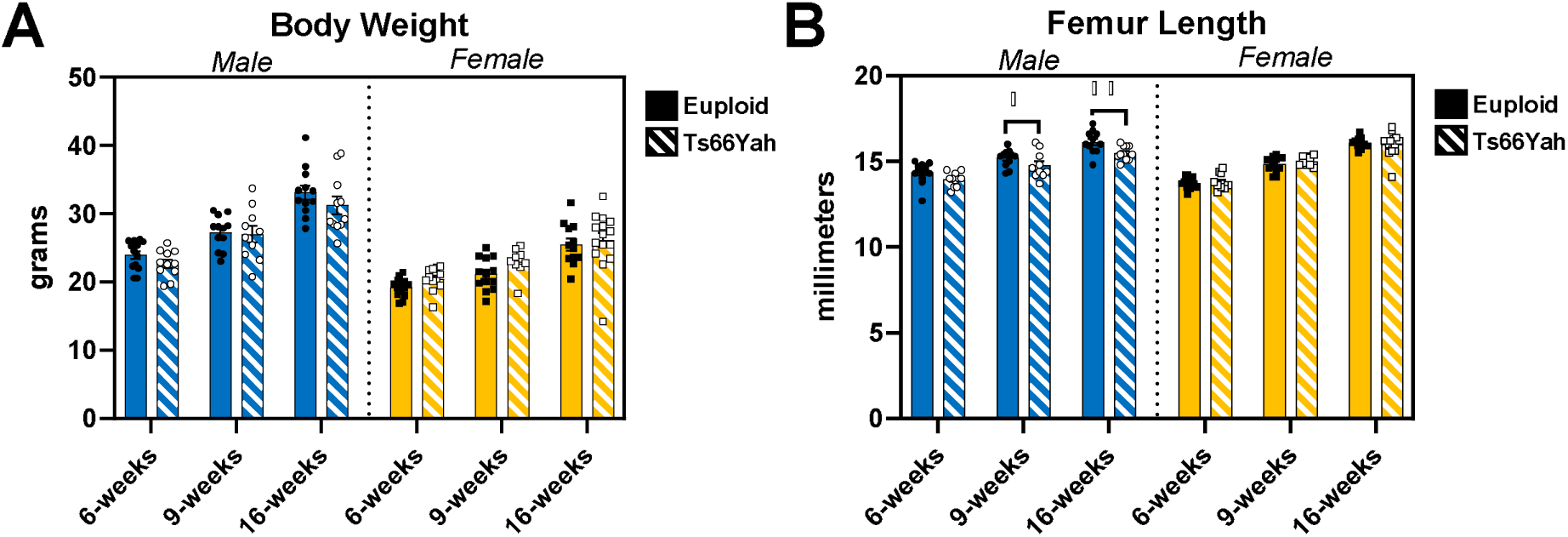
Body weight and femur lengths of 6-, 9-, and 16-week-old Ts66Yah mice. Data are mean ± SEM. Asterisks indicate a significant difference between groups in pairwise comparisons with Sidak correction. * *p* < 0.05, ** *p* < 0.01. **B)** Pairwise comparisons between ages within genotype and sex for femur length: significantly increased between each age in all four groups. 6 weeks: male euploid (n = 13), male Ts66Yah (n = 11), female euploid (n = 15 [body weight] or 13 [femur length]), female Ts66Yah (n = 12 [body weight] or 11 [femur length]); 9 weeks: male euploid (n = 12), male Ts66Yah (n = 10), female euploid (n = 13), female Ts66Yah (n = 9); 16 weeks: male euploid (n = 12 [body weight] or 11 [femur length]), male Ts66Yah (n = 11), female euploid (n = 12), female Ts66Yah (n = 15).

**Supplemental Figure 2:**
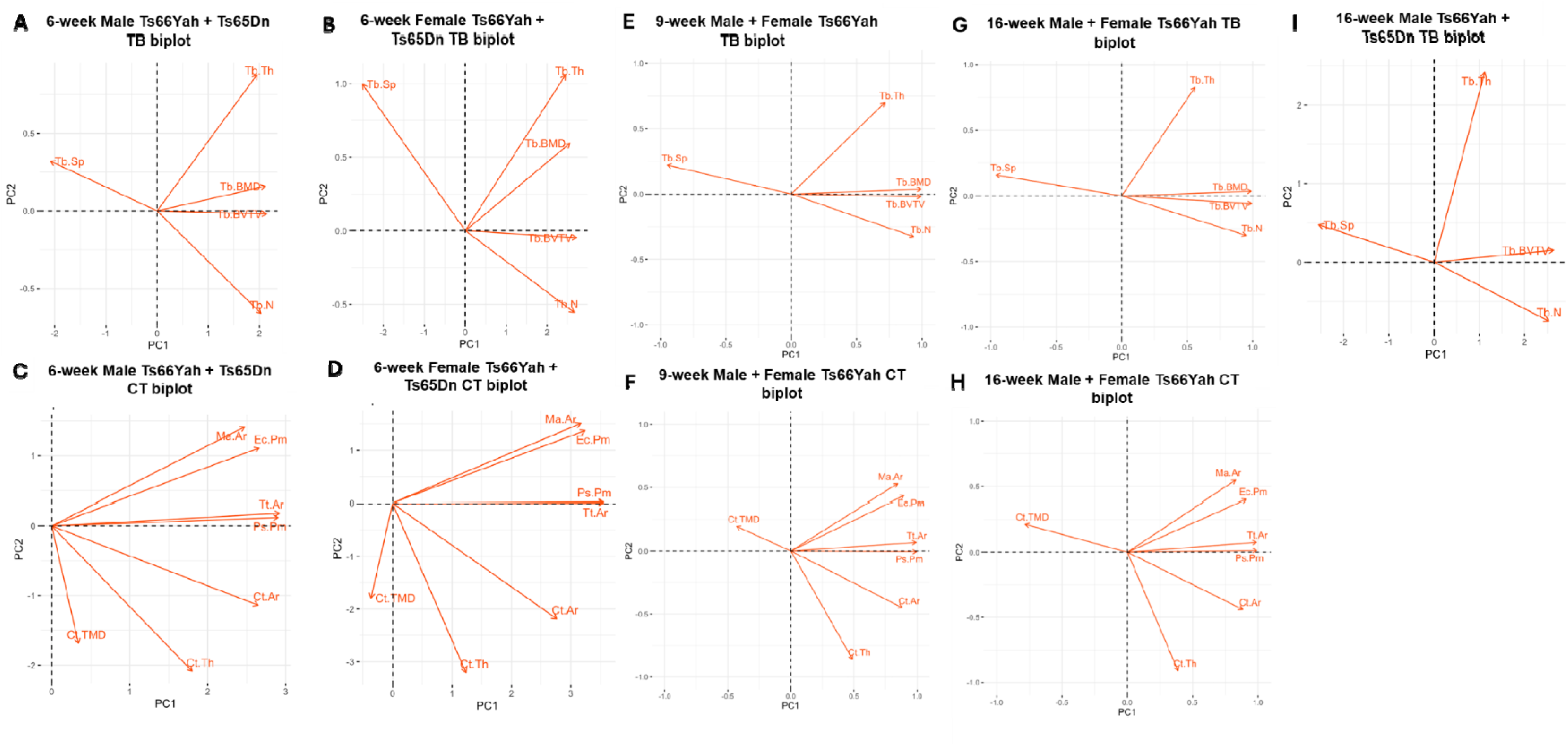
Principal component analysis biplots for trabecular (TB) and cortical (CT) variables of Ts65Dn and Ts66Yah mice. **A-D)** Biplots generated separately for male (A,C) and female (B,D) trabecular (A,B) and cortical (C,D) variables using 6-week male and female Ts65Dn data from euploid littermates and Ts65Dn mice lacking OSX-cre (THOMAS *et al*. 2021) and 6-week male and female Ts66Yah data (this study). **E-F)** Biplots generated separately for trabecular (E) and cortical (F) variables using 9-week male and female Ts66Yah data (this study). **G-H)** Biplots generated separately for trabecular (G) and cortical (H) variables using 16-week male and female Ts66Yah data (this study. **I)** Biplot generated for male trabecular variables using 16-week male Ts65Dn data (BLAZEK *et al*. 2011) and 16-week male Ts66Yah data (this study). See Supplemental Table 3 for PCA results.

**Supplemental Figure 3:**
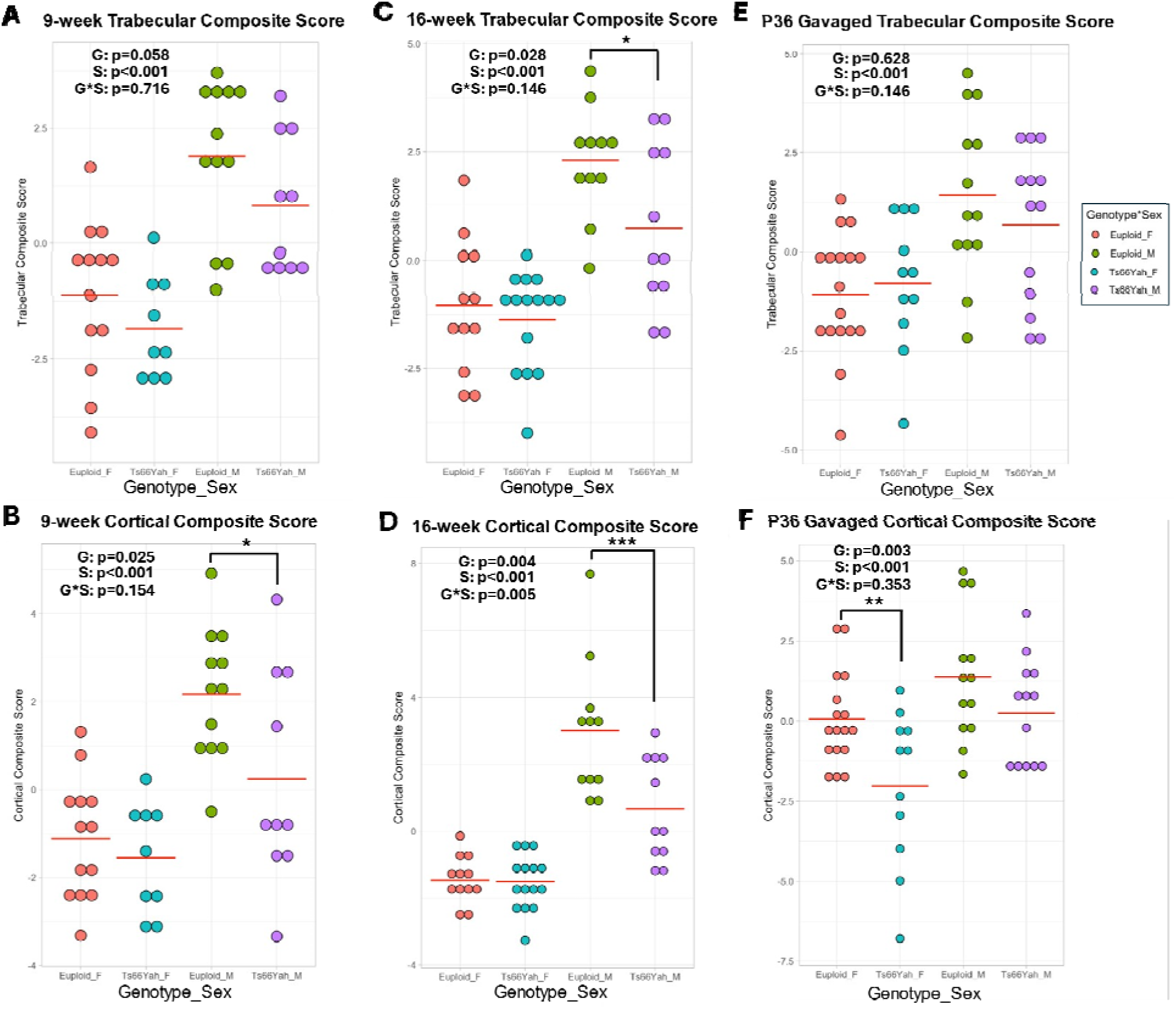
Trabecular and cortical composite scores for male and female Ts66Yah at 9 weeks (A-B), 16 weeks (C-D), and P36 vehicle-treated mice (E-F). Red horizontal line indicates group mean. Two-way ANOVA with genotype (G) and sex (S) as between subject factors. Asterisks indicate significant genotype difference in sex-stratified contrast analysis. \**p* < 0.05, ** *p* < 0.01, *** *p* < 0.001.

**Supplemental Figure 4:**
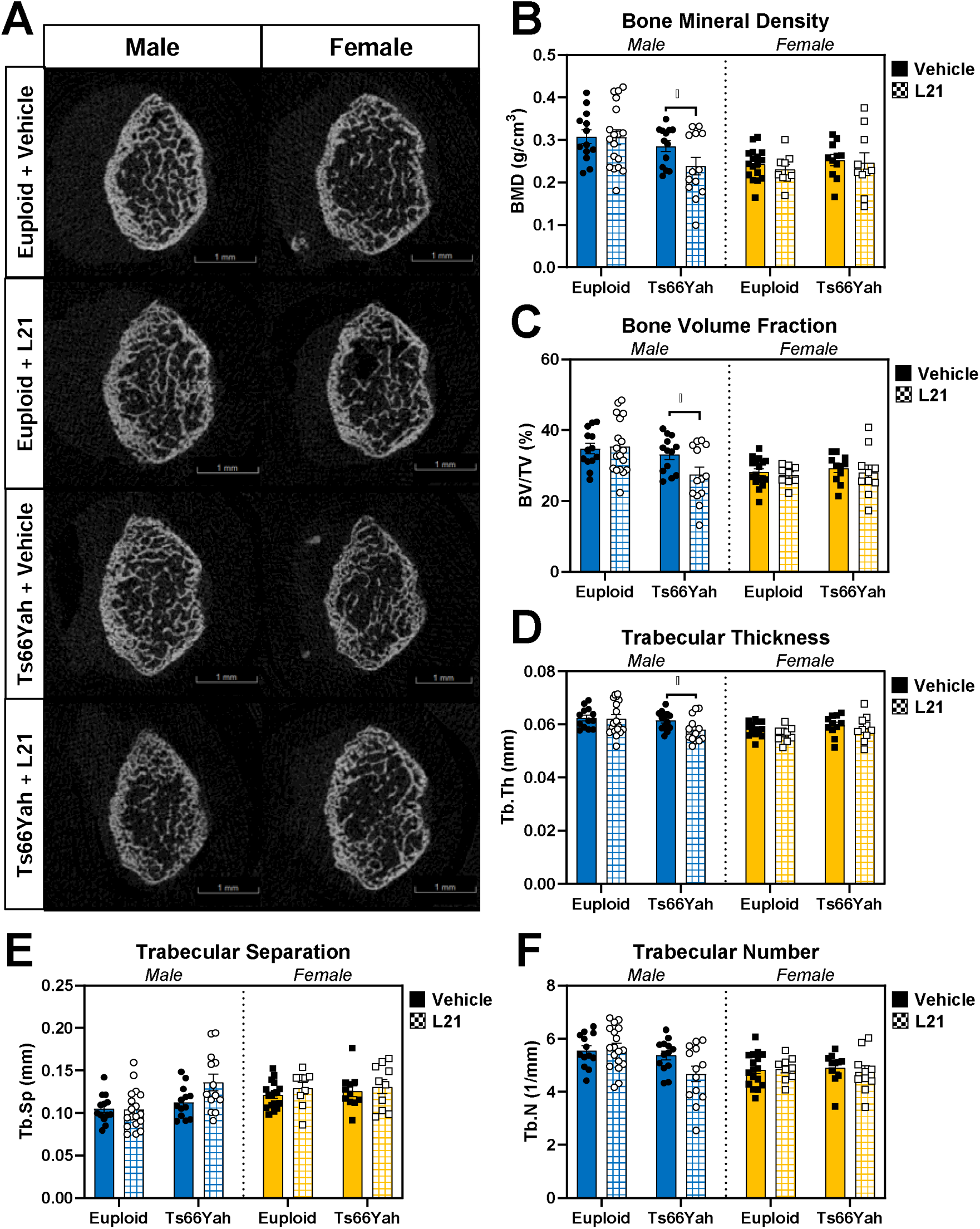
Trabecular bone variables for Leucettinib-21-treated Ts66Yah mice. **A)** Representative images of trabecular bone taken halfway way through the 1mm trabecular region as determined by finding the animal with the closest average distance away from the mean of each trabecular variable. **B-F)** Data are mean ± SEM. Asterisks indicate a significant difference between groups in pairwise comparisons with Sidak correction. * *p* < 0.05. Male mice: vehicle-treated euploid (n = 13), L21-treated euploid (n = 18), vehicle-treated Ts66Yah (n = 13), L21-treated Ts66Yah (n = 13). Female mice: vehicle-treated euploid (n = 17), L21-treated euploid (n = 9), vehicle-treated Ts66Yah (n = 11), L21-treated Ts66Yah (n = 10).

**Supplemental Figure 5:**
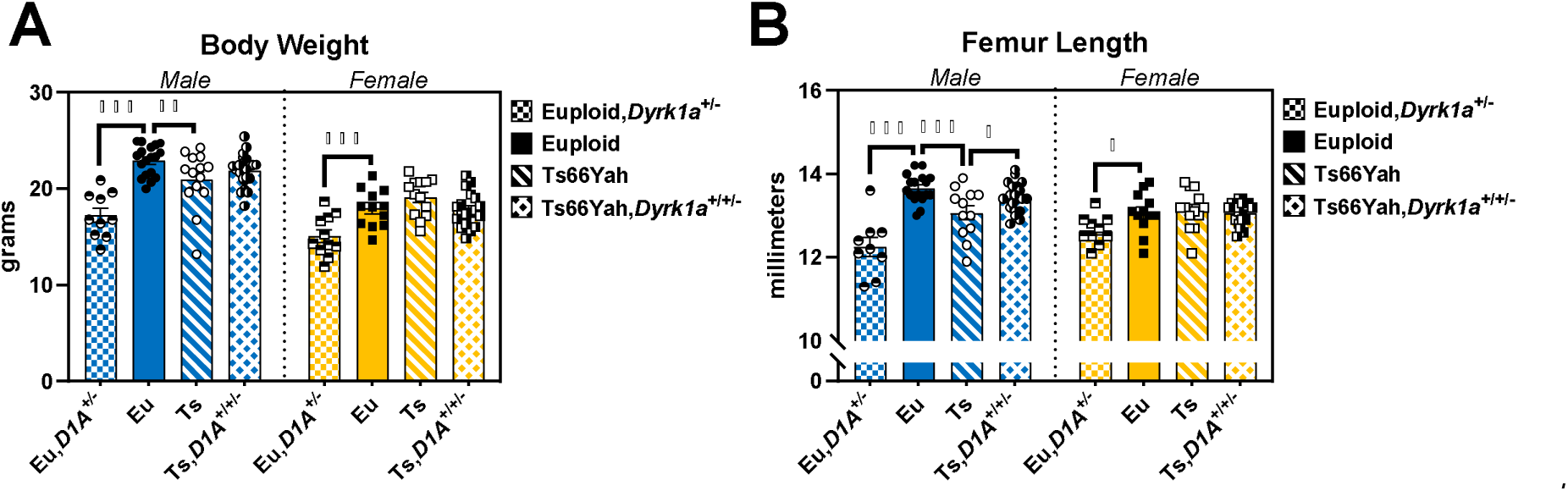
Body weight and femur lengths of postnatal day (P)36 Ts66Yah,*Dyrk1a*^+/+/-^mice. Data are mean ± SEM. Asterisks indicate a significant difference between groups in pairwise comparisons with Sidak correction. * *p* < 0.05, ** *p* < 0.01, *** *p* < 0.001. Male mice: euploid (n = 18 [body weight] or 17 [femur length]); euploid,*Dyrk1a*^+/-^ (n = 10 [body weight] or 9 [femur length]); Ts66Yah (n = 14 [body weight] or 12 [femur length]); Ts66Yah,*Dyrk1a*^+/+/-^ (n = 22 [body weight] or 21 [femur length]). Female mice: euploid (n = 12); euploid,*Dyrk1a*^+/-^ (n = 11 [body weight] or 10 [femur length]); Ts66Yah (n = 13); Ts66ah,*Dyrk1a*^+/+/-^ (n = 19).

**Supplemental Figure 6:**
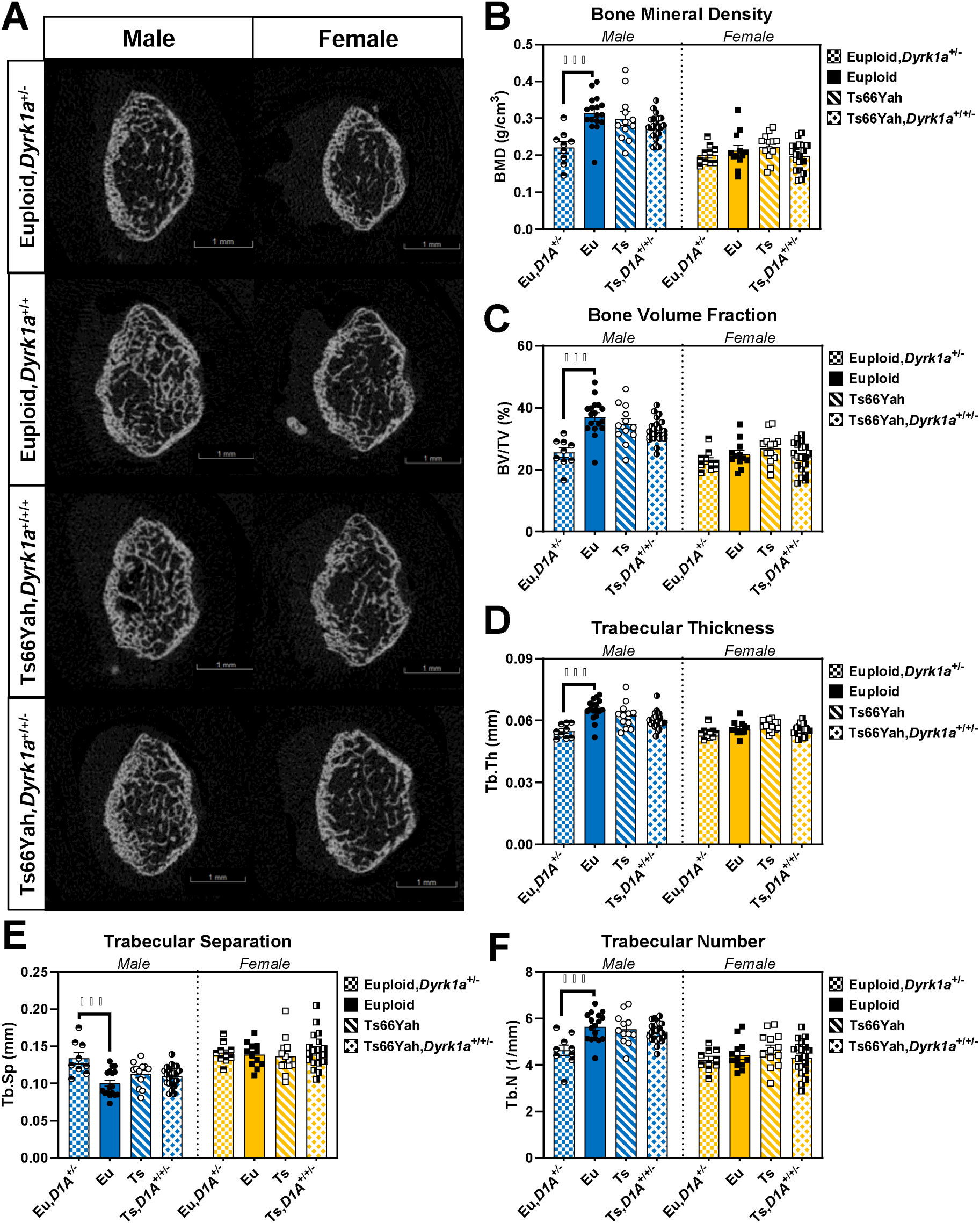
Trabecular bone variables for P36 Ts66Yah,*Dyrk1a^+/+/-^* mice. **A)** Representative images of trabecular bone taken halfway through the 1mm trabecular region as determined by finding the animal with the closest average distance away from the mean of each trabecular variable. **B-F)** Data are mean ± SEM. Asterisks indicate a significant difference between groups in pairwise comparisons with Sidak correction. *** *p* < 0.001. Male mice: euploid (n = 17); euploid,*Dyrk1a*^+/-^ (n = 9); Ts66Yah (n = 12); Ts66Yah,*Dyrk1a*^+/+/-^ (n = 21). Female mice: euploid (n = 12); euploid,*Dyrk1a*^+/-^ (n = 10); Ts66Yah (n = 13); Ts66ah,*Dyrk1a*^+/+/-^ (n = 19).

**Supplemental Figure 7:**
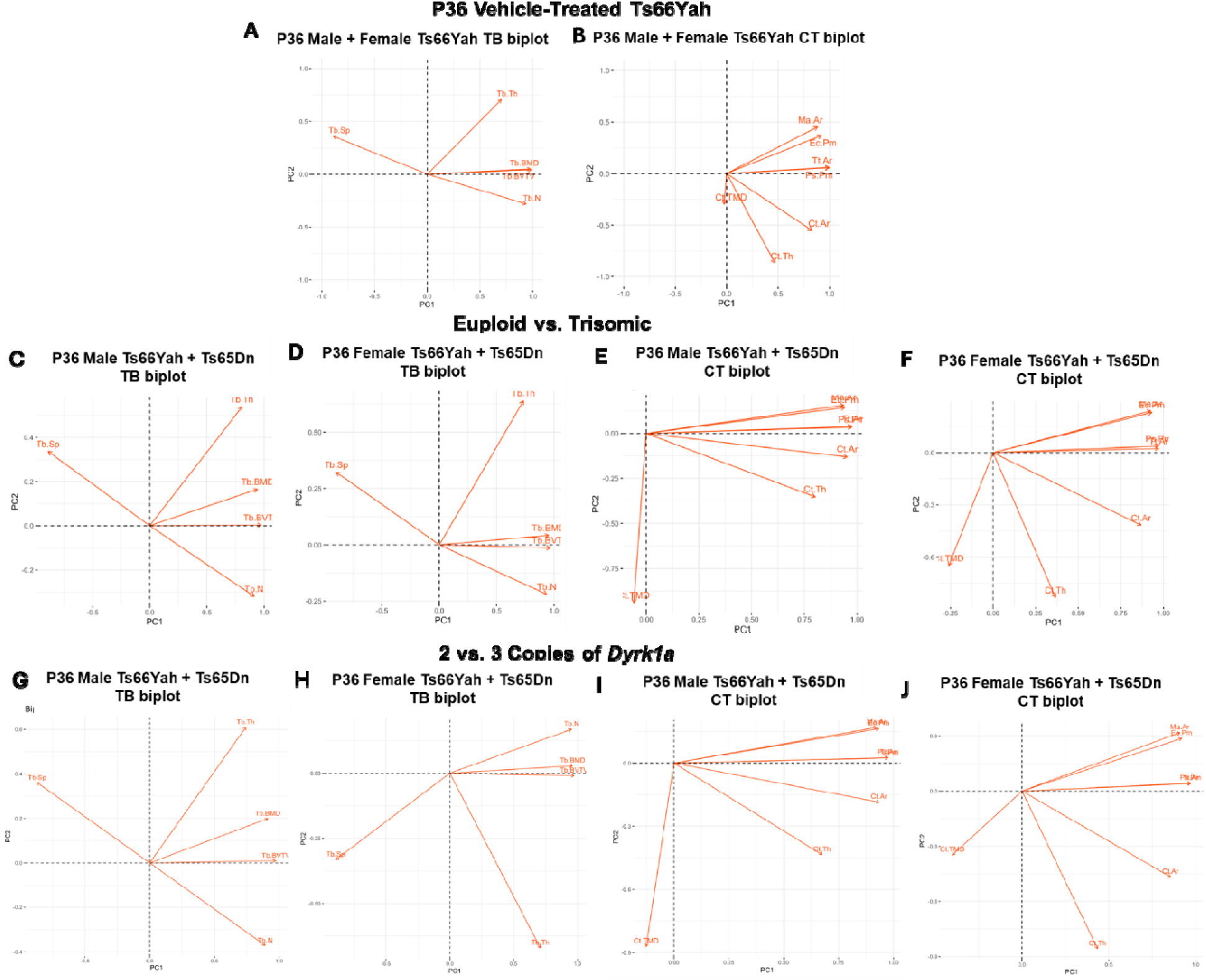
Principal component analysis biplots for trabecular (TB) and cortical (CT) variables of Ts65Dn, Ts66Yah, and germline reduction of *Dyrk1a* copy number mice. **A-B)** Biplots generated separately for trabecular (A) and cortical (B) variables using postnatal day (P)36 male and female vehicle-treated Ts66Yah mice (this study). **C-F)** Biplots generated separately for male (C,E) and female (D,F) trabecular (C,D) and cortical (E,F) variables using P36 Ts65Dn,*Dyrk1a^+/+/+^*and euploid,*Dyrk1a^+/+^* data from (LACOMBE *et al*. 2024) and P36 Ts66Yah,*Dyrk1a*^+/+/+^ and euploid,*Dyrk1a*^+/+^ data from this study. **G-J)** Biplots generated separately for male (G,I) and female (H,J) trabecular (G,H) and cortical (I,J) variables using P36 Ts65Dn,*Dyrk1a^+/+/+^* and Ts65Dn,*Dyrk1a*^+/+/-^ data from (LACOMBE *et al*. 2024) and P36 Ts66Yah,*Dyrk1a*^+/+/+^ and Ts66Yah,*Dyrk1a*^+/+/-^ data from this study. See Supplemental Table 4 for PCA results.

**Supplemental Table 1:**
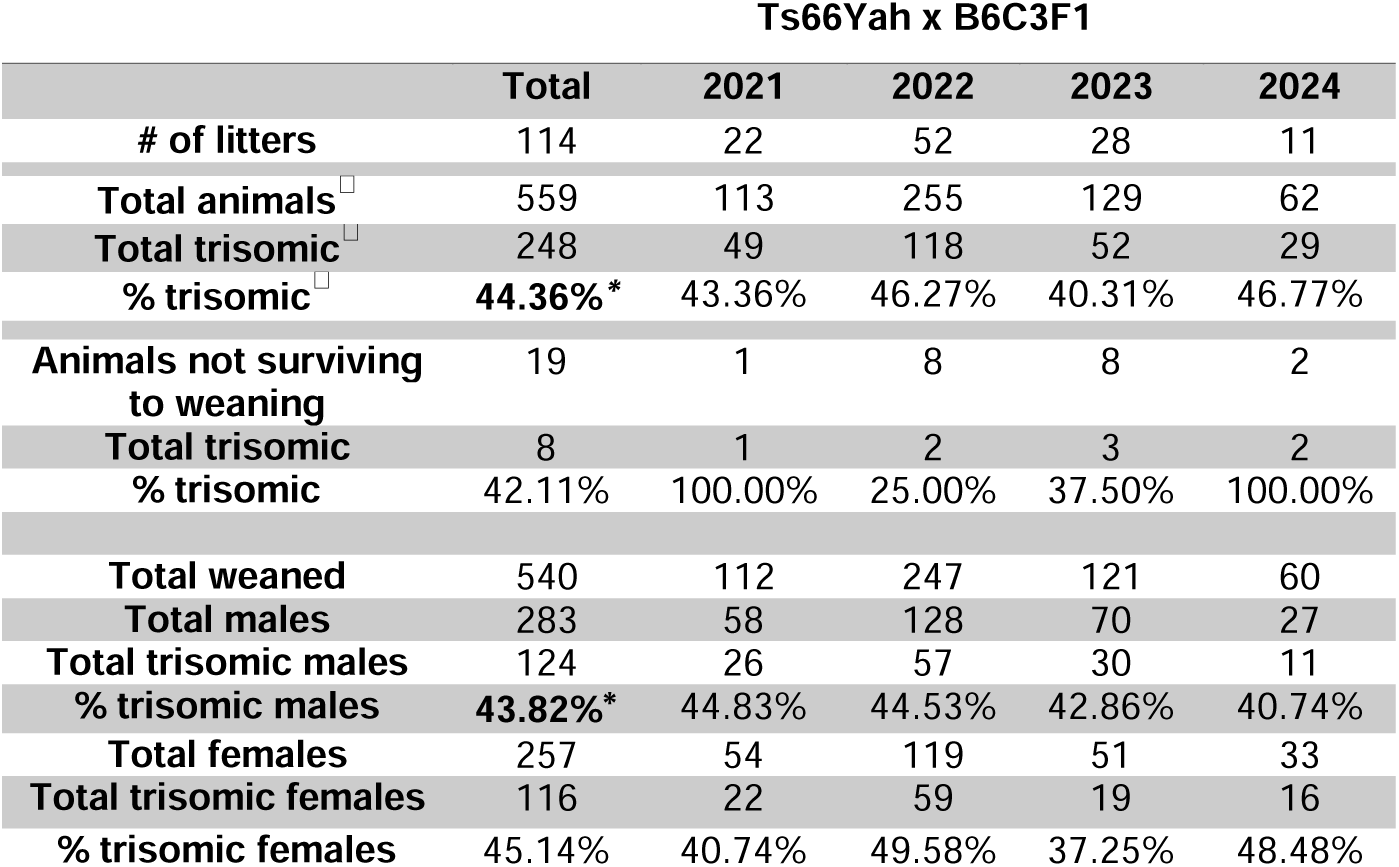
Transmission of trisomy in Ts66Yah x B6C3F1 breeding scheme. (^□^) to represent that animals that did not survive to weaning were included. **\* indicates *p* < 0.05** based on chi-square goodness of fit test performed on total data.

**Supplemental Table 2:**
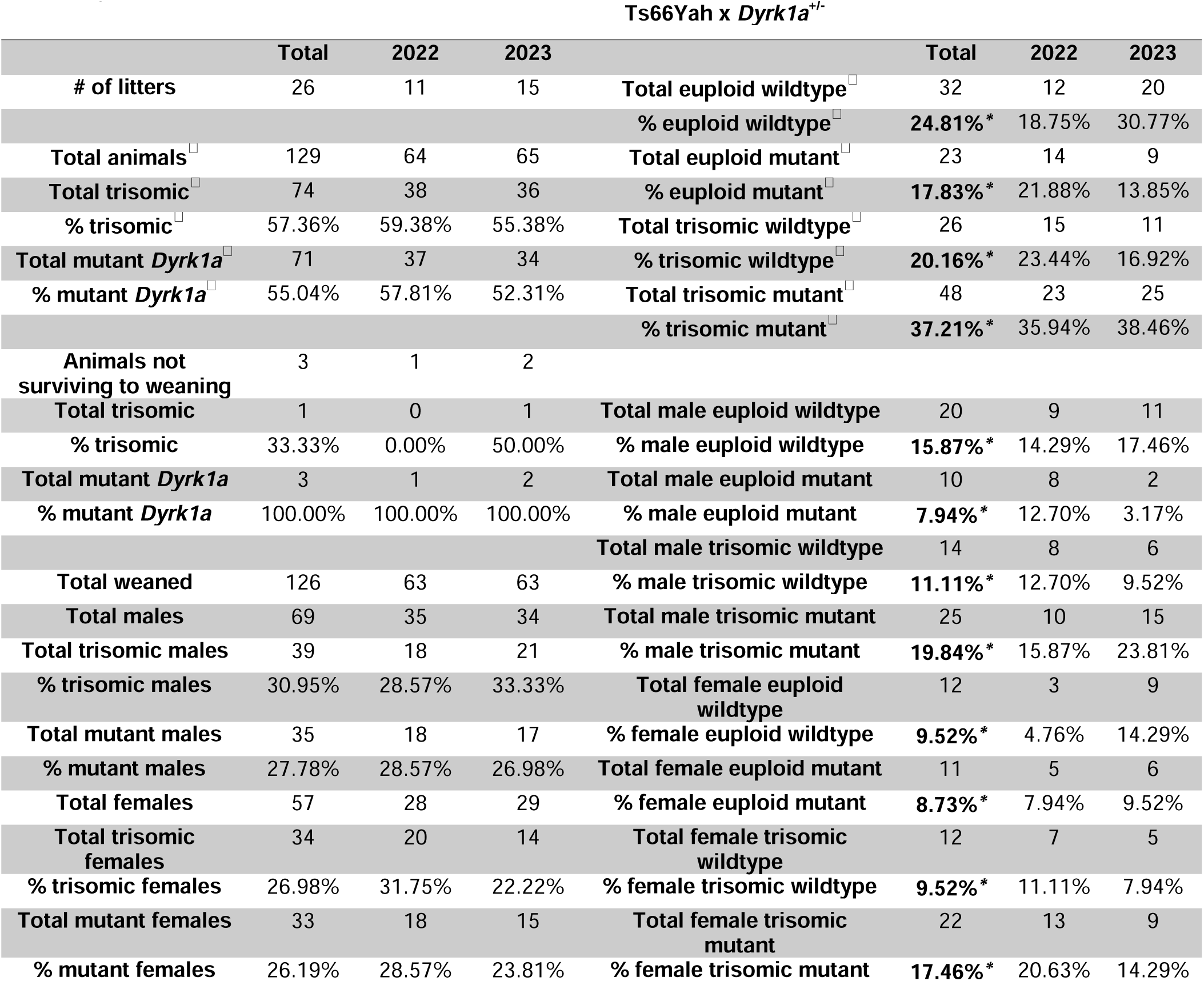
Transmission of trisomy in Ts66Yah x *Dyrk1a*^+/-^ breeding scheme. (^□^) to represent that animals that did not survive to weaning were included. **\* indicates *p* < 0.05** based on chi-square goodness of fit test performed on total data.

**Supplemental Table 3:**
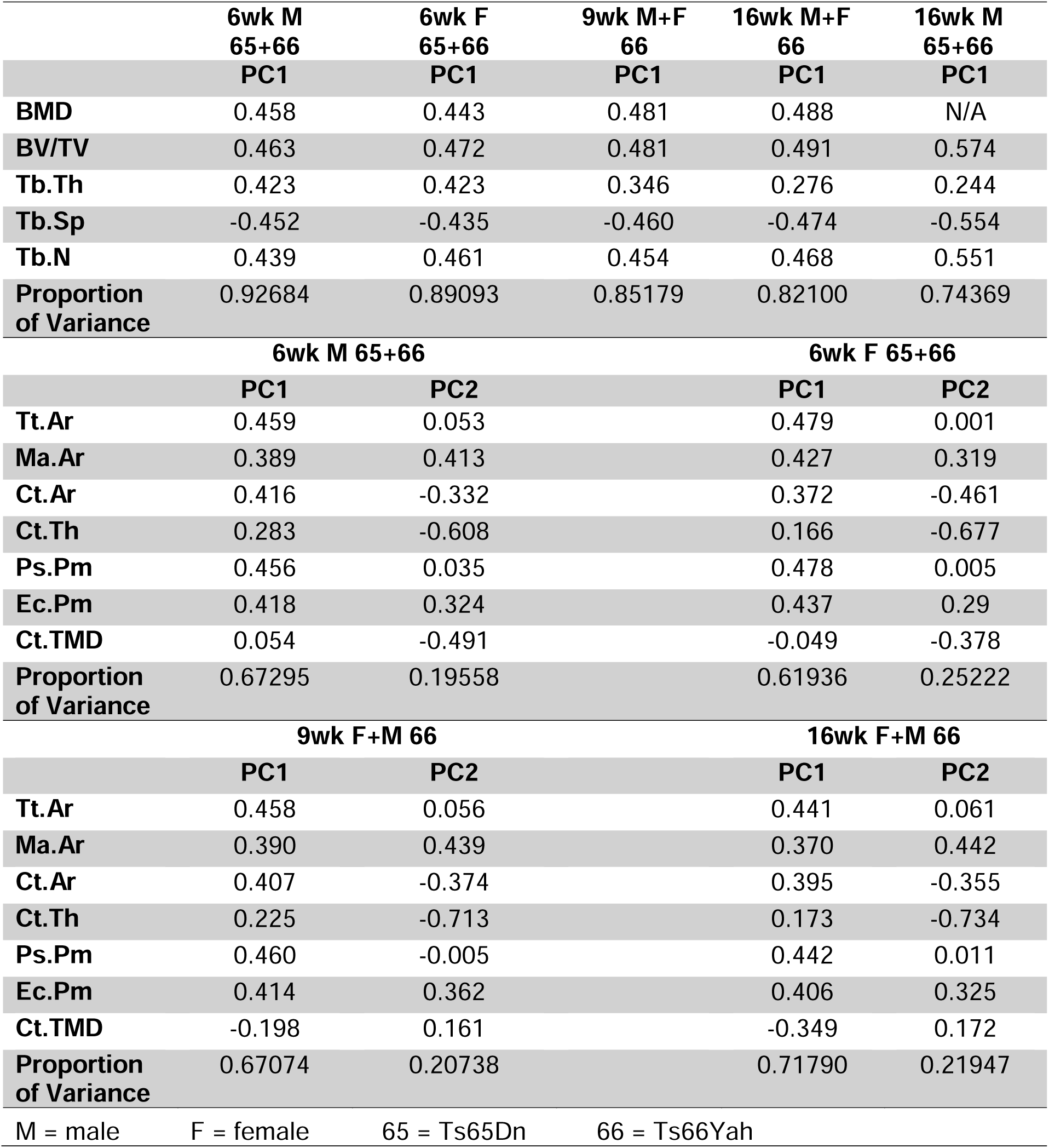
Results of principal components analysis (PCA) for trabecular (top panel) and cortical (bottom two panels) variables. Six-week male and female Ts65Dn data from euploid littermates and Ts65Dn mice lacking OSX-cre (THOMAS *et al*. 2021). Sixteen-week male Ts65Dn data from (BLAZEK *et al*. 2011). All Ts66Yah data comes from this study. N/A indicates data were not comparable, so it was not included in PCA. See Supplemental Figure 2 for PCA biplots.

**Supplemental Table 4:**
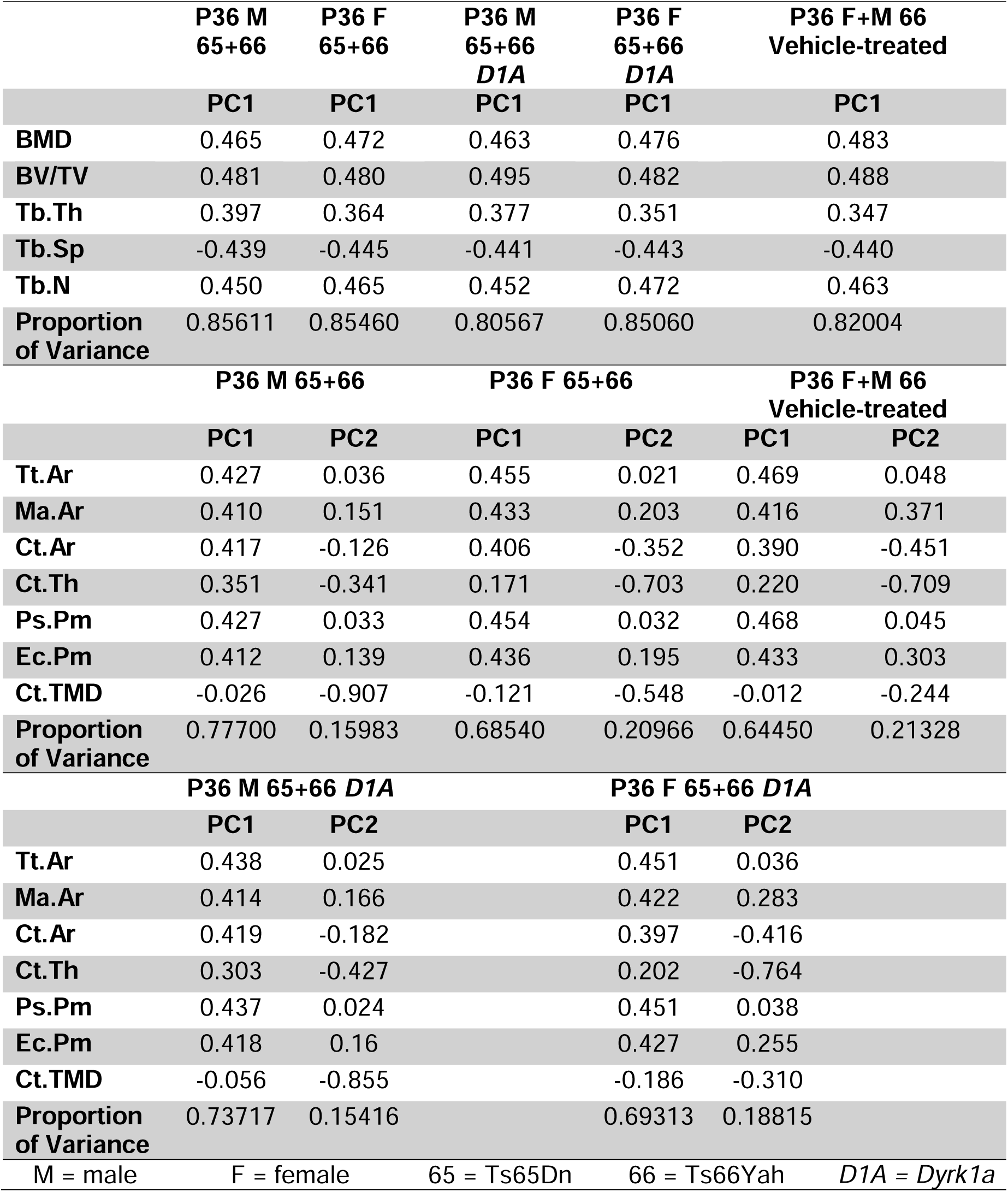
Results of principal components analysis (PCA) for trabecular (top panel) and cortical (bottom panels) variables of P36 Ts65Dn, Ts66Yah, and germline reduction of *Dyrk1a* copy number mice. Ts65Dn data from (LACOMBE *et al*. 2024). Ts66Yah data from this study. Unless otherwise indicated, data comes from euploid and trisomic mice derived from a Ts x *Dyrk1a*^+/-^ breeding scheme without the *Dyrk1a* germline reduction. *Dyrk1a* germline reduction indicates the data included trisomic mice derived from a Ts x *Dyrk1a*^+/-^ breeding scheme with and without a *Dyrk1a* germline reduction. Vehicle-treated indicates the data included mice derived from a Ts66Yah x B6C3F1 breeding scheme given 0.5% carboxymethylcellulose from P21 until P35. See Supplemental Figure 7 for PCA biplots.

