## Supplemental Tables and Figures for "Genetic analysis of triplicated genes affecting sex-specific skeletal deficits in Down syndrome model mice"

Supplemental Table 1: **Transmission of trisomy in Ts66Yah x B6C3F1 breeding scheme**. (**^ǂ^**) to represent that animals that did not survive to weaning were included. * indicates p < 0.05 based on chi-square goodness of fit test performed on total data.

|  | Ts66Yah x B6C3F1 | | | | |
| --- | --- | --- | --- | --- | --- |
|  | **Total** | **2021** | **2022** | **2023** | **2024** |
| # of litters | 114 | 22 | 52 | 28 | 11 |
| Total animals^ǂ^ | 559 | 113 | 255 | 129 | 62 |
| Total trisomic^ǂ^ | 248 | 49 | 118 | 52 | 29 |
| % trisomic^ǂ^ | 44.36%*** | 43.36% | 46.27% | 40.31% | 46.77% |
| Animals not surviving to weaning | 19 | 1 | 8 | 8 | 2 |
| Total trisomic | 8 | 1 | 2 | 3 | 2 |
| % trisomic | 42.11% | 100.00% | 25.00% | 37.50% | 100.00% |
| Total weaned | 540 | 112 | 247 | 121 | 60 |
| Total males | 283 | 58 | 128 | 70 | 27 |
| Total trisomic males | 124 | 26 | 57 | 30 | 11 |
| % trisomic males | 43.82%* | 44.83% | 44.53% | 42.86% | 40.74% |
| Total females | 257 | 54 | 119 | 51 | 33 |
| Total trisomic females | 116 | 22 | 59 | 19 | 16 |
| % trisomic females | 45.14% | 40.74% | 49.58% | 37.25% | 48.48% |

Supplemental Table 2: **Transmission of trisomy in Ts66Yah x Dyrk1a^+/-^ breeding scheme**. (**^ǂ^**) to represent that animals that did not survive to weaning were included. * indicates p < 0.05 based on chi-square goodness of fit test performed on total data.

|  | Ts66Yah x *Dyrk1a*^+/-^ | | | | | | |
| --- | --- | --- | --- | --- | --- | --- | --- |
|  | **Total** | **2022** | **2023** |  | **Total** | **2022** | **2023** |
| # of litters | 26 | 11 | 15 | **Total euploid wildtype^ǂ^** | 32 | 12 | 20 |
|  |  |  |  | **% euploid wildtype^ǂ^** | 24.81%*** | 18.75% | 30.77% |
| Total animals^ǂ^ | 129 | 64 | 65 | **Total euploid mutant^ǂ^** | 23 | 14 | 9 |
| Total trisomic^ǂ^ | 74 | 38 | 36 | **% euploid mutant^ǂ^** | 17.83%*** | 21.88% | 13.85% |
| % trisomic^ǂ^ | 57.36% | 59.38% | 55.38% | **Total trisomic wildtype^ǂ^** | 26 | 15 | 11 |
| Total mutant *Dyrk1a*^ǂ^ | 71 | 37 | 34 | **% trisomic wildtype^ǂ^** | 20.16%*** | 23.44% | 16.92% |
| % mutant *Dyrk1a*^ǂ^ | 55.04% | 57.81% | 52.31% | **Total trisomic mutant^ǂ^** | 48 | 23 | 25 |
|  |  |  |  | **% trisomic mutant^ǂ^** | 37.21%*** | 35.94% | 38.46% |
| Animals not surviving to weaning | 3 | 1 | 2 |  |  |  |  |
| Total trisomic | 1 | 0 | 1 | **Total male euploid wildtype** | 20 | 9 | 11 |
| % trisomic | 33.33% | 0.00% | 50.00% | **% male euploid wildtype** | 15.87%*** | 14.29% | 17.46% |
| Total mutant *Dyrk1a* | 3 | 1 | 2 | **Total male euploid mutant** | 10 | 8 | 2 |
| % mutant *Dyrk1a* | 100.00% | 100.00% | 100.00% | **% male euploid mutant** | 7.94%*** | 12.70% | 3.17% |
|  |  |  |  | **Total male trisomic wildtype** | 14 | 8 | 6 |
| Total weaned | 126 | 63 | 63 | **% male trisomic wildtype** | 11.11%*** | 12.70% | 9.52% |
| Total males | 69 | 35 | 34 | **Total male trisomic mutant** | 25 | 10 | 15 |
| Total trisomic males | 39 | 18 | 21 | **% male trisomic mutant** | 19.84%*** | 15.87% | 23.81% |
| % trisomic males | 30.95% | 28.57% | 33.33% | **Total female euploid wildtype** | 12 | 3 | 9 |
| Total mutant males | 35 | 18 | 17 | **% female euploid wildtype** | 9.52%*** | 4.76% | 14.29% |
| % mutant males | 27.78% | 28.57% | 26.98% | **Total female euploid mutant** | 11 | 5 | 6 |
| Total females | 57 | 28 | 29 | **% female euploid mutant** | 8.73%*** | 7.94% | 9.52% |
| Total trisomic females | 34 | 20 | 14 | **Total female trisomic wildtype** | 12 | 7 | 5 |
| % trisomic females | 26.98% | 31.75% | 22.22% | **% female trisomic wildtype** | 9.52%*** | 11.11% | 7.94% |
| Total mutant females | 33 | 18 | 15 | **Total female trisomic mutant** | 22 | 13 | 9 |
| % mutant females | 26.19% | 28.57% | 23.81% | **% female trisomic mutant** | 17.46%*** | 20.63% | 14.29% |

Supplemental Table 3: **Results of principal components analysis (PCA) for trabecular (top panel) and cortical (bottom two panels) variables.** Six-week male and female Ts65Dn data from euploid littermates and Ts65Dn mice lacking OSX-cre (Thomas *et al.* 2021). Sixteen-week male Ts65Dn data from (Blazek *et al.* 2011). All Ts66Yah data comes from this study. N/A indicates data were not comparable, so it was not included in PCA. See Supplemental Figure 2 for PCA biplots.

|  | **6wk M 65+66** | **6wk F**  **65+66** | | **9wk M+F**  **66** | | **16wk M+F**  **66** | | | **16wk M 65+66** | |
| --- | --- | --- | --- | --- | --- | --- | --- | --- | --- | --- |
|  | **PC1** | **PC1** | | **PC1** | | | | **PC1** | | **PC1** |
| **BMD** | 0.458 | 0.443 | | 0.481 | | | | 0.488 | | N/A |
| **BV/TV** | 0.463 | 0.472 | | 0.481 | | | | 0.491 | | 0.574 |
| **Tb.Th** | 0.423 | 0.423 | | 0.346 | | | | 0.276 | | 0.244 |
| **Tb.Sp** | -0.452 | -0.435 | | -0.460 | | | | -0.474 | | -0.554 |
| **Tb.N** | 0.439 | 0.461 | | 0.454 | | | | 0.468 | | 0.551 |
| **Proportion of Variance** | 0.92684 | 0.89093 | | 0.85179 | | | | 0.82100 | | 0.74369 |
|  | **6wk M 65+66** | |  | | **6wk F 65+66** | | | | | |
|  | **PC1** | **PC2** | |  | | | | **PC1** | | **PC2** |
| **Tt.Ar** | 0.459 | 0.053 | |  | | | | 0.479 | | 0.001 |
| **Ma.Ar** | 0.389 | 0.413 | |  | | | | 0.427 | | 0.319 |
| **Ct.Ar** | 0.416 | -0.332 | |  | | | | 0.372 | | -0.461 |
| **Ct.Th** | 0.283 | -0.608 | |  | | | | 0.166 | | -0.677 |
| **Ps.Pm** | 0.456 | 0.035 | |  | | | | 0.478 | | 0.005 |
| **Ec.Pm** | 0.418 | 0.324 | |  | | | | 0.437 | | 0.29 |
| **Ct.TMD** | 0.054 | -0.491 | |  | | | | -0.049 | | -0.378 |
| **Proportion of Variance** | 0.67295 | 0.19558 | |  | | | | 0.61936 | | 0.25222 |
|  | **9wk F+M 66** | | |  | | | **16wk F+M 66** | | | |
|  | **PC1** | **PC2** | |  | | | | **PC1** | | **PC2** |
| **Tt.Ar** | 0.458 | 0.056 | |  | | | | 0.441 | | 0.061 |
| **Ma.Ar** | 0.390 | 0.439 | |  | | | | 0.370 | | 0.442 |
| **Ct.Ar** | 0.407 | -0.374 | |  | | | | 0.395 | | -0.355 |
| **Ct.Th** | 0.225 | -0.713 | |  | | | | 0.173 | | -0.734 |
| **Ps.Pm** | 0.460 | -0.005 | |  | | | | 0.442 | | 0.011 |
| **Ec.Pm** | 0.414 | 0.362 | |  | | | | 0.406 | | 0.325 |
| **Ct.TMD** | -0.198 | 0.161 | |  | | | | -0.349 | | 0.172 |
| **Proportion of Variance** | 0.67074 | 0.20738 | |  | | | | 0.71790 | | 0.21947 |
| M = male | F = female | 65 = Ts65Dn | | 66 = Ts66Yah | | | | | |  |

Supplemental Table 4: **Results of principal components analysis (PCA) for trabecular (top panel) and cortical (bottom panels) variables of P36 Ts65Dn, Ts66Yah, and germline reduction of Dyrk1a copy number mice.** Ts65Dn data from (LaCombe et al. 2024). Ts66Yah data from this study. Unless otherwise indicated, data comes from euploid and trisomic mice derived from a Ts x Dyrk1a^+/-^ breeding scheme without the Dyrk1a germline reduction. Dyrk1a germline reduction indicates the data included trisomic mice derived from a Ts x Dyrk1a^+/-^ breeding scheme with and without a Dyrk1a germline reduction. Vehicle-treated indicates the data included mice derived from a Ts66Yah x B6C3F1 breeding scheme given 0.5% carboxymethylcellulose from P21 until P35. See Supplemental Figure 7 for PCA biplots.

|  | **P36 M**  **65+66** | **P36 F**  **65+66** | **P36 M**  **65+66**  ***D1A*** | **P36 F**  **65+66**  ***D1A*** | | **P36 F+M 66**  **Vehicle-treated** | |
| --- | --- | --- | --- | --- | --- | --- | --- |
|  | **PC1** | **PC1** | **PC1** | **PC1** | | **PC1** | |
| **BMD** | 0.465 | 0.472 | 0.463 | 0.476 | | 0.483 | |
| **BV/TV** | 0.481 | 0.480 | 0.495 | 0.482 | | 0.488 | |
| **Tb.Th** | 0.397 | 0.364 | 0.377 | 0.351 | | 0.347 | |
| **Tb.Sp** | -0.439 | -0.445 | -0.441 | -0.443 | | -0.440 | |
| **Tb.N** | 0.450 | 0.465 | 0.452 | 0.472 | | 0.463 | |
| **Proportion of Variance** | 0.85611 | 0.85460 | 0.80567 | 0.85060 | | 0.82004 | |
|  | **P36 M 65+66** | | **P36 F 65+66** | | | **P36 F+M 66**  **Vehicle-treated** | |
|  | **PC1** | **PC2** | **PC1** | **PC2** | | **PC1** | **PC2** |
| **Tt.Ar** | 0.427 | 0.036 | 0.455 | 0.021 | | 0.469 | 0.048 |
| **Ma.Ar** | 0.410 | 0.151 | 0.433 | 0.203 | | 0.416 | 0.371 |
| **Ct.Ar** | 0.417 | -0.126 | 0.406 | -0.352 | | 0.390 | -0.451 |
| **Ct.Th** | 0.351 | -0.341 | 0.171 | -0.703 | | 0.220 | -0.709 |
| **Ps.Pm** | 0.427 | 0.033 | 0.454 | 0.032 | | 0.468 | 0.045 |
| **Ec.Pm** | 0.412 | 0.139 | 0.436 | 0.195 | | 0.433 | 0.303 |
| **Ct.TMD** | -0.026 | -0.907 | -0.121 | -0.548 | | -0.012 | -0.244 |
| **Proportion of Variance** | 0.77700 | 0.15983 | 0.68540 | 0.20966 | | 0.64450 | 0.21328 |
|  | **P36 M 65+66 *D1A*** | |  | | **P36 F 65+66 *D1A*** | |  |
|  | **PC1** | **PC2** |  | **PC1** | | **PC2** |  |
| **Tt.Ar** | 0.438 | 0.025 |  | 0.451 | | 0.036 |  |
| **Ma.Ar** | 0.414 | 0.166 |  | 0.422 | | 0.283 |  |
| **Ct.Ar** | 0.419 | -0.182 |  | 0.397 | | -0.416 |  |
| **Ct.Th** | 0.303 | -0.427 |  | 0.202 | | -0.764 |  |
| **Ps.Pm** | 0.437 | 0.024 |  | 0.451 | | 0.038 |  |
| **Ec.Pm** | 0.418 | 0.16 |  | 0.427 | | 0.255 |  |
| **Ct.TMD** | -0.056 | -0.855 |  | -0.186 | | -0.310 |  |
| **Proportion of Variance** | 0.73717 | 0.15416 |  | 0.69313 | | 0.18815 |  |
| M = male | F = female | | 65 = Ts65Dn | 66 = Ts66Yah | | | *D1A = Dyrk1a* |

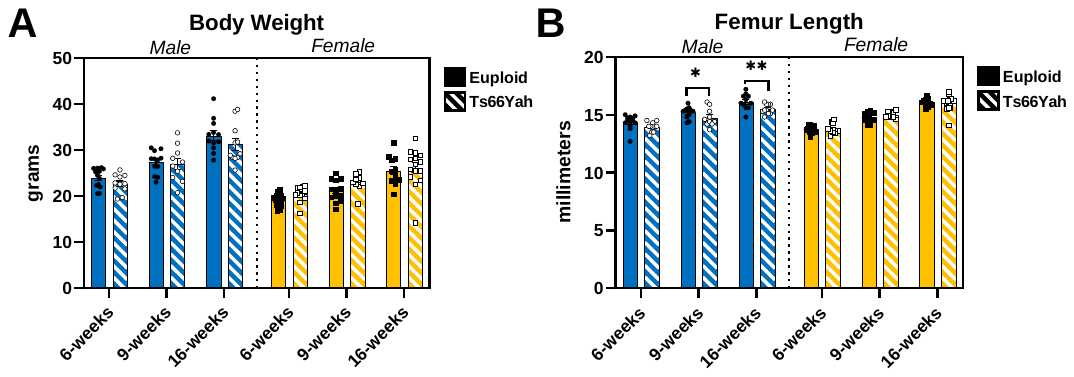

Supplemental Figure 1: **Body weight and femur lengths of 6-, 9-, and 16-week-old Ts66Yah mice.** Data are mean ± SEM. Asterisks indicate a significant difference between groups in pairwise comparisons with Sidak correction. * p < 0.05, ** p < 0.01. **A)** No significant differences were found in any pairwise comparisons for body weight. **B)** Pairwise comparisons between ages within genotype and sex for femur length: significantly increased between each age in all four groups. 6 weeks: male euploid (n = 13), male Ts66Yah (n = 11), female euploid (n = 15 [body weight] or 13 [femur length]), female Ts66Yah (n = 12 [body weight] or 11 [femur length]); 9 weeks: male euploid (n = 12), male Ts66Yah (n = 10), female euploid (n = 13), female Ts66Yah (n = 9); 16 weeks: male euploid (n = 12 [body weight] or 11 [femur length]), male Ts66Yah (n = 11), female euploid (n = 12), female Ts66Yah (n = 15).

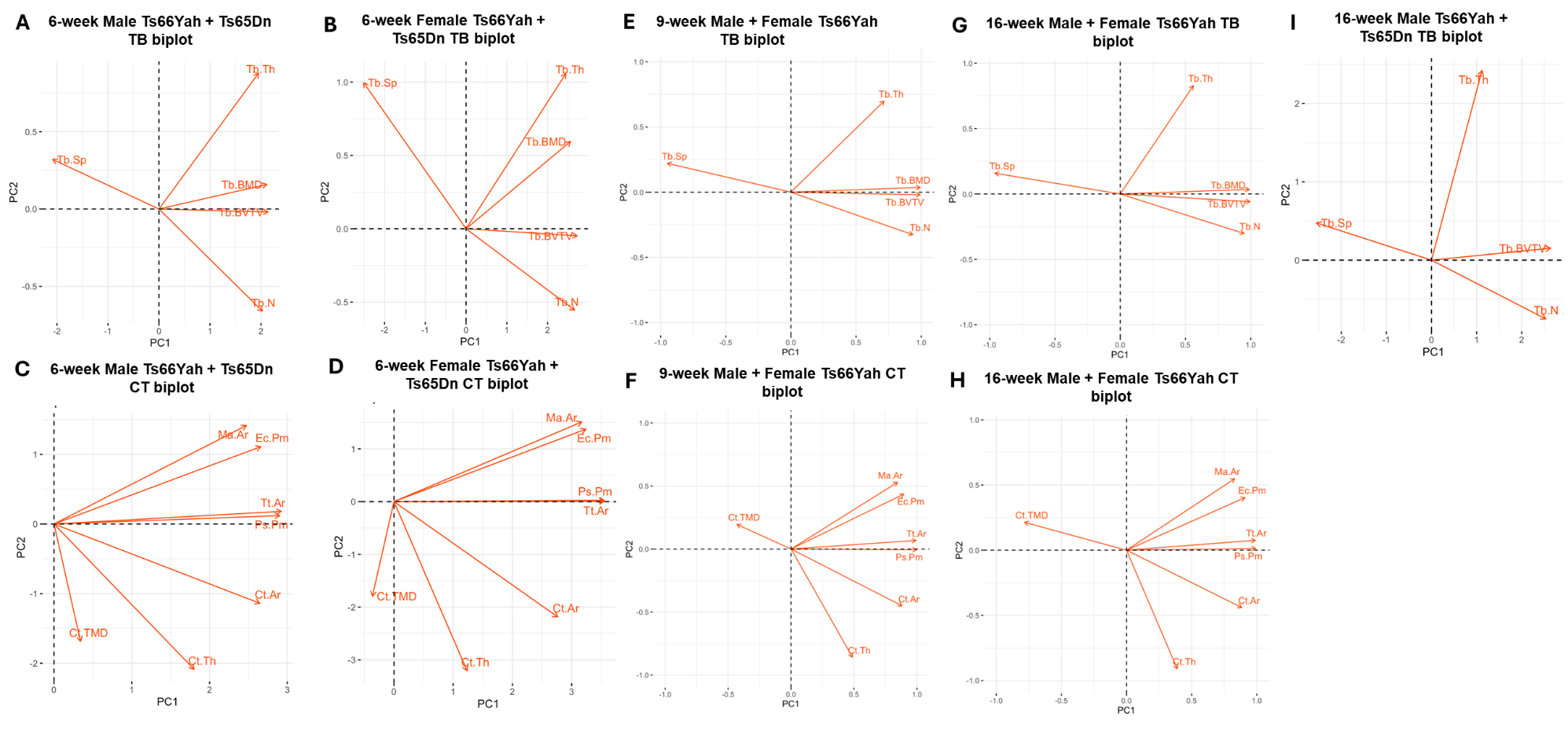

Supplemental Figure 2: **Principal component analysis biplots for trabecular (TB) and cortical (CT) variables of Ts65Dn and Ts66Yah mice. A-D)** Biplots generated separately for male (A,C) and female (B,D) trabecular (A,B) and cortical (C,D) variables using 6-week male and female Ts65Dn data from euploid littermates and Ts65Dn mice lacking OSX-cre (Thomas *et al.* 2021) and 6-week male and female Ts66Yah data (this study). **E-F)** Biplots generated separately for trabecular (E) and cortical (F) variables using 9-week male and female Ts66Yah data (this study). **G-H)** Biplots generated separately for trabecular (G) and cortical (H) variables using 16-week male and female Ts66Yah data (this study. **I)** Biplot generated for male trabecular variables using 16-week male Ts65Dn data (Blazek *et al.* 2011) and 16-week male Ts66Yah data (this study). See Supplemental Table 3 for PCA results.

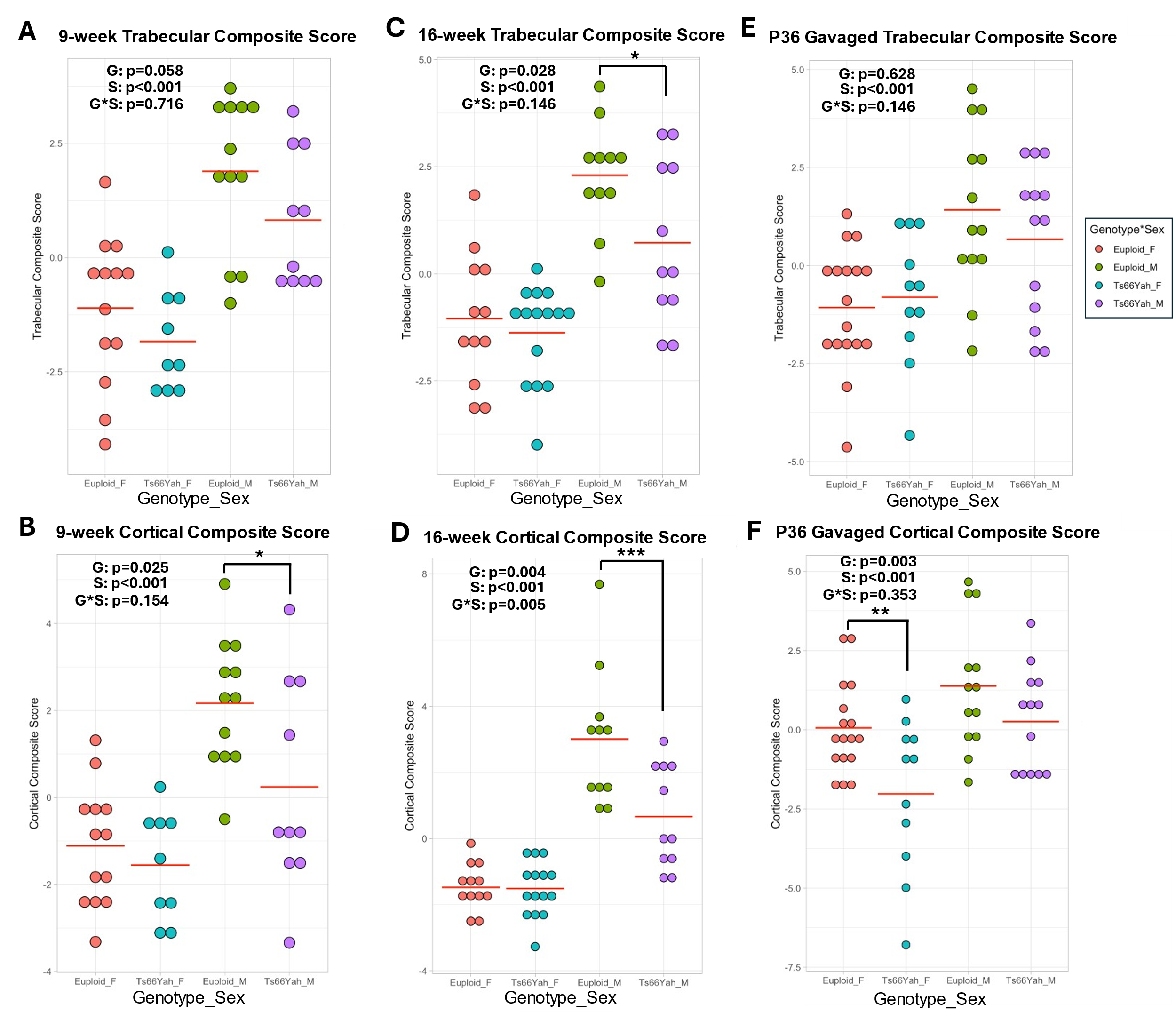

Supplemental Figure 3: **Trabecular and cortical composite scores for male and female Ts66Yah at 9 weeks (A-B), 16 weeks (C-D), and P36 vehicle-treated mice (E-F).** Red horizontal line indicates group mean. Two-way ANOVA with genotype (G) and sex (S) as between subject factors. Asterisks indicate significant genotype difference in sex-stratified contrast analysis. **p* < 0.05, ** *p* < 0.01, *** *p* < 0.001*.*

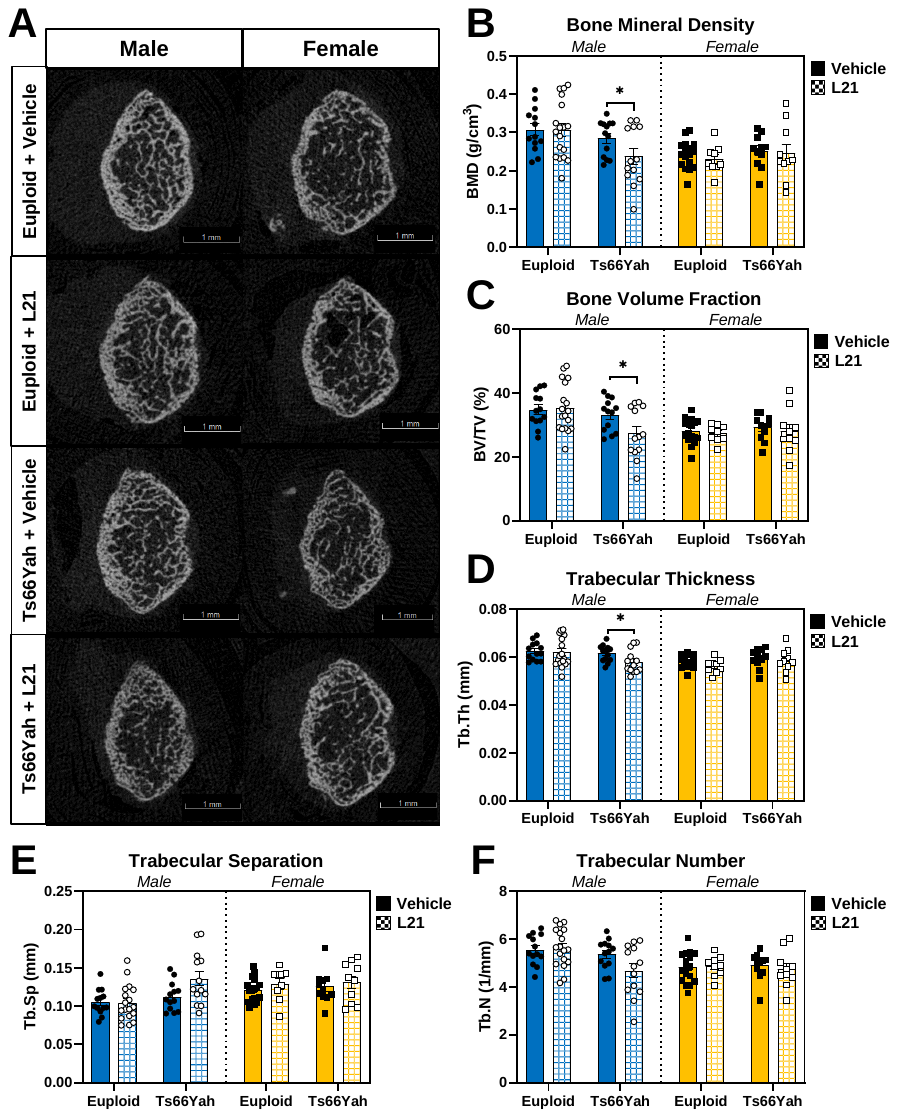

Supplemental Figure 4: **Trabecular bone variables for Leucettinib-21-treated Ts66Yah mice. A)** Representative images of trabecular bone taken halfway way through the 1mm trabecular region as determined by finding the animal with the closest average distance away from the mean of each trabecular variable. **B-F)** Data are mean ± SEM. Asterisks indicate a significant difference between groups in pairwise comparisons with Sidak correction. * p < 0.05. Male mice: vehicle-treated euploid (n = 13), L21-treated euploid (n = 18), vehicle-treated Ts66Yah (n = 13), L21-treated Ts66Yah (n = 13). Female mice: vehicle-treated euploid (n = 17), L21-treated euploid (n = 9), vehicle-treated Ts66Yah (n = 11), L21-treated Ts66Yah (n = 10).

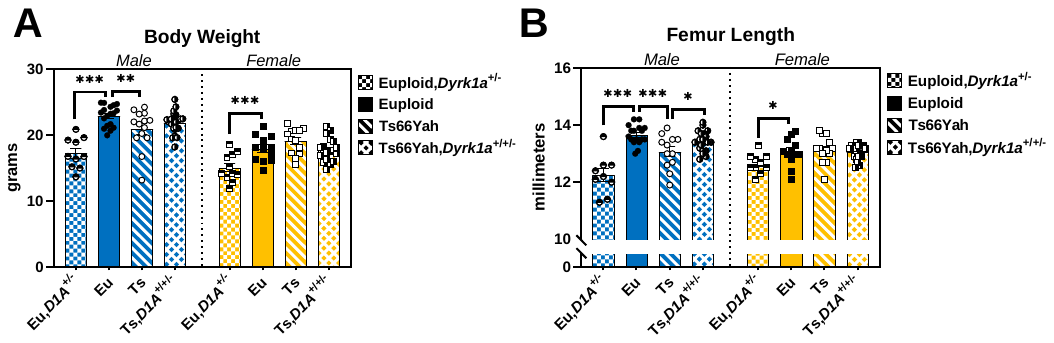

Supplemental Figure 5: **Body weight and femur lengths of postnatal day (P)36 Ts66Yah,Dyrk1a^+/+/-^ mice.** Data are mean ± SEM. Asterisks indicate a significant difference between groups in pairwise comparisons with Sidak correction. * p < 0.05, ** p < 0.01, *** p < 0.001. Male mice: euploid (n = 18 [body weight] or 17 [femur length]); euploid,Dyrk1a^+/-^ (n = 10 [body weight] or 9 [femur length]); Ts66Yah (n = 14 [body weight] or 12 [femur length]); Ts66Yah,Dyrk1a^+/+/-^ (n = 22 [body weight] or 21 [femur length]). Female mice: euploid (n = 12); euploid,Dyrk1a^+/-^ (n = 11 [body weight] or 10 [femur length]); Ts66Yah (n = 13); Ts66ah,Dyrk1a^+/+/-^ (n = 19).

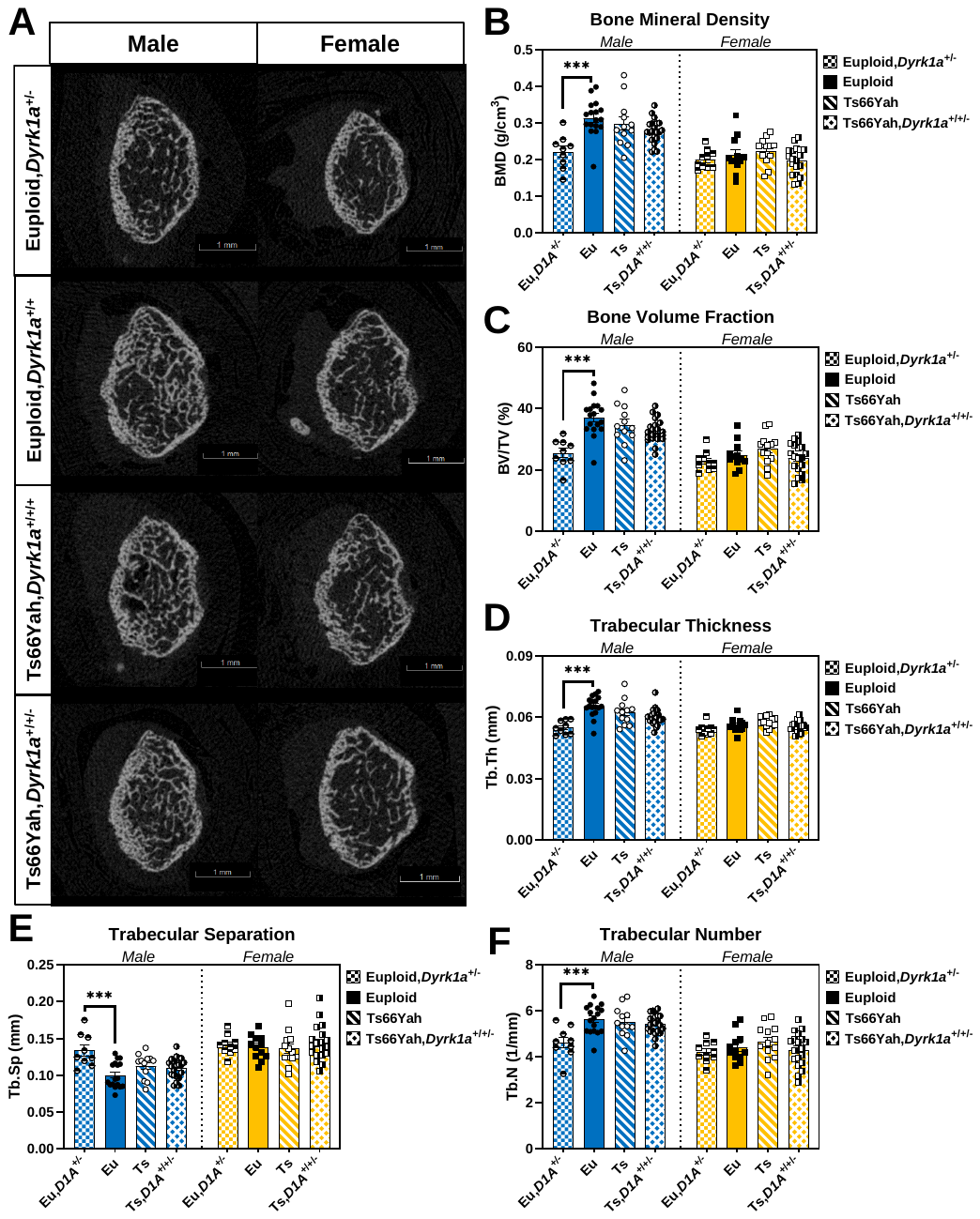

Supplemental Figure 6: **Trabecular bone variables for P36 Ts66Yah,*Dyrk1a^+/+/-^* mice.** **A)** Representative images of trabecular bone taken halfway through the 1mm trabecular region as determined by finding the animal with the closest average distance away from the mean of each trabecular variable. **B-F)** Data are mean ± SEM. Asterisks indicate a significant difference between groups in pairwise comparisons with Sidak correction. *** *p* < 0.001. Male mice: euploid (n = 17); euploid,Dyrk1a^+/-^ (n = 9); Ts66Yah (n = 12); Ts66Yah,Dyrk1a^+/+/-^ (n = 21). Female mice: euploid (n = 12); euploid,Dyrk1a^+/-^ (n = 10); Ts66Yah (n = 13); Ts66ah,Dyrk1a^+/+/-^ (n = 19).

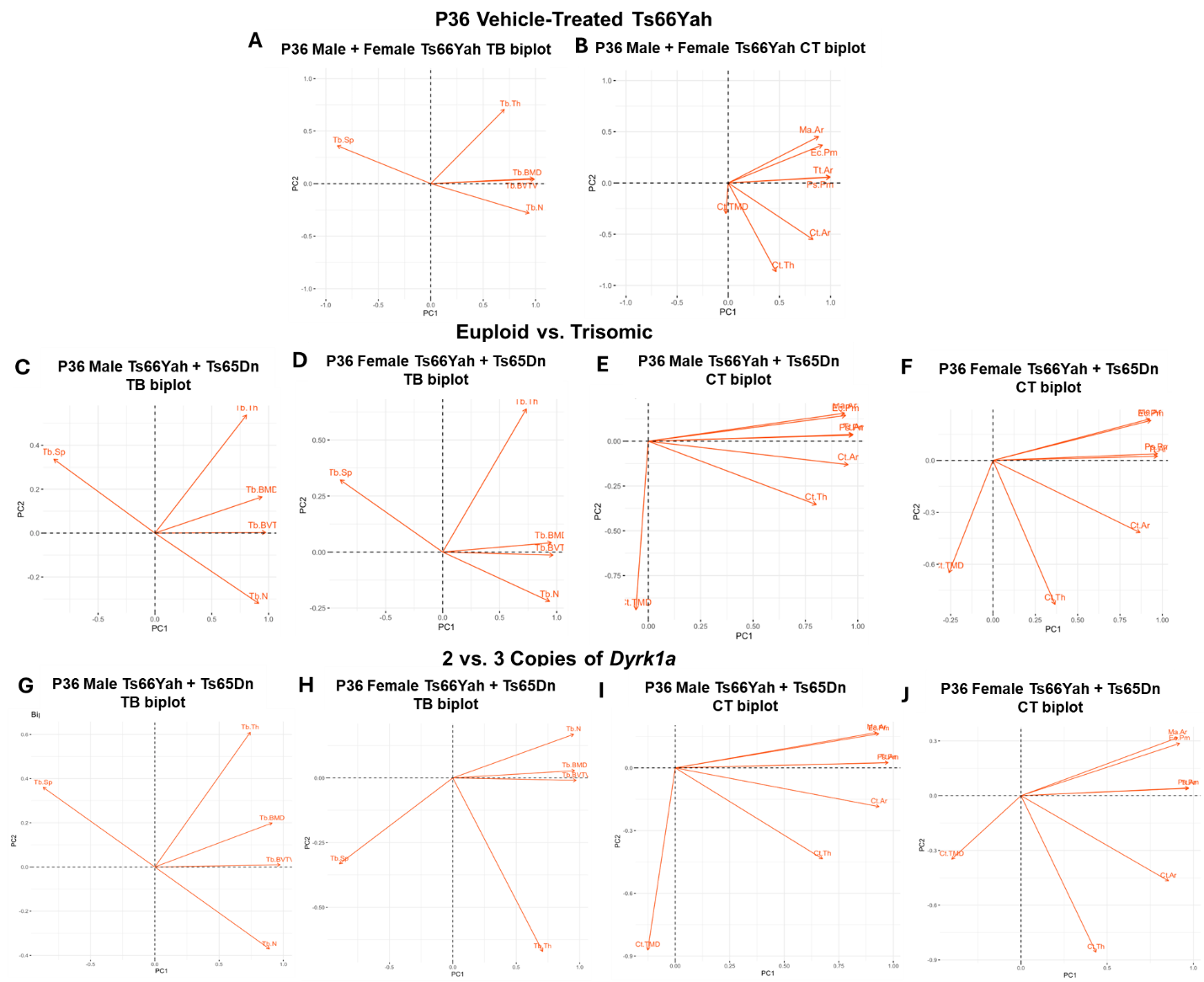

Supplemental Figure 7: **Principal component analysis biplots for trabecular (TB) and cortical (CT) variables of Ts65Dn, Ts66Yah, and germline reduction of *Dyrk1a* copy number mice. A-B)** Biplots generated separately for trabecular (A) and cortical (B) variables using postnatal day (P)36 male and female vehicle-treated Ts66Yah mice (this study). **C-F)** Biplots generated separately for male (C,E) and female (D,F) trabecular (C,D) and cortical (E,F) variables using P36 Ts65Dn,*Dyrk1a^+/+/+^* and euploid,*Dyrk1a^+/+^* data from (LaCombe *et al.* 2024) and P36 Ts66Yah,*Dyrk1a*^+/+/+^ and euploid,*Dyrk1a*^+/+^ data from this study. **G-J)** Biplots generated separately for male (G,I) and female (H,J) trabecular (G,H) and cortical (I,J) variables using P36 Ts65Dn,*Dyrk1a^+/+/+^* and Ts65Dn,*Dyrk1a*^+/+/-^ data from (LaCombe *et al.* 2024) and P36 Ts66Yah,*Dyrk1a*^+/+/+^ and Ts66Yah,*Dyrk1a*^+/+/-^ data from this study. See Supplemental Table 4 for PCA results.
